# Grapevine Rootstock and Scion Genotypes’ Symbiosis with Soil Microbiome: A Machine Learning Revelation for Climate-Resilient Viticulture

**DOI:** 10.1101/2024.02.25.581926

**Authors:** Lakshay Anand, Thanos Gentimis, Allan Bruce Downie, Carlos M. Rodriguez Lopez

## Abstract

Given the impact of climate change on agriculture, the development of resilient crop cultivars is imperative. A healthy plant microbiota is key to plant productivity, influencing nutrient absorption, disease resistance, and overall vigor. The plant genetic factors controlling the assembly of microbial communities are still unknown. Here we examine if Machine Learning can predict grapevine rootstock and scion genotypes based on soil microbiota, despite environmental variability. The study utilized soil microbial bacteriome datasets from 281 vineyards across 13 countries and five continents, featuring 34 different *Vitis vinifera* cultivars grafted onto, often ambiguous, rootstocks. Random Forests, Adaptive Boost, Gradient Boost, Support Vector Machines, Gaussian and Bernoulli Naïve Bayes, k-Nearest Neighbor, and Neural Networks algorithms were employed to predict continent, country, scion, and rootstock cultivar, under two filtering criteria: retaining sparse classes, ensuring class diversity, and excluding sparse classes assessing model robustness against overfitting. Both criteria showed remarkable F1-weighted scores (>0.8) for all classes, for most algorithms. Moreover, successful rootstock and scion genotype prediction from soil microbiomes confirms that genotypes of both plant parts shape the microbiome. These insights pave the way for identifying plant genes for use with breeding programs that enhance plant productivity and sustainability by improving the plant-microbiota relationship.

## Introduction

Wine, a symbol of heritage and culture, represents an industry with a global annual market value of USD 489.3 Billion in 2021, and it is predicted to almost double by 2030 (Acumen Research and Consulting, 2022). However, the wine industry, like many other agricultural sectors, is not immune to the adverse impacts of climate change. Projected environmental changes, such as a 2°C increase in average temperatures by the end of the century, have the potential to usher in more frequent extreme weather events, including reduced rainfall and soaring temperatures (Hall et al., 2016; Kang & Banga, 2013; Neethling et al., 2017). This could render up to 81% of US vineyards unsuitable for wine production by 2040 (Diffenbaugh et al., 2011; White et al., 2006). To navigate these challenges, there’s a dire need for growers to maintain grape quality while minimizing resource inputs, emphasizing the need for breeding environmentally resilient grapevine cultivars.

Traditionally, the targets of crop breeding programs have been the plant species genes controlling a given trait. However, it is vital to recognize that plants, including grapevines (*Vitis vinifera*), are not solitary entities. They coexist with a myriad of microorganisms collectively referred to as the plant microbiota in, on, and surrounding the plant. This intricate interplay between the host plant and its microbiota forms the ‘holobiont.’ The rhizosphere (soil around the roots), phyllosphere (leaf surfaces), and carposphere (grape surface) are major habitats hosting microbial populations that contribute to plant-microbe interactions (Bulgarelli et al., 2013; Jones et al., 2019; Mendes et al., 2013; Shi et al., 2015; Vorholt, 2012). The microbiota offers numerous advantages to the plant host, such as improved water and nutrient uptake, enhanced growth rates, and fortified defenses against biotic- and abiotic-stresses (Bettenfeld et al., 2022; Darriaut et al., 2023). The growing environment (climate and soil type), which is defined by the vineyard’s geographical location (Fabres et., 2017), has the most significant impact on the composition of the vine’s microbiota (Burns et al., 2015; Zhou et al., 2021). Other factors affecting the vine microbiota composition are the planted cultivar (both scion and rootstock) (Marasco et al., 2022) and the vineyard management (Zhou et al., 2021).

Several studies have shown that crop domestication unintentionally alters the microbial communities of target crops (Chaluvadi & Bennetzen, 2018; Germida & Siciliano, 2001; Leff et al., 2017; Szoboszlay et al., 2015), suggesting that plant-associated, soil microbiota composition can be bred as a trait (Wissuwa et al., 2009). There has been a lack of research on the study of host-microbe interactions from a community viewpoint, which is a critical aspect in the development of breeding programs aimed at improving productivity, quality, and sustainability through the enhancement of the holobiont. Understanding what constitutes a “good microbiota host” is of utmost importance in this regard.

Next-generation sequencing (NGS) approaches such as metabarcoding, metagenomics, and metatranscriptomics facilitate the collection of genetic information from plant microbiomes (Caporaso et al., 2012). These methods allow researchers to characterize microbial diversity, determine the presence and abundance of specific microorganisms, and understand their functional roles within the grapevine ecosystem. (Caporaso et al., 2012) Metabarcoding involves the extraction of genetic material (e.g., DNA) from a complex mixture of microorganisms located, and living in, such diverse substratesas soil, water, or even gut contents, and then amplifying and sequencing a specific genetic marker, often a short DNA region called a “barcode” or “marker gene.” This barcode region is typically highly conserved within a phylogenetically similar taxa but varies with the phylogenetic distance among different taxa, making it useful for rough taxonomic identification. The 16S barcode, specifically the 16S ribosomal RNA (rRNA) gene, is commonly used in metabarcoding and microbial ecology studies to identify and classify bacteria and archaea (Ward et al., 1990).

As we tread into the age of digitization, artificial intelligence (AI) and machine learning (ML) offer powerful tools for deciphering complex biological systems. ML has emerged as a potent tool for deciphering the complex interactions between the plant and it’s microbiomes. Leveraging large-scale datasets generated through sequencing technologies, ML models can potentially facilitate understanding of the complex interactions between host and microbiota (Gobbi et al., 2022). This study aims to harness the predictive capacities of ML algorithms, using them as a lens to discern the intertwined influences of cultivar and environment on soil microbiota composition. With a vast dataset representing soil samples from vineyards across five continents and 13 countries involving 34 grapevine scion cultivars, ML algorithms are utilized to predict the geographic origin (continent and country) and grapevine scion cultivar based on the composition of soil microbiota alone. Occassionally, the rootstock was known, and this was also predicted from the associated microbiome. This triad of Continent, Country, and Cultivar – the “3C’s” – presents a fascinating dimension to understanding the influences on the plant-associated, soil microbiota assembly. This research attempts to test a novel hypothesis: If ML can accurately predict the planted genotype using soil microbiota composition information, notwithstanding environmental influences, it could revolutionarily be deployed to pinpoint the plant genes whose products govern microbiota assembly.

This study provides a stepping-stone for understanding the crop-microbiome complex interaction and opens the door to the use of ML to identify the genes whose protein products control crop microbiome assembly. Moreover, grapevine provides a unique and more than usually complex test case because commercial grapevines are normally the result of the grafting of a scion onto a rootstock that are not of the same variety (nonconvarietal), which results in the interaction of two different genotypes to assemble a single soil microbiome.

### Study overview

We used previously published 885 soil bacteriome datasets collected from 281 vineyards located in 13 countries across 5 continents and planted with 34 Vitis vinifera cultivars on unidentified rootstocks, representing the largest collection of vineyard soil microbiomes analyzed to date. The dataset utilized non-redundant amplicon sequence variants as features, with continent, country, or scion cultivar (rootstock cultivar when available) as class labels. Nine algorithms, Random Forests, Adaptive Boost, Gradient Boost, Support Vector Machines (linear and radial kernel), Gaussian and Bernoulli Naïve Bayes, k-Nearest Neighbor, and Neural Networks(multi-layer perceptron), were applied under two filtering criteria: (i) retaining sparse classes, ensuring class diversity, and (ii) excluding sparse classes to assess model robustness against overfitting (Figure 1 and Methods File).

**Figure 1.**
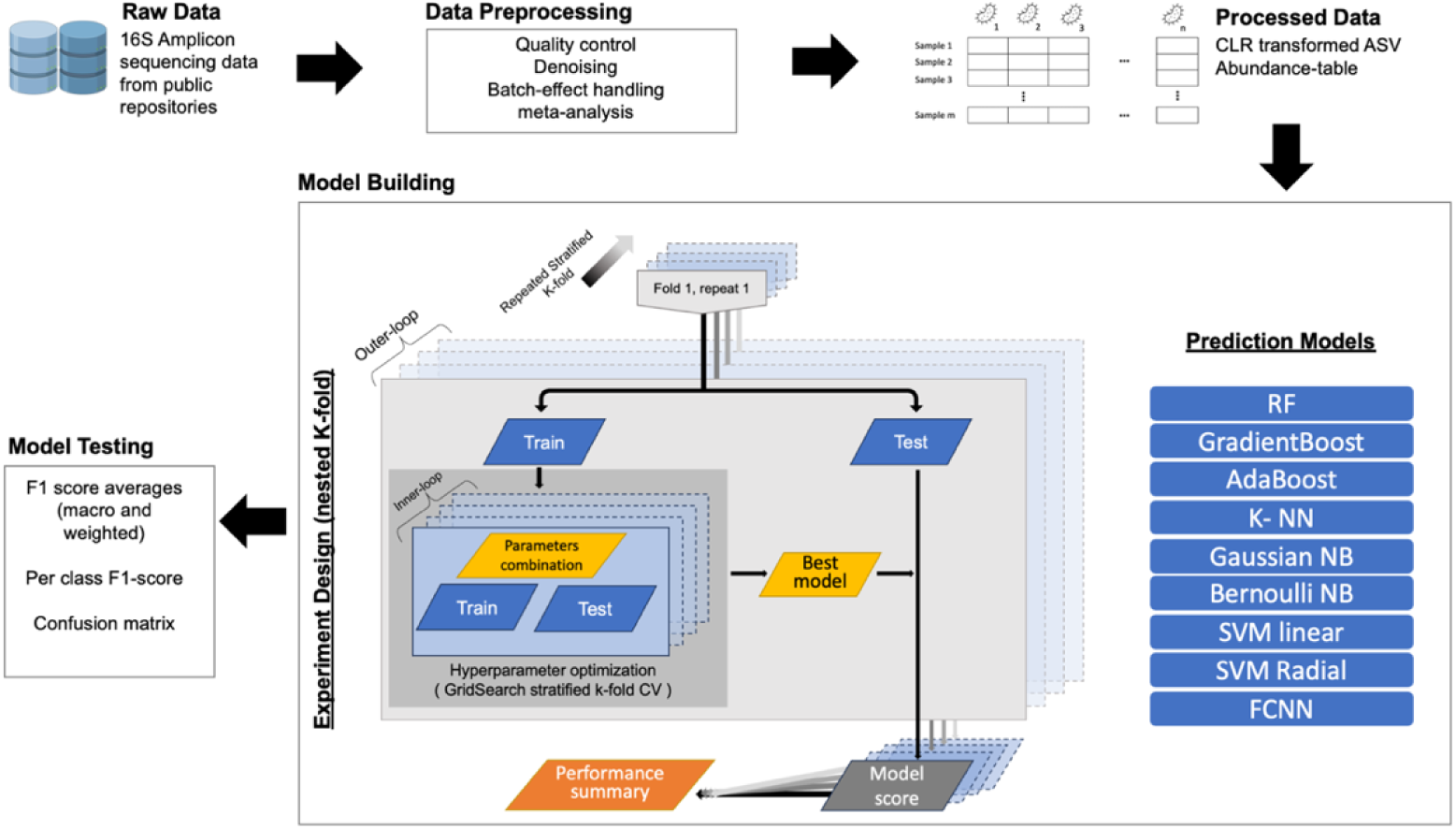
Schematic illustration of the workflow used in this study. Raw data comprising amplicons to the hypervariable region of the 16S ribosomal genes of reported soil microorganisms associated with grapevine were acquired and subjected to quality control (Data processing). Centered log-ratio transformed (CLR) amplicon sequence variants (ASV) comprised the processed data that was randomly partitioned into training data (75%/80% of the samples; inclusion/exclusion of sparse classes) and testing data (25%/20% of the samples; inclusion/exclusion of sparse classes) to assess the performance of 9 different ML algorithms using several model tests.

## Methods

### The Datasets

Previously published amplicon-sequencing datasets, targeting the hypervariable-3 and -4 (V3-V4) regions of the 16S Small SubUnit (SSU) rRNA gene, generated from 885 vineyard soil samples, were used in this study. The datasets were obtained from four sources referred to as I-IV hereafter (Supplementary Table 1). Dataset I consisted of 252 samples collected from 200 vineyards (Gobbi et al., 2022). Dataset II was generated in-house and constitutes 66 samples collected from 22 vineyards in Australia’s Barossa Valley (Zhou et al., 2021). Datasets III and IV, respectively, consisted of 426 and 141 samples obtained from studies (Zarraonaindia et al., 2015) and (Marasco et al., 2022), collected from 5 and 10 vineyards, respectively. Soil samples used to generate these datasets were collected from vineyards located in 13 countries across five continents and planted with 34 *Vitis vinifera* scion cultivars on, in some cases, unspecified rootstocks, representing the largest collection of vineyard microbiomes analyzed to date (See Supplementary Table 2 for the metadata for each sample/dataset).

### Data pre-processing

#### Dataset heterogeneity and batch effects

Although all the datasets used in this study were generated using Illumina short-read sequencing technology, raw sequences were heterogenous with respect to sample size, sequencing type (single-end (SE) versus paired-end (PE) reads), hypervariable region sequenced, and read length (Supplementary Table 1). Batch effect is a common bias caused especially when datasets from different studies are combined and, if not addressed properly, could lead to a risk of misleading results (Henschel et al., 2015). Several studies reported that the risk is comparatively higher when the samples are from similar experimental studies (similar in terms of experiment design, methodology etc.) (Henschel et al., 2015; Lozupone & Knight, 2007; Lozupone et al., 2013). Both heterogeneity and batch effects were expected here as the datasets originated from diverse experimental studies. The rest of this section describes the methods used to address these issues.

Dataset II consisted solely of paired-end results. Datasets I and III consisted of both single-end and paired-end samples, while the publicly available raw data for Dataset IV was already merged and, therefore, treated as single-end reads. Although the pipelines used downstream can handle both single- and paired-end sequences as input, to maintain consistency in the sequencing type among all the datasets, all the paired-end sequences were merged using the paired-end read merger (PEARS) tool (Zhang et al., 2014). The quality of the sequencing data was assessed using Fastqc version 0.11.9, while multiqc version 1.11 (Ewels et al., 2016) was used to visualize the quality control results for the entire dataset. No adapters were identified in any dataset; hence, the adapter-removal step was omitted and the and the raw merged (single-end or PEARS-merged PE) sequences, for each dataset, were used for QIIME2 (v2022.1;(Bolyen et al., 2019)) analysis.

Due to the differences in average read lengths (Supplementary Table 1), each dataset was analyzed separately in QIIME2. The DADA2 (Callahan et al., 2016) pipeline (denoise-single in QIIME2 with --p-trunc length of 253, 253, 150, and 430 for datasets I, II, III, and IV, respectively) was applied to each dataset. This is a denoising step that performs quality filtering, chimera detection, and, optionally, merging of paired-end reads, which was not necessary in this case. The end-product of this pipeline is an Amplicon Sequencing Variant (ASV) table containing ASV abundance per sample. ASVs are exact sequence variants that provide higher resolution (Callahan et al., 2017) and hence, can resolve down to a single-nucleotide difference between target genes that subsequently helps in identification of bacteria down to the species level (only limited by the availability of reference genomes of known bacterial species) (Henschel et al., 2015; Lozupone & Knight, 2007; Lozupone et al., 2013). Once the ASV tables were obtained for each dataset, they were consolidated into a single unified table using the overlapping method (the sum procedure in QIIME2) to merge the tables with all the samples. In addition to merging the tables, this procedure identifies and enumerates exact matches in sequence and read length among the ASVs from different datasets. The resultant merged table possibly contains redundant ASVs. This redundancy is caused due to any of the following factors: 1) differences in read length among batches, which results in ASVs with an exact match in sequence but not in size failing to be merged; and 2) amplification of different molecular barcodes (i.e.,16S rRNA gene hypervariable region V3-V4 or V4), being considered redundant in the table since ASVs generated using different hypervariable regions will not be resolved using this method even when amplified from the same taxon.

Both types of batch effects were addressed before further analysis as follows. First, the single unified ASV table was subjected to open-reference clustering using the vsearch module (Rognes et al., 2016) in QIIME2 with identity percentage of 100% to ensure exact matches between sequences. The green genes 13_8 database (DeSantis et al., 2006; McDonald et al., 2012) was used for this method. The open-reference clustering technique first clusters the ASVs by aligning each with known 16S reference sequences from the database using the specified similar percentage, which is also called closed-reference clustering. It then performs a *de novo* clustering with the ASVs that could not be matched with any known sequences. The first step not only helps in collapsing the redundant ASVs varying in size, but it also handles the ASVs from different hypervariable regions by collapsing these ASVs into single long sequences. The *de novo* clustering then collapsed the varying size ASVs with exact sequence matches, since percentage identity chosen was 100%. Then, a fragment-insertion step was performed on the merged and clustered ASV table using the SATé-enabled phylogenetic placement (SEPP) method (Janssen et al., 2018). SEPP dramatically reduces the bias due to different hypervariable regions. Finally, the output abundance table, consisting of 80,140 non-redundant ASVs, was used for further analysis. The only remaining batch effect in the data was due to differences in sequencing experiments that resulted in varying sequencing library sizes. Usually, for traditional meta-analysis, such batch effects are addressed using standard normalization methods, such as rarefaction, assessing abundance frequency, etc. As discussed in the next section, we used a method that resolved this batch effect in the dataset.

### Data normalization

Microbiome data is sparse, heterogeneous, and compositional in nature (Busato et al., 2023). The compositionality of the microbiome data cannot be ignored, and appropriate methods should be applied when accounting for it. Gloor et al. (2017) discussed the potential issues originating from the use of conventional standardization methods for compositional data and suggested more sophisticated, appropriate replacements for them. The impact of data normalization prior to analysis is not only limited to microbiome meta-analysis but is also applicable to machine learning (ML). In fact, many studies have examined the importance of using normalized data for supervised ML classification and have reported improved classification performance on various ML algorithms when using normalized data (Jain et al., 2005; li & Liu, 2011; Noda, 2008; Singh & Singh, 2020). We used Centered Log-Ratio (CLR) transformation, a method designed for compositional data, which is considered to be the best performing approach for data normalization for the ML analysis using microbiome data (Kubinski et al., 2022). CLR transforms the data by obtaining the logarithm of the ratio between the ASV abundances and their geometric mean. A pseudo-count of 1 was added to replace zeroes. CLR transformations (Kubinski et al., 2022) were applied on the final ASV table with raw counts using the compositions package (van den Boogaart & Tolosana-Delgado, 2008) in R (R Core Team, 2013). It is to be noted that while in most machine learning based analyses, the ASV tables are collapsed to taxonomic levels before analysis, we decided to apply machine learning using the ASVs themselves since the former method filters out thousands of unknown bacterial species prior to analysis and, also, the models generated using the ASV’s themselves, will be useful in the future when more bacterial reference genomes are available, allowing retrospective identification of ASV’s currently unassigned to a taxa.

### Exploratory data analysis (EDA) and identification of mislabeled samples

Before proceeding to the ML analysis, various statistical and bioinformatics methods were used to explore the microbiome data. Alpha diversity provides insight into microbial richness and evenness within samples, while beta diversity reveals differences in microbial communities between samples. We performed alpha- and beta-diversity analyses in QIIME2. Bacterial compositions, at the phylum level, among the cultivars, countries, and continents were compared. For this, the final ASV table (prior to CLR transformation) was first rarefied to an even depth of 1,000 sequences per sample. Furthermore, taxonomic classification was performed using classifiers trained toward GreenGenes2 (McDonald et al., 2023) to assign the taxonomy to ASVs. The abundances among the samples within a group were merged using the mean-ceiling method of QIIME2’s feature-table merge. The composition comparison among the top 10 phyla, where all other identified phylum grouped as “Other”, was visualized as stacked bar plots using a Python module, Dokdo module version 1.16.0 (Lee, 2022).

For beta diversity, we used two distance metrics: Bray-Curtis and weighted UniFrac. The Bray-Curtis dissimilarity is a measure of community composition that accounts for abundance, while the weighted UniFrac distance incorporates phylogenetic relationships between observed organisms, weighting them by relative abundance. With these metrices, we constructed three-dimensional Principal Coordinates Analysis (PCoA) plots using the Emperor tool (Vázquez-Baeza et al., 2013) in QIIME2. These plots provide a spatial representation of the microbial communities per sample, which can be further color-coded based on specific interest groups, illustrating the distances and dissimilarities between samples based on their microbial composition. Visual examination of PCoA performed using the initial 885 samples showed 96 samples, which formed a separate cluster from the rest of the samples and were considered potential outliers (Supplementary Figure 1). All 96 samples were part of Dataset III which contained samples generated from microbiome data from the roots, leaves, or berries. Further inspection of these potential outliers revealed that these samples were misclassified or mislabeled as soil/rhizosphere. The evidence of the misclassification was obtained by examining a specific column in the metadata file named “Library Name”, where the name encodes the information about the source. Specifically, the library names contain the patterns, “gp”, “leav”, “root”, “bulk”, and “rhizo”, suggesting that these libraries were generated from “grapes”, “leaves”, “root”, “bulk soil”, and the “rhizosphere”, respectively. Additionally, out of the total 401 samples from this sequencing run, 88 were categorized as “rhizosphere”, 153 were categorized as “soil”, and 160 were categorized as “root”. Moreover, the number of samples displaying the patterns, “gp”, “leav”, “root”, “bulk”, and “rhizo”in the name are, respectively, 47, 49, 153, 64, and 88, which suggested that the leaves and grapes samples were classified as “soil”. These samples were removed from further analysis since there was sufficient evidence of potential mislabeling (See Supplementary Table 3 for all removed samples.). We used another beta diversity distance metric for the visualization of the remaining samples, Aitchison. This is an statistic approach highlighted for its robustness when dealing with compositional data since it takes into account the relative proportions of species, rather than their absolute abundances, thus controlling for different sample sizes (Martino et al., 2019). Essentially, it uses the CLR transformed abundances. Robust Aitchison principal component analysis was performed using DEICODE (Martino et al., 2019), and the plots were constructed using the Python matplotlib library. Finally, multivariate permutational analyses of variance (PERMANOVA) with 999 permutations were performed using QIIME2’s ADONIS module (Anderson, 2001), to determine whether groups of samples significantly differed from one another.

### Applying Machine Learning

The purpose of this study was to decipher the influence of the 3C’s -Continent, Country, and planted Cultivar on the assembly of plant microbiota, as represented by 16S amplicon data using supervised machine learning classification techniques. The dataset comprises tabulated 16S amplicon data with columns representing Amplicon Sequence Variants (ASVs) and rows representing individual samples. The entries within this table are CLR-transformed abundances. Class labels assigned to each sample included a continent, a country, and cultivar. Although commercial grapevines are almost exclusively planted in a grafted form, here cultivar was defined as the genotype of the scion (aerial part of the vine) since the rootstock information was not always available (ambiguous) for all the datasets. However, since the rootstock information was known for a small number of samples, we also trained the models to predict the scion/rootstock combination.

Despite encompassing samples from 5 continents, 13 countries, and 34 scion cultivars, the dataset revealed a pronounced class imbalance. As a result, two different approaches were used to build the ML models, taking into consideration the filtering criteria for sparse classes.

#### Approach A: Inclusion of Sparse Classes

In this approach, any class with a minimum of 3 samples was retained, maximizing the number of classes at the cost of potential over-representation of the majority class(es). This resulted in a dataset consisting of samples from 5 continents, 12 countries, and 19 scion cultivars (scion/rootstock combination predictions were not attempted), resulting in sample sizes of 788, 787, and 763 for continent, country, and scion cultivar classifications, respectively. The entire dataset was divided into a 75/25 train/test split. The 75% allocated for training was subjected to hyperparameter optimization using GridSearch with stratified repeated k-fold cross-validation (k = 3 and repeats = 5). The stratified split was used to ensure classes were split proportionally in training and test sets which was necessary in our case due to high class imbalance. The split proportions (and choice of k) included representatives from all classes in the test set since sparse classes were included. Additionally, our study is based on a previous study (Gobbi et al., 2022) that used a 75/25 train/test split. Finally, the optimal models identified through the grid search were then assessed on the naïve 25% test data.

#### Approach B: Exclusion of Sparse Classes

Here, only classes with at least 15 samples were retained for the prediction of continent, country and cultivar, and a minimum of 12 samples for the prediction of the rootstock/scion combination. This resulted in a dataset consisting of samples from 3 continents, 5 countries, 6 scion cultivars and 8 scion/rootstoock combinations, resulting in sample sizes of 781, 743, 685, and 470 for continent, country, scion cultivar and scion/rootstock combination classifications, respectively. This criterion ensured a more balanced representation of classes at the expense of reduced class diversity. Moreover, this approach was used to ascertain the performance and generalizability of our models. For this purpose, we used nested k-fold cross-validation (CV), which, besides assisting in hyperparameter tuning, helps prevent the overfitting of our models to the training data, ensuring they are well-tuned to naïve data. (Cawley & Talbot, 2010), we chose k = 5 (for both outer and inner loops) to ensure the 80/20 split.

### Choice of ML algorithms

The research undertook the task of classifying microbiome data using a suite of nine different machine-learning algorithms. Gobbi et al. (Gobbi et al., 2022) already provided evidence that Random Forests (RFs) could be successfully used for the classification of country/continent from soil microbiome data. We added more algorithms ranging from ensemble methods (Zhou & Zhou, 2021) such as Adaptive Boost (ADA), and Gradient Boosting Machine (GBM) to linear and kernel-based approaches, including Support Vector Machines (Cortes & Vapnik, 1995) with linear (SVML) and radial (SVMR) kernels. Furthermore, we incorporated probabilistic algorithms, Gaussian (GNB) and Bernoulli Naïve Bayes (BNB) (Zhang, 2004), as well as the instance-based method k-nearest neighbor (KNN) (Cover & Hart, 1967) and the more complex Neural Networks (NN), specifically multi-layer perceptron; (LeCun et al., 2015).

All algorithms, except NN, were implemented using Scikit-learn (v1.1.2) library (Pedregosa et al., 2011), while the NN models were built using TensorFlow (v2.8.0) library (Abadi et al., 2016) both in python (v3.8.2). Analysis was performed using the University of Kentucky’s High-Performance Compute clusters: the Lipscomb Compute Cluster (LCC) and the Morgan Compute Cluster (MCC).

### Fine-tuning with hyper-parameter optimization

To fine-tune our machine learning models, a grid search on hyper-parameters was carried out. This was achieved using Scikit-learn’s GridSearchCV function, which assessed every possible combination of hyper-parameter values specified. The specific hyper-parameters probed for each algorithm, as well as the values of the best-fit models, are provided in Supplementary Table 4.

The selection of hyper-parameters was algorithm-specific. Instead of starting with a default weak learner, the ADA was initiated with a strong learner (a decision tree with hyper-parameters obtained from the best fit from the RF). The BNB stands apart from other machine learning techniques in its need for data adhering to a Bernoulli distribution, characterized by discrete values. This distinction aligns with certain statistical distance methods for microbiome data meta-analysis, like the Jaccard distance, which operates on a presence-absence basis (Real & Vargas, 1996). Testing via BNB thus provided insights into the efficacy of classification grounded in mere presence or absence, as opposed to actual abundance. Specifically for BNB, the dataset was transformed into the presence-absence matrix where a value of 1 is assigned if an ASV exists in that sample and 0 otherwise, but losing any information potentially linked to abundance in the process.

### Addressing Class Imbalance

When datasets are imbalanced, models tend to be biased towards the majority class. The “Class weights” provide a way to tell the ML models to “pay more attention” to samples from underrepresented classes (Anand et al., 2010). Class weights are provided as hyper-parameters in most ML methods and were calculated here using “balanced” heuristic (King & Zeng, 2001) methodologies implemented in the Scikit-learn’s compute_class_weight’s function. Class weights were provided for all machine learning algorithms except KNN, BNB, and GNB in the analysis using approach A. For the analysis using approach B, a similar workflow was adopted except for the neural network (NN), where class imbalance was addressed by using “focal loss” (Lin et al., 2017).

### Evaluation Metrics in light of class imbalance

Given the inherent class imbalance in our dataset. We recognized the potential bias in accuracy-based performance metrics. To account for this and to provide a more comprehensive understanding of the models’ performance by calculating F1-macro and F1-weighted scores (Wegier & Ksieniewicz, 2020). The F-measure, or F-score is an analysis methodology useful in binary classified data that is a measure of a test’s, like Type-I errors in parametric statistics. The F1 metric combines the accuracy of both Precision and Recall. Precision (also known as Positive Predictive Value) measures the accuracy of the positive predictions made by the classifier. It is the ratio of true positive (TP) predictions to the total predicted positives, which includes both true positives and false positives (FP). A higher precision means that a higher proportion of positive identifications was actually correct, which is crucial in fields where false positives are a significant concern. Recall (also known as Sensitivity or True Positive Rate) assesses the model’s ability to correctly identify all actual positives from the data. It is calculated by dividing the number of true positives by the sum of true positives and false negatives (FN), the latter being actual positives that the model incorrectly labeled as negatives. High recall indicates that the model is adept at finding and correctly labeling the positive cases. F1-score is the harmonic mean of precision and recall, providing a single score that balances the two metrics, given by the equation:

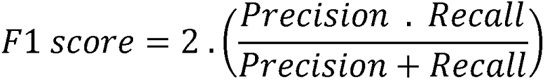

Where, 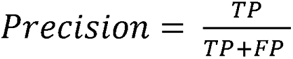, and 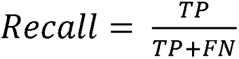

The F1-score reaches its best value at 1 (perfect precision and recall) and worst at 0, acting as a combined measure that assesses the accuracy and completeness of the positive predictions. The F1-macro score computes the F1 score for each class independently and then takes the average. This treats all classes equally, irrespective of their size, which makes it particularly useful for datasets with imbalanced classes. The F1-weighted score, on the other hand, computes the F1 score for each class independently but when it takes the average, it weights the F1 scores by the number of true instances for each class. This gives more weight to the larger classes in the dataset. We considered models with weighted F1-score ≥ 0.8 to be the best performing model. Additionally, we reported per class F1 scores which provides a granular view of model performance, ensuring that the evaluation is comprehensive and not just driven by the dominant classes.

### Neural Networks

In the two approaches for model building described above, the hyperparameter selection for all algorithms except neural networks remained largely consistent. Yet, when constructing neural networks, the strategies employed varied, especially in aspects like hyperparameter choice and addressing class imbalance, among others. This section delves into these distinctions.

Neural networks present a distinctive methodology within the machine learning landscape. Unlike traditional machine learning algorithms, neural networks necessitate the specification of an architecture, including the number of hidden layers, neurons per layer, and other related parameters such as number of epochs (i.e., how many single pass (forward and backward together) training iterations were performed which depends in turn on how the training data were partitioned into individual training batches). Two distinct strategies were used to identify the optimal neural network architecture. In the analysis using approach A, a heuristic approach was employed, involving a series of hit-and-trial experiments. Multiple architectures were empirically tested to gauge performance, and the configuration yielding the most promising results was selected. The final NN architecture used is presented (Supplementary Figure 2). The list of hyper-parameters used in the final model is listed (Supplementary Table 5).

To ensure a more systematic search for the optimal architecture, we employed a grid search method for the analysis using approach B. A custom Python script was used to emulate the scikit-learn’s GridSearch functionality. The hyper-parameters used in grid search are listed in Supplementary Table 6.

Additional measures were incorporated into the architecture to enhance the model’s generalizability and to counteract potential overfitting. In our architecture, we incorporated Dropout Layers (Srivastava et al., 2014). During training, certain neurons are randomly set to zero, forcing the model to forget information and preventing it from overfitting the training data. Dropout layers used in approach A are shown in the network architecture (Supplementary Figure 2). In approach B, a dropout layer was added after each hidden layer.

Finally, the methods employed to address class imbalance varied between the two approaches. Approach A utilized the class weight methods previously discussed. In Approach B, we used the “focal loss” as a loss function instead of standard sparse categorical crossentropy (SCCE; (Zhang & Sabuncu, 2018)). The focal loss method, introduced by Lin et al., (2017) in the context of object detection, modifies the standard cross-entropy loss function by adding a modulating factor to down-weight easy examples and focus the learning on hard negatives. It essentially applies a scaling factor to the cross-entropy loss, with the scaling factor tending to zero for well-classified examples. By doing so, focal loss ensures that the classifier does not become overwhelmed by the majority class and ignores the minority class. It encourages the model to pay more attention to difficult, misclassified cases that are often found in the minority class, improving performance in imbalanced datasets.

### Identification of important taxonomical predictors for all class types

We used the built-in mechanisms in Random Forests, Gradient Boosting, and Adaptive Boosting algorithms to evaluate and quantify the significance of each feature in the dataset with respect to the predictive power of the model. These models compute feature importance as part of their algorithmic process, allowing us to understand which variables contribute most to the model’s decision-making. In Random Forests, feature importance is determined by looking at how much each feature decreases the impurity of the split (e.g., Gini impurity) across all trees in the forest. Gradient Boosting computes feature importance by measuring each feature’s contribution to minimizing the loss function, often weighted by the number of times a feature is used in the base learners.

Similarly, AdaBoost assigns a weight to each feature based on how well it helps to correct the errors of the weak learners in the ensemble. Leveraging these intrinsic methods, we extracted the computed importances to identify and rank the ASVs that are most influential for the predictions of the different class types analyzed.

We began by setting a threshold for feature selection at the level of average importance, as determined across all features within each model. Features that exhibited importance scores above this average were deemed significant and were thus selected for further consideration. This process helps to filter out noise and less informative features, focusing the analysis on those variables that have a more substantial impact on the model’s predictive performance. After applying this criterion independently to each of the three models, we compiled the lists of selected features to identify commonalities among them. The overlapping features—those identified as above-average in importance across Random Forest, AdaBoost, and Gradient Boosting—were highlighted as being particularly relevant. We then used the sequences of these important ASVs and performed taxonomic classification using classifiers trained using GreenGenes2 and SILVA v138.

### Code availability

All code used in this study can be freely accessed at: https://github.com/Lakshay-Anand/grapevine-microbiome-ml.git

## Results

The primary aim of this study was to harness the predictive capabilities of machine learning (ML) to determine the influence of grapevine scion genotype and geographical origin on the composition of the vineyard soil microbiome. This was approached with the hypothesis that despite the predominant role of the environment, if the vine’s genotype significantly impacts soil microbiome assembly, then the scion’s and rootstock influence should be discernible through ML models.

### Microbiome samples grouped based on all C’s

Principal coordinate analysis (PCoA) plots for the microbiome data showed grouping of samples based on continent and country of origin, and the scion’s cultivar (Figure 1A-C). There was a significant effect (Adonis, p <= 0.001) of continent, country, and scion cultivar on soil microbial assemblage. Overall, geographical location explained a higher degree of variation in microbial composition (Continent: R^2^ = 0.215, p = 0.001; Country R^2^ = 0.205, p = 0.001) than scion cultivar (R^2^ =0.122, p = 0.001).

### Algorithm Performance

Two distinct approaches to data filtering criteria were undertaken to validate the robustness of our models against class imbalance. Approach A retained classes with a minimum of three samples, while approach B narrowed it down to classes with at least 15 samples.

#### Geographical origin prediction

For the first approach, all algorithms, except BNB and KNN, achieved F1-weighted scores of ≥ 0.9 for continent classification and > 0.85 for country classification when tested on the withheld 25% test data (Supplementary Figure 4 and Supplementary Table 7, and Supplementary Figure 5 and Supplementary Table 8, for continent and country, respectively). The average F1-weighted scores for models generated excluding sparse classes (approach B) exceeded 0.8 for all continents and countries, with the exceptions being BNB and KNN (Supplementary Figure 6).

#### Scion genotype prediction

For cultivar classification using approach A, only RF, GBM, SVML, and NN surpassed F1-weighted scores of ≥ 0.8 on the excluded 25% of the test data (Figure 3 and Supplementary Table 9). Using approach B the average F1-weighted score exceeded 0.8 for all retained cultivars regardless of predictive model, barring BNB and KNN (Supplementary Figure 6).

**Figure 2.**
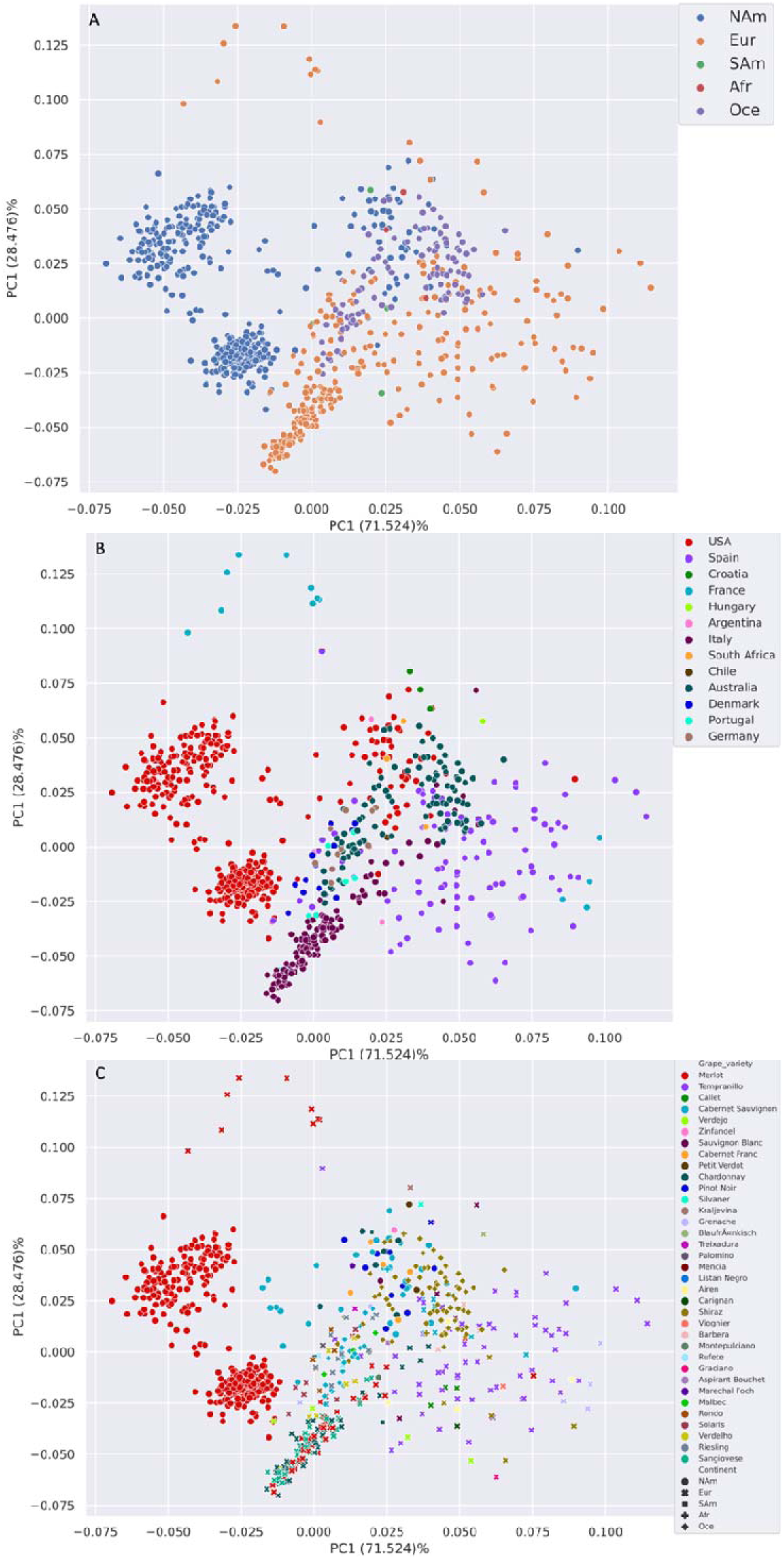
Effect of continent, country, and planted cultivar on vineyard microbiome. **(A-C)** PcoAs plot constructed using Aitchison pairwise distances for all samples. Samples are color coded according to the continent (A), and country (B) of origin, and the scion’s cultivar (C).

**Figure 3.**
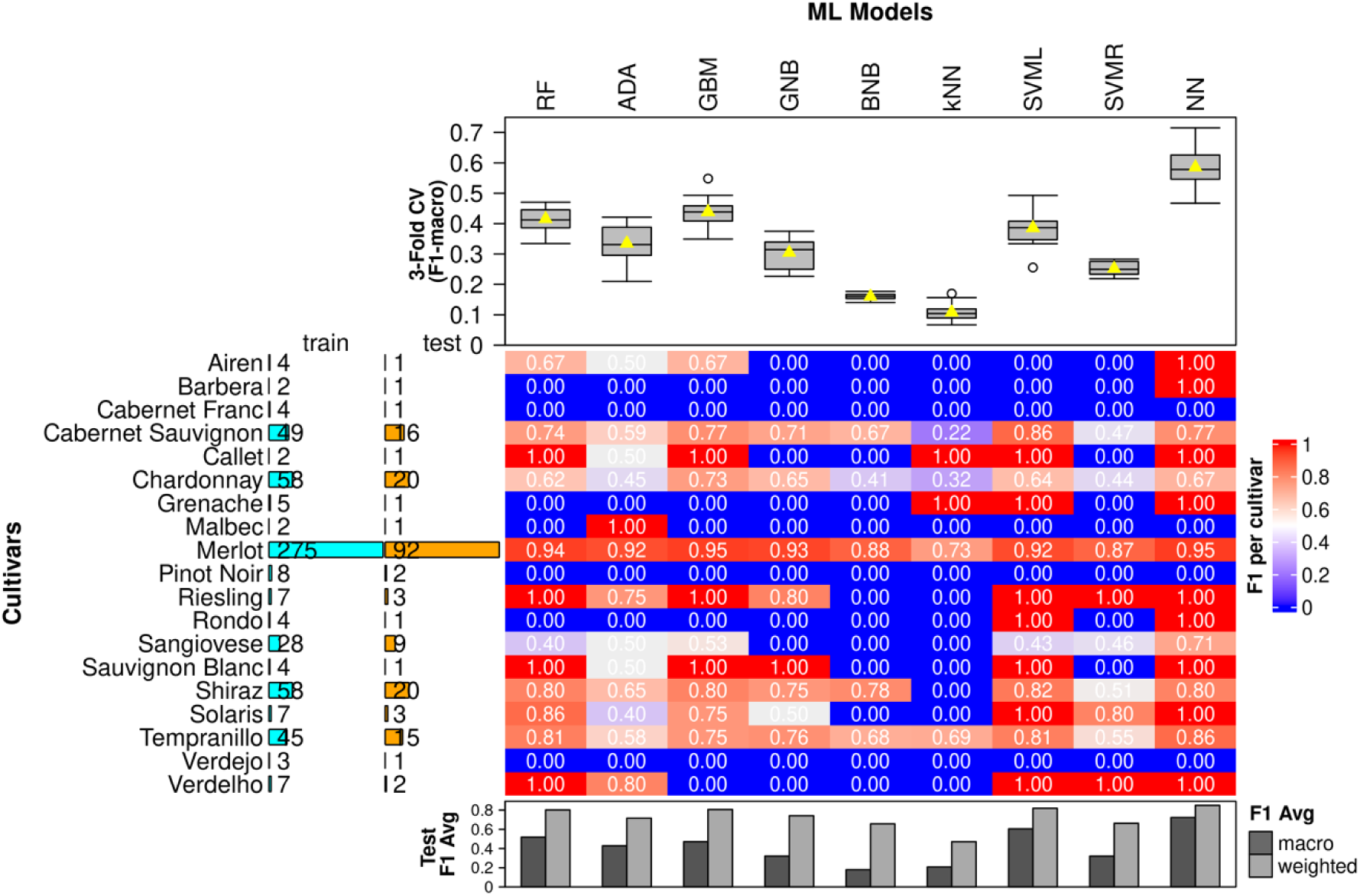
Performance results for cultivar classification using approach A. Performance results of the nine machine learning algorithms, Random Forests (RF), Adaptive Boost (ADA), Gradient Boost (GBM), Support Vector Machines with linear (SVML) and radial (SVMR) kernels, Gaussian (GNB) and Bernoulli Naïve Baye (BNB), k-Nearest Neighbor(KNN), and Neural Networks (NN) on classification of 19 scion cultivars using soil microbiome data The F1 scores for each continent for each ML method are represented as a heatmap. Above the heat map: The repeated stratified 3-fold Cross-Validation (CV) results are shown as boxplots where the average F1-macro scores are represented as yellow triangles. Left of the heatmap: number of train (light blue) and test (orange) samples, respectively, for each continent after a 75%/25% train-test split where each continent class has a minimum of three samples. Below the heatmap: F1 averages (macro and weighted) are depicted as bar graphs. The model was tested on 25% withheld test set.

Neural Network was the best predictive model for Continent, Country, and scion genotype according to cross-validation results (Supplementary Figure 4 & 5 (continent and country, respectively), and Figure 3 (Scion genotype)), test scores (Supplementary Table 7-9), and the statistical comparison of models using the Wilcoxon rank-sum test (Supplementary Figure 7). Neural Network algorithm trained using approach B showed higher performance for all class types (Continent, Country, and scion genotype) than approach A (See confusion matrices for all cases in Supplementary Figures 8-10).

#### Scion/rootstock genotype prediction

Analysis of the predictive capacity of the NN model when using the scion and rootstock genotype independently and combined showed all had high predictive power (Average F1-weighted score of 0.91 ± 0.004, 0.93 ± 0.004, and 0.94 ± 0.004 for scion, scion/rootstock combined, and rootstock, respectively). Paired T-test showed that the predictive power of the model built using rootstock as label was significantly greater than those built using scion (P-val < 0.0001) and scion/rootstock combined (P-val < 0.05) (Figure 4A). When trained with scion/rootstock combination as class, the NN model’s overall accuracy was 97% (see Supplementary Table 9 for the rest of the performance scores). Moreover, the NN model was capable of classifying microbiomes with high accuracy (Accuracy > 83%) for all but one scion/rootstock combination (i.e., Chardonnay-3309C) (Fig. 17). However, although the accuracy in prediction for such a class was 33%, the model confused this sample to a class with the same scion but a different rootstock (i.e., Chardonnay-420A) (Figure 4B). Similarly, the only other class not showing 100% accuracy, Sangiovese-420A, was predicted as the correct rootstock but the incorrect scion (i.e., Chardonnay-420A).

**Figure 4.**
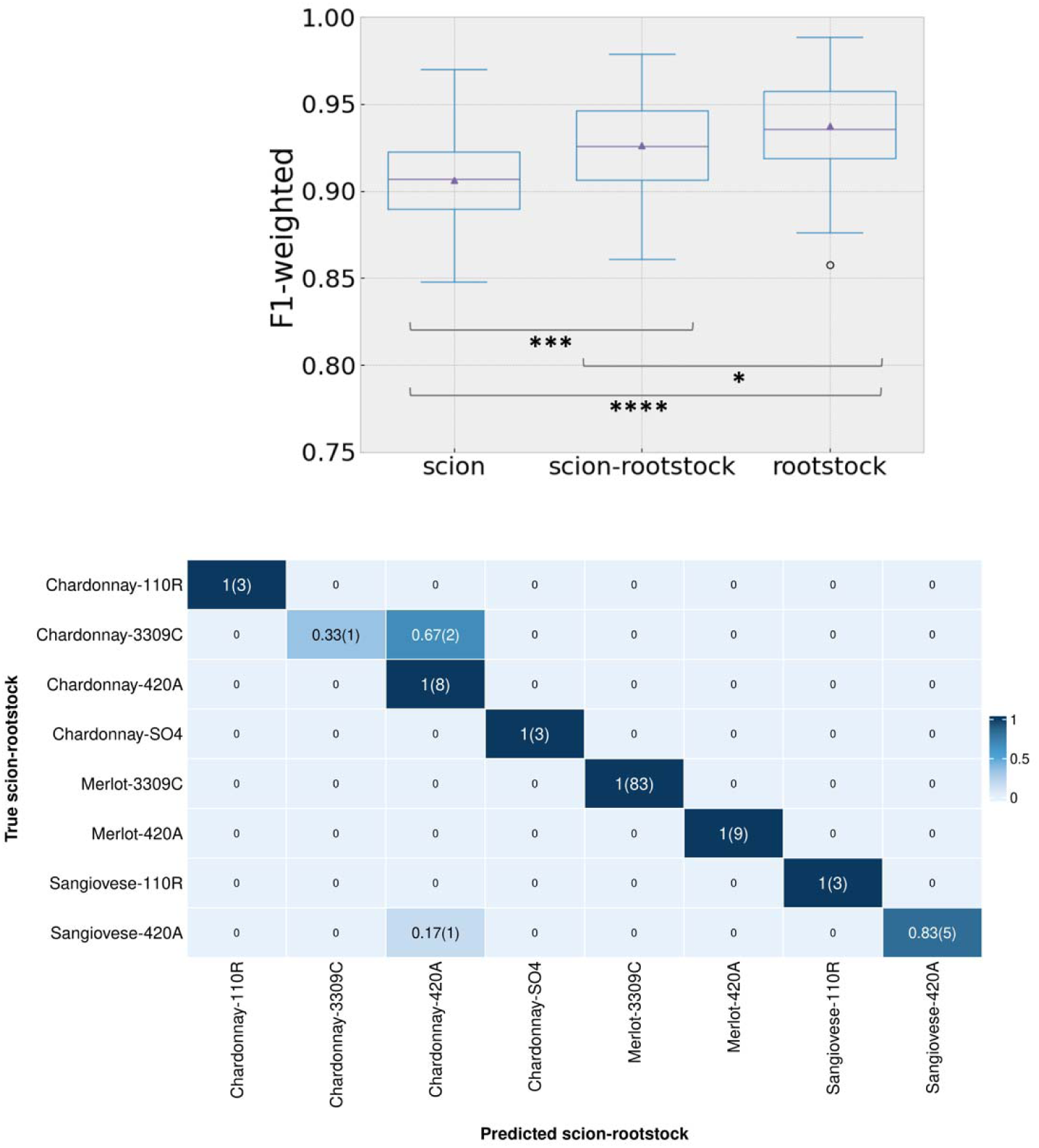
Vine soil microbiome contains the signal of both the scion and the rootstock. (A) Neural Networks model performance predicting scion, scion/rootstock combination, and rootstock genotypes. Total number per model classes: 6 scions, 4 rootstocks, and 8 scion-rootstock combinations. Asterisks indicate p-values of * < 0.05, *** < 0.001, and **** < 0.0001. (B) Confusion matrix for NN model for scion-rootstock combination prediction. Ratio of correctly predicted and falsely predicted scion-rootstock combinations using the Neural Networks (NN) model. The approach utilized nested cross-validation (CV), where the inner CV loop is used for hyperparameter optimization to return the best model, and the outer CV loop with 5-fold 10-repeats is used to test those best models. The confusion matrix generated from one of these models is depicted. The heatmap depicts the confusion matrix normalized over the true class, and the actual count is depicted within parentheses.

### Identification of important taxonomical predictors for all class types

A total of 1630, 1969, and 3415 significant features were identified for continent, country, and scion genotype, respectively using Random Forests, Gradient Boosting, and Adaptive Boosting (Figure 5 and Supplementary Tables 11-19). For all class types, Random Forest presented the highest number of features, while Adaptive Boosting presented the lowest (Figure 5). Taxonomic classification using classifiers trained on GreenGenes2 and SILVA v138 resulted in small discrepancies in the predicted taxa at the genus and species level, but in general both databases yielded very similar results (Supplementary Tables 11-19).

**Figure 5.**
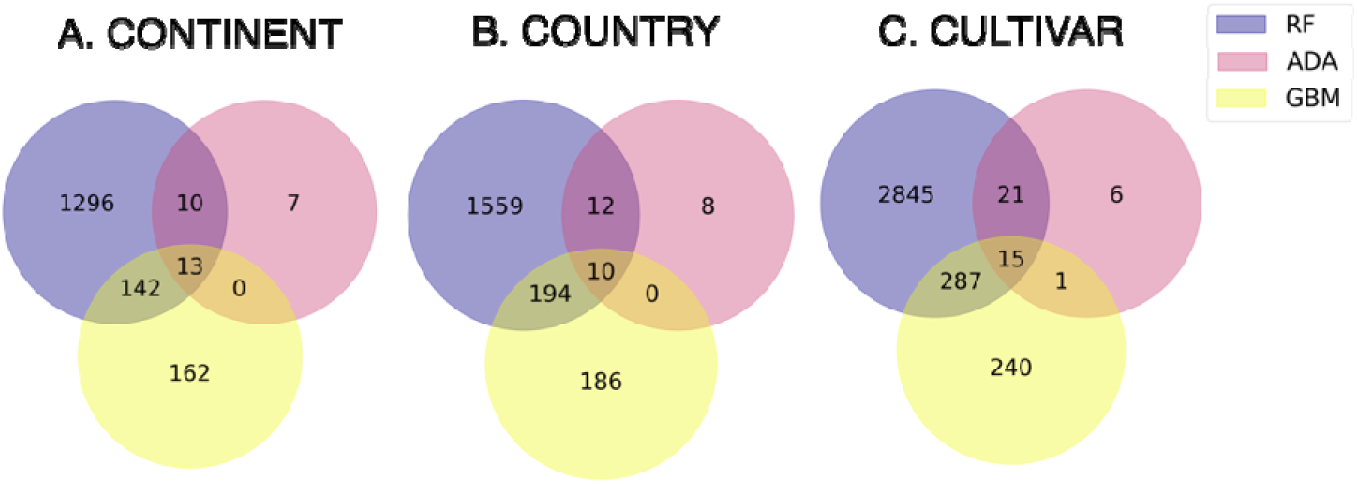
Overlapping most informative ASVs identified using three ML models. The features with importance above the average importance score were selected as important feature for the model. The overall of these features for the three ML models, Random forest (RF), adaptive boost (ADA), and gradient boost (GBM) is represented a Venn diagram.

Finally, only a small proportion of the significant features identified were common to all three algorithms (0.8% (13/1630), 0.5% (10/1969), and 0.4% (15/3415) for continent, country and scion genotype, respectively) (Supplementary Tables 20-22 for a complete list of overlapping features).

## Discussion

The classification of microbiome data by machine learning, focusing on the 3C’s—Continent, Country, and Cultivar—yielded models with considerable predictive capacity. Despite an imbalanced dataset, measures were taken to avoid overfitting towards the predominant class. Models such as RF, ADA, GBM, GNB, SVML, SVMR, NN demonstrated substantial predictive power across all classes (F1-weighted scores > 0.8), underscoring the robustness of the machine-learning frameworks utilized and the geographic and genotypic signals present in the microbiome data. In contrast, Bayesian Network Based (BNB) and K-Nearest Neighbors (KNN) models did not meet the benchmark established by other algorithms, potentially due to their model-specific characteristics, sensitivity to dataset imbalances, or the high dimensionality of the data (Pestov, 2013)

Our predictions of sample provenance corroborated the results by Gobbi et al. (2022), who applied the Random Forest algorithm to classify geographical origin using microbiome data. However, their methodology primarily relied on prediction accuracy, which can be deceptive in imbalanced classes (Grandini et al., 2020). Our approach enhances robustness by utilizing additional datasets for a more diverse training environment and employing nine different machine learning models. Furthermore, our evaluation framework, which incorporates F1-scores, offers a more reliable and generalizable assessment (Wegier & Ksieniewicz, 2020).

Contrasting with the findings of Marasco et al. (2022), our study challenges the assertion that the combination of rootstock and scion is the most significant influence the diversity of the grapevine root microbial community. We discovered that rootstock alone provides the highest predictive power, highlighting its importance in microbiome shaping and implications for viticulture, particularly in rootstock selection to promote beneficial microbial communities. These findings contribute a new dimension to the dialogue concerning factors influencing the grapevine root microbiome and underline the complexity of grapevine-microbe interactions.

We diverged from the convention of simplifying microbial data into taxonomic units, choosing instead to build models on amplicon sequence variants (ASVs) for a more detailed analysis. Using ensemble models, we extracted significant features that revealed inconsistencies between taxonomic classifications across different databases, suggesting that taxonomic reliance may lead to misinterpretations (Jasner et al., 2021; Kubinski et al., 2022; Namkung, 2020). The robustness of sequence-based models is highlighted, with the potential for reclassification as databases improve, proposing neural network models for future refined analysis.

The implications of this study are profound for viticulture and agriculture, especially in the face of challenges such as climate change. Understanding and manipulating plant-microbiota relationships can lead to sustainable practices, shifting breeding programs towards considering the holobiont as the breeding target, instead of just the plant genotype (Corbin et al., 2020). Our investigation paves the way for using machine learning to identify genes affecting crop microbiome assembly, aligning breeding programs with ecological principles for sustainable crop production.

Future research must validate these results in diverse vineyards and environmental conditions, accounting for the dynamic nature of microbial communities. Extending studies to various seasons and years, and exploring the influence of different rootstocks on the soil microbiome, will improve predictive model robustness and applicability. This study highlights machine learning’s potential in discerning grapevine cultivar and geographical origins from soil microbiota, supporting the hypothesis that machine learning can identify plant genes influencing microbiome composition despite environmental influences.

