## Supplementary Figures for "Grapevine Rootstock and Scion Genotypes’ Symbiosis with Soil Microbiome: A Machine Learning Revelation for Climate-Resilient Viticulture"

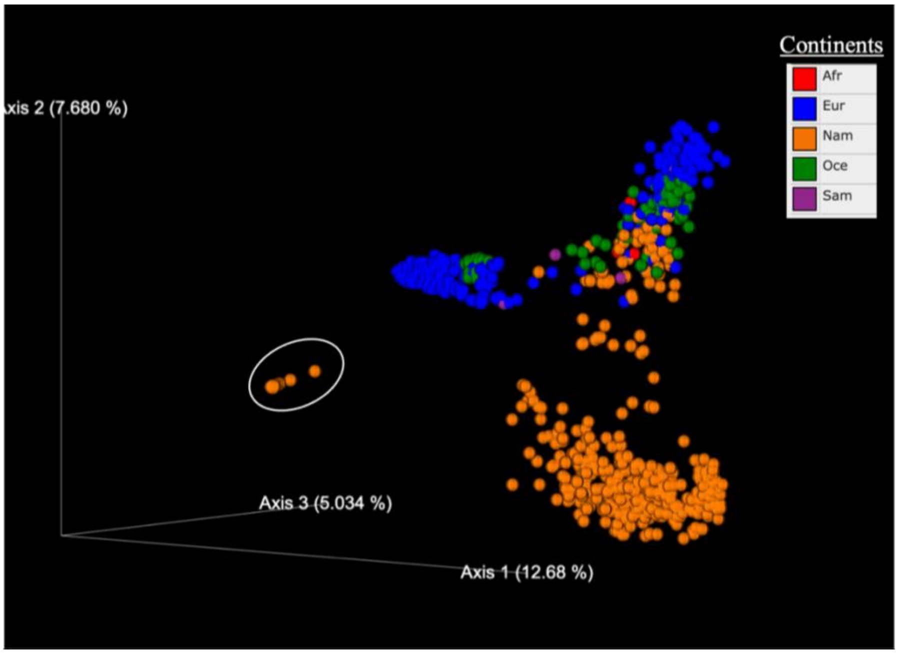


**Supplementary Figure 1. Detection of potential mislabelled samples.** PcoA analysis of 885 soil microbiome samples and visualized according to their continent of origin (Red = Africa; Blue = Europe; Orange = North America; Green = Oceania; and Purple = South America) using QIMME2 emperor. Samples highlighted with white circle were identified as potential.

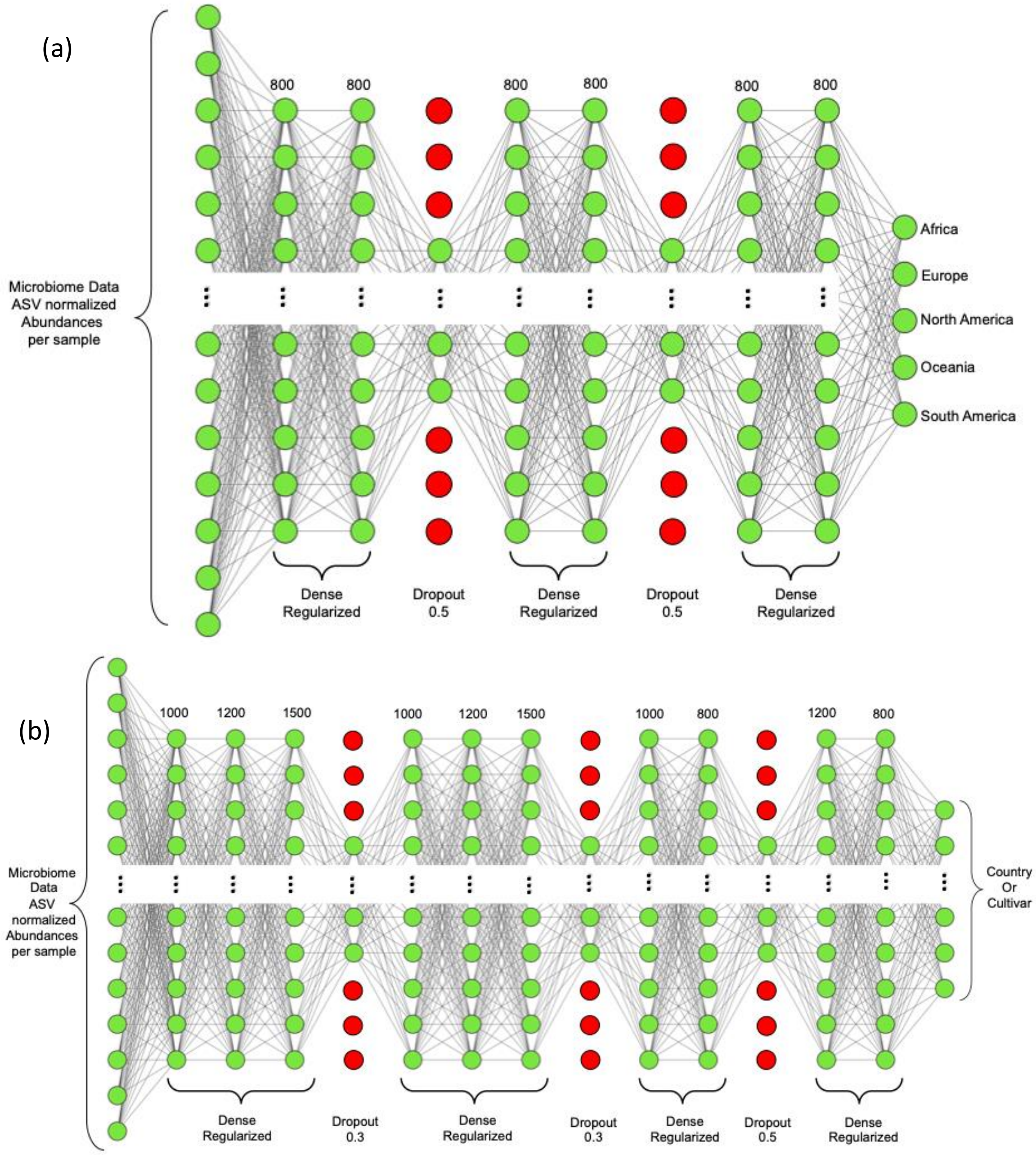


**Supplementary Figure 2. Architecture of Neural Networks.** Depictions of the NN architecture of the classifiers used for (a) continent and (b) country/cultivar classification respectively. Hidden layers are fully-connected layers (Dense) with L2 kernel regularization and neurons represented as green circles/nodes (number of neurons for the layer shown on top of the respective layer). The dropped-out neurons are represented as red nodes in the Dropout layers. NN architecture schematics are constructed using the NN SVG tool (https://alexlenail.me/NN-SVG/).

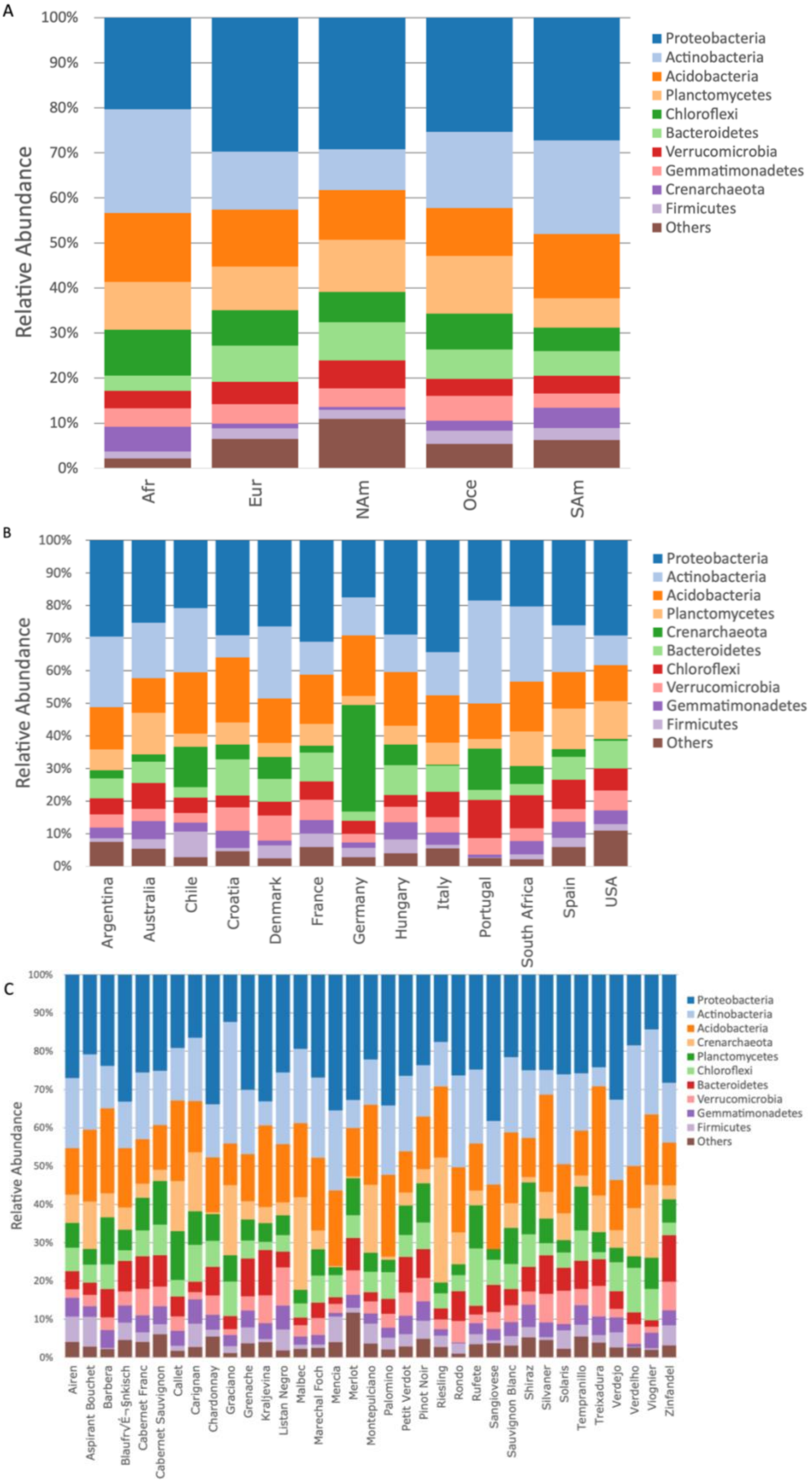
A
Relative Abundance
100%
90%
80%
70%
60%
50%
40%
30%
20%
10%
0%
Afr
Eur
NAm
Oce
SAm
Proteobacteria
Actinobacteria
Acidobacteria
Planctomycetes
Chloroflexi
Bacteroidetes
Verrucomicrobia
Gemmatimonadetes
Crenarchaeota
Firmicutes
Others
B
Relative Abundance
100%
90%
80%
70%
60%
50%
40%
30%
20%
10%
0%
Argentina
Australia
Chile
Croatia
Denmark
France
Germany
Hungary
Italy
Portugal
South Africa
Spain
USA
Proteobacteria
Actinobacteria
Acidobacteria
Planctomycetes
Crenarchaeota
Bacteroidetes
Chloroflexi
Verrucomicrobia
Gemmatimonadetes
Firmicutes
Others
C
Relative Abundance
100%
90%
80%
70%
60%
50%
40%
30%
20%
10%
0%
Airen
Aspirant Bouchet
Barbera
BlaufrVÉ¬§nkisch
Cabernet Franc
Cabernet Sauvignon
Callet
Carignan
Chardonnay
Graciano
Grenache
Kraljevina
Listan Negro
Malbec
Marechal Foch
Mencia
Merlot
Montepulciano
Palomino
Petit Verdot
Pinot Noir
Riesling
Rondo
Rufete
Sangiovese
Sauvignon Blanc
Shiraz
Silvaner
Solaris
Tempranillo
Treixadura
Verdejo
Verdelho
Viognier
Zinfandel
Proteobacteria
Actinobacteria
Acidobacteria
Crenarchaeota
Planctomycetes
Chloroflexi
Bacteroidetes
Verrucomicrobia
Gemmatimonadetes
Firmicutes
Others

**Supplementary Figure 3**. Effect of continent, country, and planted scion cultivar on bacterial Phylum abundance in vineyard soil microbiomes. Bacterial community composition in vineyard soil samples planted in different continents (A), countries (B), and with different cultivars (C). Composition is characterized to the phylum level (top 10 taxa).

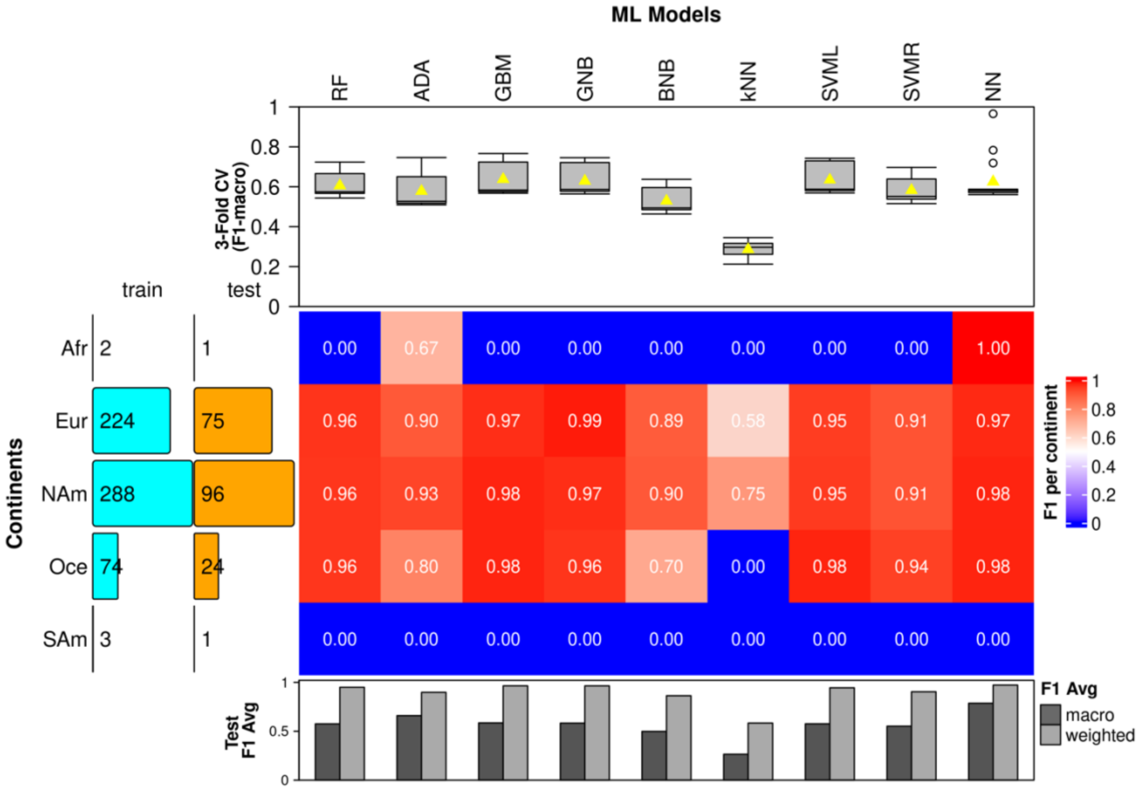


**Supplementary Figure 4**. Performance results for continent classification using approach A. Performance results of the nine machine learning (ML) algorithms, Random Forests (RF), Adaptive Boost (ADA), Gradient Boost (GBM), Support Vector Machines with linear (SVML) and radial (SVMR) kernels, Gaussian (GNB) and Bernoulli Naïve Bayes (BNB), k-Nearest Neighbor(KNN), and Neural Networks (NN) on classification of five continents ( Africa (Afr), Europe (Eur), North America (NAm), Oceana (Oce), and South America (SAm)) using soil microbiome data. The F1 scores for each continent for each ML method are represented as a heatmap. Above the heat map: The repeated stratified 3-fold Cross-Validation (CV) results are shown as boxplots where the average F1-macro scores are represented as yellow triangles. Left of the heatmap: number of train (light blue) and test (orange) samples, respectively, for each continent after a 75%/25% train-test split where each continent class has a minimum of three samples. Below the heatmap: F1 averages (macro and weighted) are depicted as bar graphs. The model was tested on an excluded 25%of the test set.

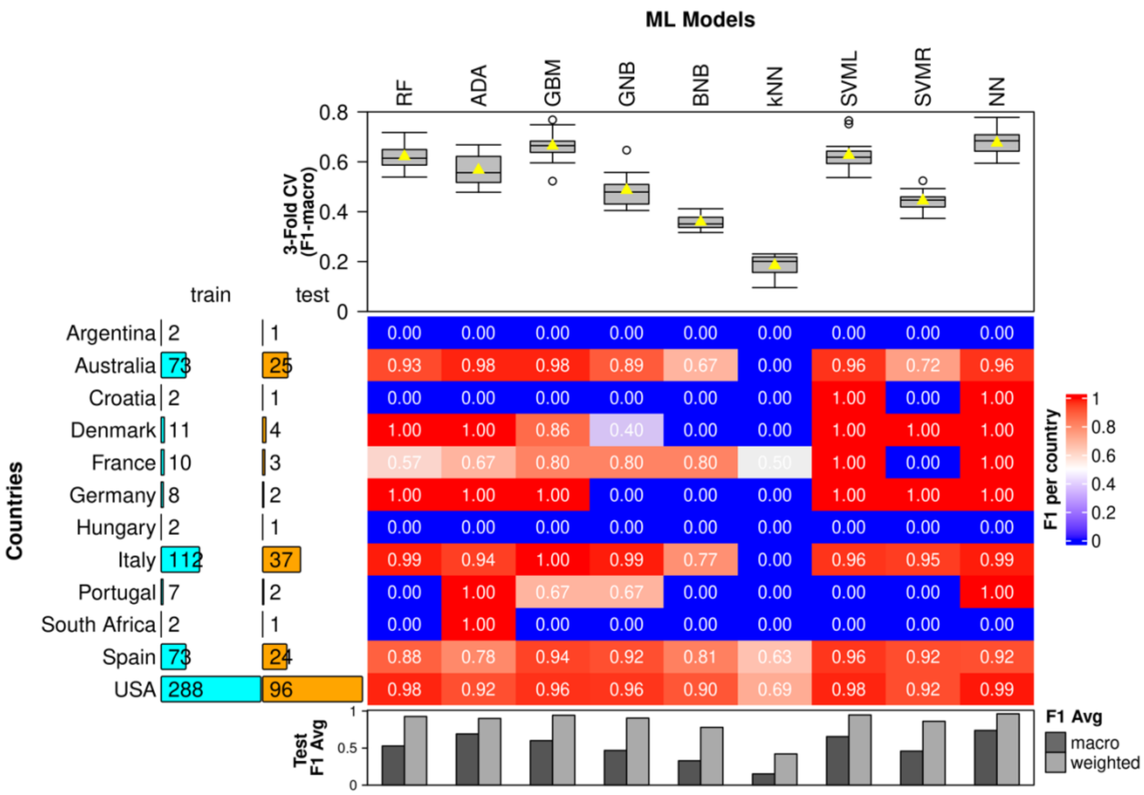


**Supplementary Figure 5**. Performance results for country classification using approach A. Performance results of the nine machine learning (ML) algorithms, Random Forests (RF), Adaptive Boost (ADA), Gradient Boost (GBM), Support Vector Machines with linear (SVML) and radial (SVMR) kernels, Gaussian (GNB) and Bernoulli Naïve Bayes (BNB), k-Nearest Neighbor (KNN), and Neural Networks (NN) on classification of 12 countries using soil microbiome data. The F1 scores for each continent for each ML method are represented as a heatmap. Above the heat map: The repeated stratified 3-fold Cross-Validation (CV) results are shown as boxplots where the average F1-macro scores are represented as yellow triangles. Left of the heatmap: number of train (light blue) and test (orange) samples, respectively, for each continent after a 75%/25% train-test split where each continent class has a minimum of three samples. Below the heatmap: F1 averages (macro and weighted) are depicted as bar graphs. The model was tested on an excluded 25% out of the test set.

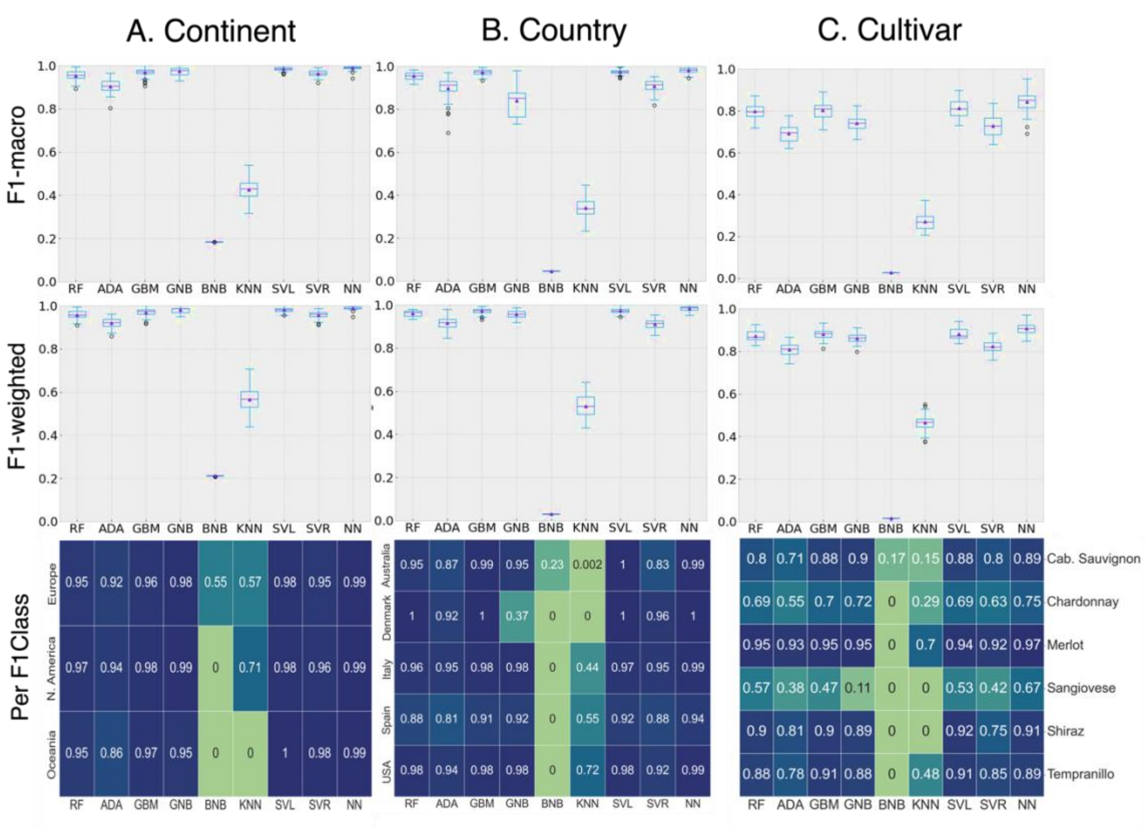


**Supplementary Figure 6**. Performance results for continent, country, and scion cultivar classification using approach B. Performance results of the nine machine learning (ML) algorithms, Random Forests (RF), Adaptive Boost (ADA), Gradient Boost (GBM), Support Vector Machines with linear (SVML) and radial (SVMR) kernels, Gaussian (GNB) and Bernoulli Naïve Bayes (BNB), k-Nearest Neighbor (KNN), and Neural Networks (NN) on classification of 3 continents (A), 5 countries (B), and 6 cultivars (C) using soil microbiome data. The results are obtained through nested cross-validation (CV), where the inner CV loop is used for hyperparameter optimization to return the best model, and the outer CV loop with 5-fold 10-repeats is used to test those best models. F1-macro (top panel) and F1-weighted (middle panel) show test results of 50 models from the outer loop as box plots (mean values depicted as purple triangles). The bottom panel depicts the average F1-score for each class and each ML model obtained from outer CV.

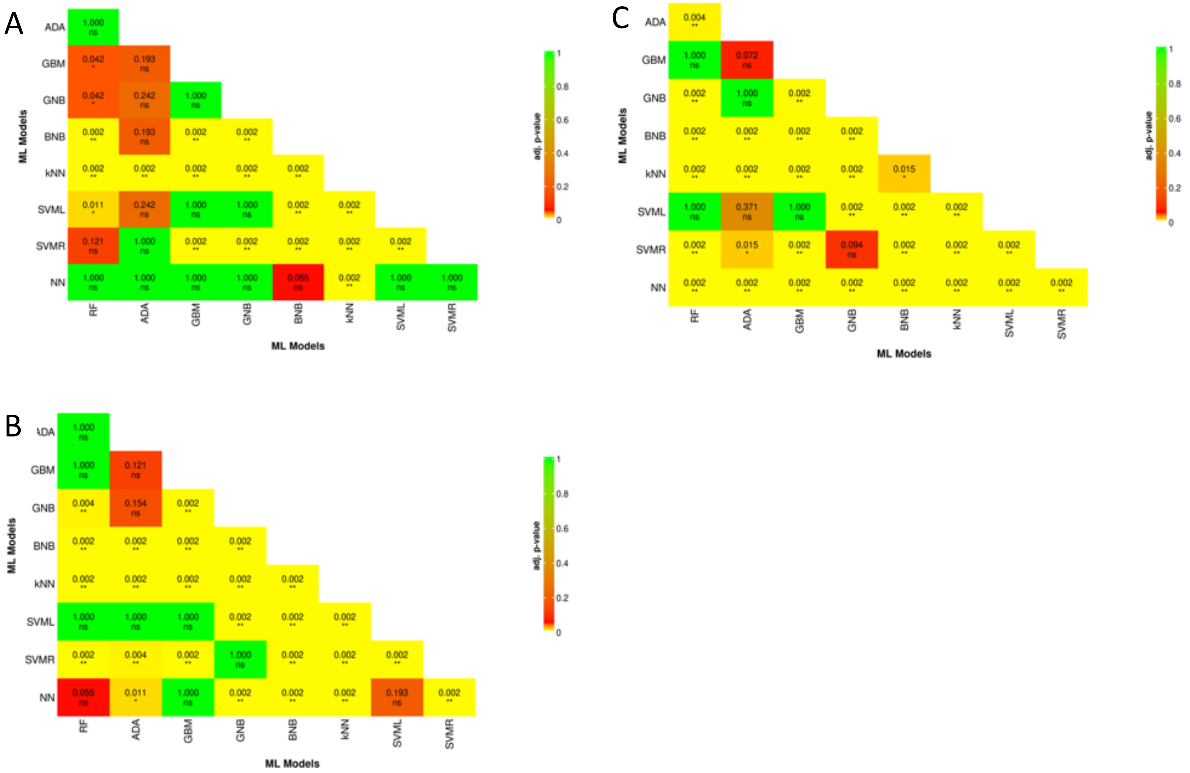


**Supplementary Figure 7**. Performance comparison among ML models for geographical origin and scion genotype prediction using approach A. Pairwise Wilcoxon ranksum test performed on the F1-macro scores from for the continent (A), country (B), and scion genotype classification by nine machine learning (ML) algorithms, Random Forests (RF), Adaptive Boost (ADA), Gradient Boost (GBM), Support Vector Machines with linear (SVML) and radial (SVMR) kernels, Gaussian (GNB) and Bernoulli Naïve Bayes (BNB), k-Nearest Neighbor (KNN), and Neural Networks (NN). The adjusted p-value scores (Bonferroni) are depicted where "ns" is "non-significant," and asterisks indicate ad. p-values of * < 0.05, ** < 0.01.

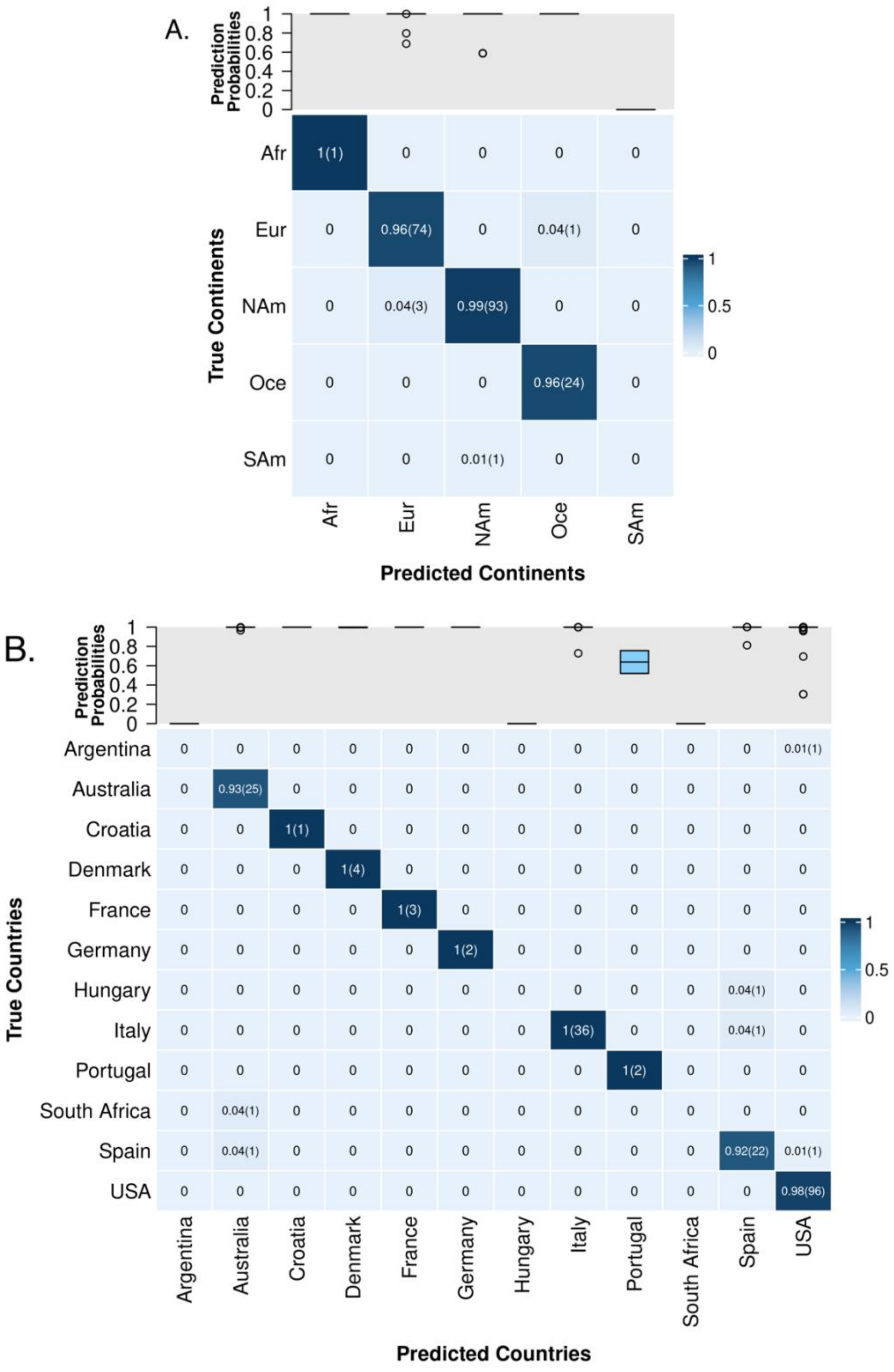


**Supplementary Figure 8**. Confusion matrices for geographical location prediction using NN for approach A. Confusion matrices showing the percentage of correctly predicted and falsely predicted continents (A) and countries (B) using the Neural Networks (NN) model tested on the 25% withheld test data. In this approach, 5 continents and 12 countries were chosen so that each class (continent or country) has a minimum of 3 samples. The heatmap depicts the confusion matrix normalized over the true class, and the actual count is depicted within parentheses. Probabilities for each prediction were retrieved from the NN model and visualized as boxplots against their respective predictions (continent or country) at the top of the heatmap.

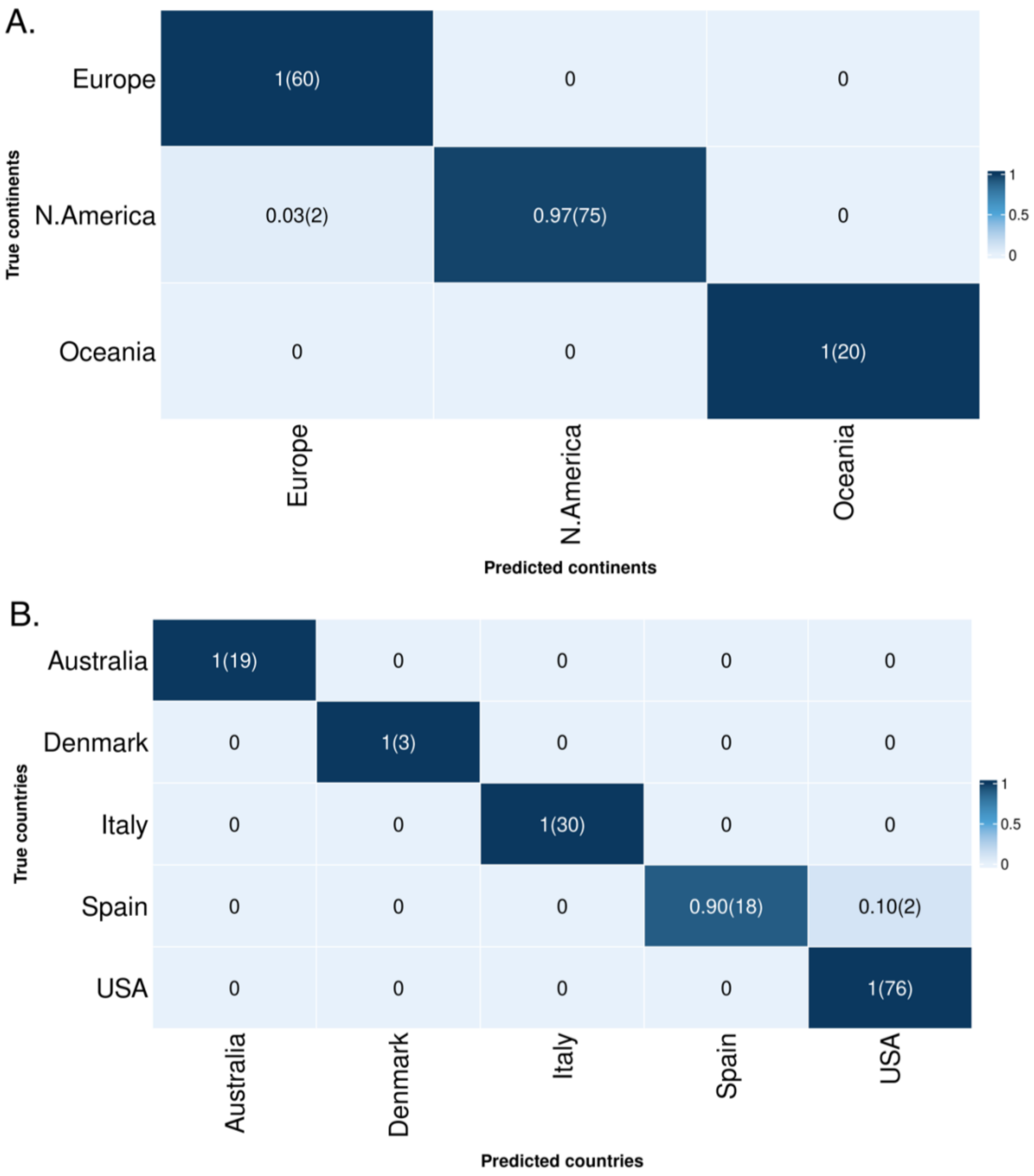


**Supplementary Figure 9**. Confusion matrices for geographical location prediction using NN for approach B. Confusion matrices showing the percentage of correctly predicted and falsely predicted continents (A) and countries (B) using the Neural Networks (NN) model. In this approach, three continents and five countries were chosen such that each class (continent or country) has a minimum of 15 samples. The approach utilized nested cross-validation (CV), where the inner CV loop is used for hyperparameter optimization to return the best model, and the outer CV loop with 5-fold 10-repeats is used to test those best models. Confusion matrix generated from one of these models are depicted. The heatmap depicts the confusion matrix normalized over the true class, and the actual count is depicted within parentheses.

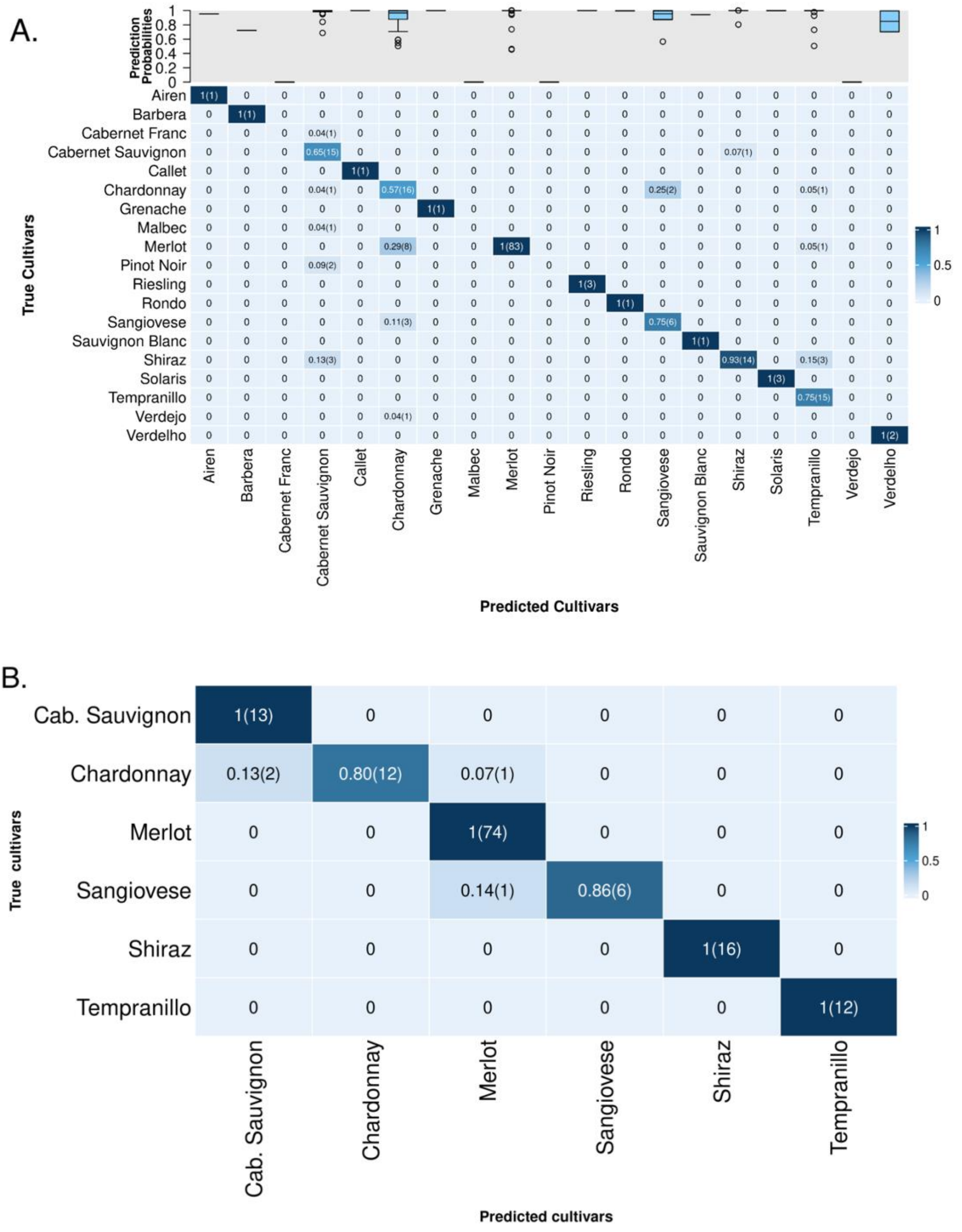
A.
Prediction Probabilities
True Cultivars
Predicted Cultivars
Airen
Barbera
Cabernet Franc
Cabernet Sauvignon
Callet
Chardonnay
Grenache
Malbec
Merlot
Pinot Noir
Riesling
Rondo
Sangiovese
Sauvignon Blanc
Shiraz
Solaris
Tempranillo
Verdejo
Verdelho
B.
True cultivars
Predicted cultivars
Cab. Sauvignon
Chardonnay
Merlot
Sangiovese
Shiraz
Tempranillo
1(13)
0.13(2)
0.80(12)
0.07(1)
1(74)
0.14(1)
0.86(6)
1(16)
1(12)

**Supplementary Figure 10**. **Confusion matrices for cultivar prediction for NN.** Ratio of correctly predicted and falsely predicted cultivars built using predictions from the Neural Networks (NN) model. Two distinct approaches are used in this study. The first approach divided the entire dataset into a 75%/25% train/test split. The confusion matrix generated from NN model tested on the 25% withheld test data is depicted in A. Probabilities for each prediction were retrieved from the NN model and visualized as boxplots against their respective predictions  at the top of the heatmap The second approach utilized nested cross-validation (CV), where the inner CV loop is used for hyperparameter optimization to return the best model, and the outer CV loop with 5-fold 10-repeats is used to test those best models. Confusion matrix generated from one of these modes is depicted in B.
