## Supplementary Table 1 for "Grapevine Rootstock and Scion Genotypes’ Symbiosis with Soil Microbiome: A Machine Learning Revelation for Climate-Resilient Viticulture"

**Supplementary Table 1**. **Summary of the Datasets used in this study and the public repository from which they were retrieved.** ENA is European Nucleotide Archive, SRA indicates NCBI Sequence Read Archive, and QIITA is a microbial study management platform. Sample#: Sample number. SE: Single end reads. PE: Paired-end reads

| **Dataset** | **Sample number** | **Source** | **Public Repository/ID** | **16S Ribosomal Hypervariable Region** | **Sequencing Type** | **Average Read Length (bps)** |
|---|---|---|---|---|---|---|
| I | 252 | (Gobbi et al., 2022) | ENA/ PRJEB40350 | V4, V3-V4 | SE and PE | 268.41 ± 1.28 |
| II | 66 | (Zhou et al., 2021) | SRA/ PRJNA601984 | V4 | PE | 254.14 ± 0.02 |
| III | 426 | (Zarraonaindia et al., 2015) | SRA/ PRJEB6677 QIITA / 1024 | V4 | SE and PE | 149.03 ± 0.08 |
| IV | 141 | (Marasco et al., 2022) | SRA/ PRJNA807110 | V3-V4 | PE | 454.10 ± 0.17 |
