## Supplementary Table 2 for "Grapevine Rootstock and Scion Genotypes’ Symbiosis with Soil Microbiome: A Machine Learning Revelation for Climate-Resilient Viticulture"

**Supplementary Table 2.** Metadata inlcued for each sample/dataset

| Dataset | Sample_ID | Country | Cultivar(scion) | Continent | Rootstock |
|---|---|---|---|---|---|
| I | ERR4588494 | Spain | Tempranillo | Europe | unkown |
| I | ERR4588495 | Spain | Tempranillo | Europe | unkown |
| I | ERR4588496 | Spain | Tempranillo | Europe | unkown |
| I | ERR4588497 | Spain | Tempranillo | Europe | unkown |
| I | ERR4588498 | Spain | Tempranillo | Europe | unkown |
| I | ERR4588499 | Spain | Callet | Europe | unkown |
| I | ERR4588500 | Spain | Tempranillo | Europe | unkown |
| I | ERR4588501 | Spain | Tempranillo | Europe | unkown |
| I | ERR4588502 | Spain | Callet | Europe | unkown |
| I | ERR4588503 | Spain | Tempranillo | Europe | unkown |
| I | ERR4588504 | Spain | Tempranillo | Europe | unkown |
| I | ERR4588505 | Spain | Tempranillo | Europe | unkown |
| I | ERR4588506 | Spain | Tempranillo | Europe | unkown |
| I | ERR4588507 | Spain | Tempranillo | Europe | unkown |
| I | ERR4588508 | Spain | Callet | Europe | unkown |
| I | ERR4588509 | Spain | Tempranillo | Europe | unkown |
| I | ERR4588510 | Spain | Cabernet Sauvignon | Europe | unkown |
| I | ERR4588511 | Spain | Tempranillo | Europe | unkown |
| I | ERR4588512 | Spain | Tempranillo | Europe | unkown |
| I | ERR4588513 | Spain | Tempranillo | Europe | unkown |
| I | ERR4588514 | Spain | Tempranillo | Europe | unkown |
| I | ERR4588515 | Spain | Verdejo | Europe | unkown |
| I | ERR4588516 | Spain | Tempranillo | Europe | unkown |
| I | ERR4588517 | Spain | Tempranillo | Europe | unkown |
| I | ERR4588518 | Spain | Verdejo | Europe | unkown |
| I | ERR4588519 | Spain | Tempranillo | Europe | unkown |
| I | ERR4588520 | Spain | Tempranillo | Europe | unkown |
| I | ERR4588521 | Spain | Tempranillo | Europe | unkown |
| I | ERR4588522 | USA | Cabernet Sauvignon | North America | unkown |
| I | ERR4588523 | USA | Cabernet Sauvignon | North America | unkown |
| I | ERR4588524 | USA | Zinfandel | North America | unkown |
| I | ERR4588525 | USA | Zinfandel | North America | unkown |
| I | ERR4588526 | USA | Cabernet Sauvignon | North America | unkown |
| I | ERR4588527 | USA | Sauvignon Blanc | North America | unkown |
| I | ERR4588528 | USA | Cabernet Sauvignon | North America | unkown |
| I | ERR4588529 | USA | Cabernet Sauvignon | North America | unkown |
| I | ERR4588530 | USA | Cabernet Sauvignon | North America | unkown |
| I | ERR4588531 | USA | Cabernet Sauvignon | North America | unkown |
| I | ERR4588532 | USA | Cabernet Franc | North America | unkown |
| I | ERR4588533 | USA | Cabernet Sauvignon | North America | unkown |
| I | ERR4588534 | USA | Cabernet Sauvignon | North America | unkown |
| I | ERR4588535 | USA | Cabernet Sauvignon | North America | unkown |

| | | | | | |
|---|---|---|---|---|---|
| I | ERR4588536 | USA | Cabernet Franc | North America | unkown |
| I | ERR4588537 | USA | Cabernet Sauvignon | North America | unkown |
| I | ERR4588538 | USA | Cabernet Sauvignon | North America | unkown |
| I | ERR4588539 | USA | Cabernet Sauvignon | North America | unkown |
| I | ERR4588540 | USA | Cabernet Sauvignon | North America | unkown |
| I | ERR4588541 | USA | Petit Verdot | North America | unkown |
| I | ERR4588542 | USA | Cabernet Sauvignon | North America | unkown |
| I | ERR4588543 | USA | Cabernet Franc | North America | unkown |
| I | ERR4588544 | USA | Chardonnay | North America | unkown |
| I | ERR4588545 | USA | Chardonnay | North America | unkown |
| I | ERR4588546 | USA | Pinot Noir | North America | unkown |
| I | ERR4588547 | USA | Pinot Noir | North America | unkown |
| I | ERR4588548 | USA | Chardonnay | North America | unkown |
| I | ERR4588549 | USA | Chardonnay | North America | unkown |
| I | ERR4588550 | USA | Chardonnay | North America | unkown |
| I | ERR4588551 | USA | Pinot Noir | North America | unkown |
| I | ERR4588552 | USA | Chardonnay | North America | unkown |
| I | ERR4588553 | USA | Pinot Noir | North America | unkown |
| I | ERR4588554 | USA | Cabernet Sauvignon | North America | unkown |
| I | ERR4588555 | USA | Pinot Noir | North America | unkown |
| I | ERR4588556 | USA | Pinot Noir | North America | unkown |
| I | ERR4588557 | USA | Cabernet Sauvignon | North America | unkown |
| I | ERR4588558 | USA | Cabernet Sauvignon | North America | unkown |
| I | ERR4588559 | USA | Cabernet Sauvignon | North America | unkown |
| I | ERR4588560 | USA | Cabernet Sauvignon | North America | unkown |
| I | ERR4588561 | USA | Cabernet Sauvignon | North America | unkown |
| I | ERR4588562 | Croatia | Silvaner | Europe | unkown |
| I | ERR4588563 | Croatia | Kraljevina | Europe | unkown |
| I | ERR4588564 | Croatia | Pinot Noir | Europe | unkown |
| I | ERR4588565 | France | Grenache | Europe | unkown |
| I | ERR4588566 | France | Grenache | Europe | unkown |
| I | ERR4588567 | France | Grenache | Europe | unkown |
| I | ERR4588568 | France | Grenache | Europe | unkown |
| I | ERR4588569 | Hungary | Other | Europe | unkown |
| I | ERR4588570 | Hungary | Blaufr√É¬§nkisch | Europe | unkown |
| I | ERR4588571 | Hungary | Other | Europe | unkown |
| I | ERR4588572 | Italy | Other | Europe | unkown |
| I | ERR4588573 | Spain | Treixadura | Europe | unkown |
| I | ERR4588574 | Spain | Palomino | Europe | unkown |
| I | ERR4588575 | Spain | Grenache | Europe | unkown |
| I | ERR4588576 | Spain | Mencia | Europe | unkown |
| I | ERR4588577 | USA | Other | North America | unkown |
| I | ERR4588578 | USA | Other | North America | unkown |
| I | ERR4588579 | Spain | Tempranillo | Europe | unkown |

| | | | | | |
|---|---|---|---|---|---|
| I | ERR4588580 | Spain | Tempranillo | Europe | unkown |
| I | ERR4588581 | Spain | Tempranillo | Europe | unkown |
| I | ERR4588582 | Spain | Tempranillo | Europe | unkown |
| I | ERR4588583 | Spain | Tempranillo | Europe | unkown |
| I | ERR4588584 | Spain | Tempranillo | Europe | unkown |
| I | ERR4588585 | Spain | Tempranillo | Europe | unkown |
| I | ERR4588586 | Spain | Tempranillo | Europe | unkown |
| I | ERR4588587 | Spain | Tempranillo | Europe | unkown |
| I | ERR4588588 | Spain | Tempranillo | Europe | unkown |
| I | ERR4588589 | Spain | Tempranillo | Europe | unkown |
| I | ERR4588590 | Spain | Merlot | Europe | unkown |
| I | ERR4588591 | Argentina | Chardonnay | South America | unkown |
| I | ERR4588592 | Argentina | Chardonnay | South America | unkown |
| I | ERR4588593 | Argentina | Chardonnay | South America | unkown |
| I | ERR4588594 | Spain | Tempranillo | Europe | unkown |
| I | ERR4588595 | Spain | Tempranillo | Europe | unkown |
| I | ERR4588596 | Spain | Tempranillo | Europe | unkown |
| I | ERR4588597 | Spain | Tempranillo | Europe | unkown |
| I | ERR4588598 | Spain | Tempranillo | Europe | unkown |
| I | ERR4588599 | Spain | Tempranillo | Europe | unkown |
| I | ERR4588600 | Spain | Tempranillo | Europe | unkown |
| I | ERR4588601 | Spain | Tempranillo | Europe | unkown |
| I | ERR4588602 | Spain | Tempranillo | Europe | unkown |
| I | ERR4588603 | Spain | Tempranillo | Europe | unkown |
| I | ERR4588604 | Spain | Tempranillo | Europe | unkown |
| I | ERR4588605 | Spain | Verdejo | Europe | unkown |
| I | ERR4588606 | Spain | Verdejo | Europe | unkown |
| I | ERR4588607 | Spain | Listan Negro | Europe | unkown |
| I | ERR4588608 | Spain | Tempranillo | Europe | unkown |
| I | ERR4588609 | Spain | Tempranillo | Europe | unkown |
| I | ERR4588610 | Spain | Tempranillo | Europe | unkown |
| I | ERR4588611 | Spain | Tempranillo | Europe | unkown |
| I | ERR4588612 | Spain | Airen | Europe | unkown |
| I | ERR4588613 | Spain | Airen | Europe | unkown |
| I | ERR4588614 | Spain | Airen | Europe | unkown |
| I | ERR4588615 | Spain | Airen | Europe | unkown |
| I | ERR4588616 | Spain | Airen | Europe | unkown |
| I | ERR4588617 | Spain | Carignan | Europe | unkown |
| I | ERR4588618 | Spain | Shiraz | Europe | unkown |
| I | ERR4588619 | Spain | Viognier | Europe | unkown |
| I | ERR4588620 | Spain | Shiraz | Europe | unkown |
| I | ERR4588621 | Spain | Shiraz | Europe | unkown |
| I | ERR4588622 | Spain | Shiraz | Europe | unkown |
| I | ERR4588623 | Spain | Merlot | Europe | unkown |

| | | | | | |
|---|---|---|---|---|---|
| I | ERR4588624 | Italy | Sauvignon Blanc | Europe | unkown |
| I | ERR4588625 | Italy | Sauvignon Blanc | Europe | unkown |
| I | ERR4588626 | Italy | Sauvignon Blanc | Europe | unkown |
| I | ERR4588627 | Italy | Barbera | Europe | unkown |
| I | ERR4588628 | Italy | Barbera | Europe | unkown |
| I | ERR4588629 | Italy | Barbera | Europe | unkown |
| I | ERR4588630 | USA | Grenache | North America | unkown |
| I | ERR4588631 | USA | Pinot Noir | North America | unkown |
| I | ERR4588632 | USA | Pinot Noir | North America | unkown |
| I | ERR4588633 | USA | Chardonnay | North America | unkown |
| I | ERR4588634 | USA | Pinot Noir | North America | unkown |
| I | ERR4588635 | USA | Cabernet Sauvignon | North America | unkown |
| I | ERR4588636 | USA | Cabernet Sauvignon | North America | unkown |
| I | ERR4588637 | USA | Cabernet Sauvignon | North America | unkown |
| I | ERR4588638 | USA | Cabernet Sauvignon | North America | unkown |
| I | ERR4588639 | USA | Cabernet Sauvignon | North America | unkown |
| I | ERR4588640 | USA | Cabernet Sauvignon | North America | unkown |
| I | ERR4588641 | USA | Cabernet Sauvignon | North America | unkown |
| I | ERR4588642 | USA | Cabernet Sauvignon | North America | unkown |
| I | ERR4588643 | USA | Cabernet Sauvignon | North America | unkown |
| I | ERR4588644 | USA | Cabernet Sauvignon | North America | unkown |
| I | ERR4588645 | USA | Cabernet Franc | North America | unkown |
| I | ERR4588646 | USA | Cabernet Franc | North America | unkown |
| I | ERR4588647 | USA | Petit Verdot | North America | unkown |
| I | ERR4588648 | USA | Montepulciano | North America | unkown |
| I | ERR4588649 | USA | Montepulciano | North America | unkown |
| I | ERR4588650 | Italy | Shiraz | Europe | unkown |
| I | ERR4588651 | USA | Cabernet Sauvignon | North America | unkown |
| I | ERR4588652 | Spain | Cabernet Sauvignon | Europe | unkown |
| I | ERR4588653 | Spain | Cabernet Sauvignon | Europe | unkown |
| I | ERR4588654 | Spain | Cabernet Sauvignon | Europe | unkown |
| I | ERR4588655 | Spain | Cabernet Sauvignon | Europe | unkown |
| I | ERR4588656 | Spain | Cabernet Sauvignon | Europe | unkown |
| I | ERR4588657 | Spain | Cabernet Sauvignon | Europe | unkown |
| I | ERR4588658 | Spain | Cabernet Sauvignon | Europe | unkown |
| I | ERR4588659 | Spain | Rufete | Europe | unkown |
| I | ERR4588660 | Spain | Rufete | Europe | unkown |
| I | ERR4588661 | Spain | Sauvignon Blanc | Europe | unkown |
| I | ERR4588662 | Spain | Tempranillo | Europe | unkown |
| I | ERR4588663 | Spain | Tempranillo | Europe | unkown |
| I | ERR4588664 | Spain | Tempranillo | Europe | unkown |
| I | ERR4588665 | Spain | Tempranillo | Europe | unkown |
| I | ERR4588666 | Spain | Tempranillo | Europe | unkown |
| I | ERR4588667 | Spain | Tempranillo | Europe | unkown |

| | | | | | |
|---|---|---|---|---|---|
| I | ERR4588668 | Spain | Graciano | Europe | unkown |
| I | ERR4588669 | Spain | Tempranillo | Europe | unkown |
| I | ERR4588670 | Spain | Tempranillo | Europe | unkown |
| I | ERR4588671 | Spain | Tempranillo | Europe | unkown |
| I | ERR4588672 | Spain | Tempranillo | Europe | unkown |
| I | ERR4588673 | Spain | Tempranillo | Europe | unkown |
| I | ERR4588674 | Spain | Tempranillo | Europe | unkown |
| I | ERR4588741 | South Africa | Shiraz | Africa | unkown |
| I | ERR4588742 | South Africa | Cabernet Sauvignon | Africa | unkown |
| I | ERR4588743 | South Africa | Shiraz | Africa | unkown |
| I | ERR4588744 | Chile | Aspirant Bouchet | South America | unkown |
| I | ERR4588745 | USA | Marechal Foch | North America | unkown |
| I | ERR4588675 | Australia | Shiraz | Oceania | unkown |
| I | ERR4588676 | Australia | Shiraz | Oceania | unkown |
| I | ERR4588677 | Australia | Shiraz | Oceania | unkown |
| I | ERR4588678 | Australia | Shiraz | Oceania | unkown |
| I | ERR4588679 | Australia | Cabernet Sauvignon | Oceania | unkown |
| I | ERR4588680 | Australia | Cabernet Sauvignon | Oceania | unkown |
| I | ERR4588681 | Australia | Cabernet Sauvignon | Oceania | unkown |
| I | ERR4588682 | Australia | Cabernet Sauvignon | Oceania | unkown |
| I | ERR4588683 | Australia | Cabernet Sauvignon | Oceania | unkown |
| I | ERR4588684 | Australia | Cabernet Sauvignon | Oceania | unkown |
| I | ERR4588685 | Australia | Cabernet Sauvignon | Oceania | unkown |
| I | ERR4588686 | Australia | Cabernet Sauvignon | Oceania | unkown |
| I | ERR4588687 | Australia | Cabernet Sauvignon | Oceania | unkown |
| I | ERR4588688 | Australia | Cabernet Sauvignon | Oceania | unkown |
| I | ERR4588689 | Australia | Malbec | Oceania | unkown |
| I | ERR4588690 | Australia | Malbec | Oceania | unkown |
| I | ERR4588691 | Australia | Malbec | Oceania | unkown |
| I | ERR4588692 | Australia | Cabernet Sauvignon | Oceania | unkown |
| I | ERR4588693 | Australia | Cabernet Sauvignon | Oceania | unkown |
| I | ERR4588694 | Australia | Cabernet Sauvignon | Oceania | unkown |
| I | ERR4588695 | Australia | Cabernet Sauvignon | Oceania | unkown |
| I | ERR4588696 | Australia | Cabernet Sauvignon | Oceania | unkown |
| I | ERR4588697 | Denmark | Rondo | Europe | unkown |
| I | ERR4588698 | Denmark | Rondo | Europe | unkown |
| I | ERR4588699 | Denmark | Rondo | Europe | unkown |
| I | ERR4588700 | Denmark | Rondo | Europe | unkown |
| I | ERR4588701 | Denmark | Rondo | Europe | unkown |
| I | ERR4588702 | Denmark | Solaris | Europe | unkown |
| I | ERR4588703 | Denmark | Solaris | Europe | unkown |
| I | ERR4588704 | Denmark | Solaris | Europe | unkown |
| I | ERR4588705 | Denmark | Solaris | Europe | unkown |
| I | ERR4588706 | Denmark | Solaris | Europe | unkown |

| | | | | | |
|---|---|---|---|---|---|
| I | ERR4588707 | Portugal | Verdelho | Europe | unkown |
| I | ERR4588708 | Portugal | Verdelho | Europe | unkown |
| I | ERR4588709 | Portugal | Verdelho | Europe | unkown |
| I | ERR4588710 | Portugal | Verdelho | Europe | unkown |
| I | ERR4588711 | Portugal | Verdelho | Europe | unkown |
| I | ERR4588712 | Portugal | Verdelho | Europe | unkown |
| I | ERR4588713 | Portugal | Verdelho | Europe | unkown |
| I | ERR4588714 | Portugal | Verdelho | Europe | unkown |
| I | ERR4588715 | Portugal | Verdelho | Europe | unkown |
| I | ERR4588716 | Denmark | Solaris | Europe | unkown |
| I | ERR4588717 | Denmark | Solaris | Europe | unkown |
| I | ERR4588718 | Denmark | Solaris | Europe | unkown |
| I | ERR4588719 | Denmark | Solaris | Europe | unkown |
| I | ERR4588720 | Denmark | Solaris | Europe | unkown |
| I | ERR4588721 | Australia | Cabernet Sauvignon | Oceania | unkown |
| I | ERR4588722 | Australia | Cabernet Sauvignon | Oceania | unkown |
| I | ERR4588723 | Australia | Cabernet Sauvignon | Oceania | unkown |
| I | ERR4588724 | Australia | Cabernet Sauvignon | Oceania | unkown |
| I | ERR4588725 | Australia | Cabernet Sauvignon | Oceania | unkown |
| I | ERR4588726 | Australia | Cabernet Sauvignon | Oceania | unkown |
| I | ERR4588727 | Australia | Cabernet Sauvignon | Oceania | unkown |
| I | ERR4588728 | Australia | Cabernet Sauvignon | Oceania | unkown |
| I | ERR4588729 | Australia | Cabernet Sauvignon | Oceania | unkown |
| I | ERR4588730 | Australia | Shiraz | Oceania | unkown |
| I | ERR4588731 | Germany | Riesling | Europe | unkown |
| I | ERR4588732 | Germany | Riesling | Europe | unkown |
| I | ERR4588733 | Germany | Riesling | Europe | unkown |
| I | ERR4588734 | Germany | Riesling | Europe | unkown |
| I | ERR4588735 | Germany | Riesling | Europe | unkown |
| I | ERR4588736 | Germany | Riesling | Europe | unkown |
| I | ERR4588737 | Germany | Riesling | Europe | unkown |
| I | ERR4588738 | Germany | Riesling | Europe | unkown |
| I | ERR4588739 | Germany | Riesling | Europe | unkown |
| I | ERR4588740 | Germany | Riesling | Europe | unkown |
| II | SRR10917592 | Australia | Shiraz | Oceania | Shiraz |
| II | SRR10917593 | Australia | Shiraz | Oceania | Shiraz |
| II | SRR10917594 | Australia | Shiraz | Oceania | Shiraz |
| II | SRR10917595 | Australia | Shiraz | Oceania | Shiraz |
| II | SRR10917596 | Australia | Shiraz | Oceania | Shiraz |
| II | SRR10917597 | Australia | Shiraz | Oceania | Shiraz |
| II | SRR10917598 | Australia | Shiraz | Oceania | Shiraz |
| II | SRR10917599 | Australia | Shiraz | Oceania | Shiraz |
| II | SRR10917600 | Australia | Shiraz | Oceania | Shiraz |
| II | SRR10917601 | Australia | Shiraz | Oceania | Shiraz |

| | | | | | |
|---|---|---|---|---|---|
| II | SRR10917583 | Australia | Shiraz | Oceania | Shiraz |
| II | SRR10917602 | Australia | Shiraz | Oceania | Shiraz |
| II | SRR10917603 | Australia | Shiraz | Oceania | Shiraz |
| II | SRR10917604 | Australia | Shiraz | Oceania | Shiraz |
| II | SRR10917605 | Australia | Shiraz | Oceania | Shiraz |
| II | SRR10917606 | Australia | Shiraz | Oceania | Shiraz |
| II | SRR10917607 | Australia | Shiraz | Oceania | Shiraz |
| II | SRR10917608 | Australia | Shiraz | Oceania | Shiraz |
| II | SRR10917609 | Australia | Shiraz | Oceania | Shiraz |
| II | SRR10917610 | Australia | Shiraz | Oceania | Shiraz |
| II | SRR10917611 | Australia | Shiraz | Oceania | Shiraz |
| II | SRR10917584 | Australia | Shiraz | Oceania | Shiraz |
| II | SRR10917612 | Australia | Shiraz | Oceania | Shiraz |
| II | SRR10917613 | Australia | Shiraz | Oceania | Shiraz |
| II | SRR10917614 | Australia | Shiraz | Oceania | Shiraz |
| II | SRR10917615 | Australia | Shiraz | Oceania | Shiraz |
| II | SRR10917616 | Australia | Shiraz | Oceania | Shiraz |
| II | SRR10917617 | Australia | Shiraz | Oceania | Shiraz |
| II | SRR10917618 | Australia | Shiraz | Oceania | Shiraz |
| II | SRR10917619 | Australia | Shiraz | Oceania | Shiraz |
| II | SRR10917620 | Australia | Shiraz | Oceania | Shiraz |
| II | SRR10917621 | Australia | Shiraz | Oceania | Shiraz |
| II | SRR10917585 | Australia | Shiraz | Oceania | Shiraz |
| II | SRR10917622 | Australia | Shiraz | Oceania | Shiraz |
| II | SRR10917623 | Australia | Shiraz | Oceania | Shiraz |
| II | SRR10917624 | Australia | Shiraz | Oceania | Shiraz |
| II | SRR10917625 | Australia | Shiraz | Oceania | Shiraz |
| II | SRR10917626 | Australia | Shiraz | Oceania | Shiraz |
| II | SRR10917627 | Australia | Shiraz | Oceania | Shiraz |
| II | SRR10917628 | Australia | Shiraz | Oceania | Shiraz |
| II | SRR10917629 | Australia | Shiraz | Oceania | Shiraz |
| II | SRR10917630 | Australia | Shiraz | Oceania | Shiraz |
| II | SRR10917631 | Australia | Shiraz | Oceania | Shiraz |
| II | SRR10917632 | Australia | Shiraz | Oceania | Shiraz |
| II | SRR10917633 | Australia | Shiraz | Oceania | Shiraz |
| II | SRR10917634 | Australia | Shiraz | Oceania | Shiraz |
| II | SRR10917635 | Australia | Shiraz | Oceania | Shiraz |
| II | SRR10917636 | Australia | Shiraz | Oceania | Shiraz |
| II | SRR10917637 | Australia | Shiraz | Oceania | Shiraz |
| II | SRR10917638 | Australia | Shiraz | Oceania | Shiraz |
| II | SRR10917639 | Australia | Shiraz | Oceania | Shiraz |
| II | SRR10917640 | Australia | Shiraz | Oceania | Shiraz |
| II | SRR10917641 | Australia | Shiraz | Oceania | Shiraz |
| II | SRR10917587 | Australia | Shiraz | Oceania | Shiraz |

| | | | | | |
|---|---|---|---|---|---|
| II | SRR10917642 | Australia | Shiraz | Oceania | Shiraz |
| II | SRR10917643 | Australia | Shiraz | Oceania | Shiraz |
| II | SRR10917644 | Australia | Shiraz | Oceania | Shiraz |
| II | SRR10917645 | Australia | Shiraz | Oceania | Shiraz |
| II | SRR10917646 | Australia | Shiraz | Oceania | Shiraz |
| II | SRR10917647 | Australia | Shiraz | Oceania | Shiraz |
| II | SRR10917648 | Australia | Shiraz | Oceania | Shiraz |
| II | SRR10917649 | Australia | Shiraz | Oceania | Shiraz |
| II | SRR10917588 | Australia | Shiraz | Oceania | Shiraz |
| II | SRR10917589 | Australia | Shiraz | Oceania | Shiraz |
| II | SRR10917590 | Australia | Shiraz | Oceania | Shiraz |
| II | SRR10917591 | Australia | Shiraz | Oceania | Shiraz |
| III | ERR557124 | USA | Merlot | North America | 3309C |
| III | ERR557125 | USA | Merlot | North America | 3309C |
| III | ERR557126 | USA | Merlot | North America | 3309C |
| III | ERR557127 | USA | Merlot | North America | 3309C |
| III | ERR557128 | USA | Merlot | North America | 3309C |
| III | ERR557129 | USA | Merlot | North America | 3309C |
| III | ERR557132 | USA | Merlot | North America | 3309C |
| III | ERR557135 | USA | Merlot | North America | 3309C |
| III | ERR557137 | France | Merlot | Europe | 3309C |
| III | ERR557142 | USA | Merlot | North America | 3309C |
| III | ERR557145 | France | Merlot | Europe | 3309C |
| III | ERR557146 | USA | Merlot | North America | 3309C |
| III | ERR557147 | USA | Merlot | North America | 3309C |
| III | ERR557149 | USA | Merlot | North America | 3309C |
| III | ERR557151 | USA | Merlot | North America | 3309C |
| III | ERR557152 | USA | Merlot | North America | 3309C |
| III | ERR557153 | USA | Merlot | North America | 3309C |
| III | ERR557155 | USA | Merlot | North America | 3309C |
| III | ERR557156 | USA | Merlot | North America | 3309C |
| III | ERR557157 | USA | Merlot | North America | 3309C |
| III | ERR557159 | USA | Merlot | North America | 3309C |
| III | ERR557160 | USA | Merlot | North America | 3309C |
| III | ERR557161 | USA | Merlot | North America | 3309C |
| III | ERR557163 | USA | Merlot | North America | 3309C |
| III | ERR557164 | USA | Merlot | North America | 3309C |
| III | ERR557168 | USA | Merlot | North America | 3309C |
| III | ERR557170 | USA | Merlot | North America | 3309C |
| III | ERR557171 | USA | Merlot | North America | 3309C |
| III | ERR557177 | USA | Merlot | North America | 3309C |
| III | ERR557178 | USA | Merlot | North America | 3309C |
| III | ERR557179 | USA | Merlot | North America | 3309C |
| III | ERR557181 | USA | Merlot | North America | 3309C |

| | | | | | |
|---|---|---|---|---|---|
| III | ERR557182 | USA | Merlot | North America | 3309C |
| III | ERR557183 | USA | Merlot | North America | 3309C |
| III | ERR557184 | USA | Merlot | North America | 3309C |
| III | ERR557185 | USA | Merlot | North America | 3309C |
| III | ERR557186 | USA | Merlot | North America | 3309C |
| III | ERR557188 | USA | Merlot | North America | 3309C |
| III | ERR557189 | France | Merlot | Europe | 3309C |
| III | ERR557192 | USA | Merlot | North America | 3309C |
| III | ERR557193 | France | Merlot | Europe | 3309C |
| III | ERR557194 | USA | Merlot | North America | 3309C |
| III | ERR557195 | USA | Merlot | North America | 3309C |
| III | ERR557196 | France | Merlot | Europe | 3309C |
| III | ERR557197 | USA | Merlot | North America | 3309C |
| III | ERR557198 | USA | Merlot | North America | 3309C |
| III | ERR557200 | USA | Merlot | North America | 3309C |
| III | ERR557201 | USA | Merlot | North America | 3309C |
| III | ERR557202 | USA | Merlot | North America | 3309C |
| III | ERR557203 | USA | Merlot | North America | 3309C |
| III | ERR557204 | USA | Merlot | North America | 3309C |
| III | ERR557208 | USA | Merlot | North America | 3309C |
| III | ERR557209 | USA | Merlot | North America | 3309C |
| III | ERR557211 | USA | Merlot | North America | 3309C |
| III | ERR557213 | France | Merlot | Europe | 3309C |
| III | ERR557219 | USA | Merlot | North America | 3309C |
| III | ERR557221 | USA | Merlot | North America | 3309C |
| III | ERR557222 | USA | Merlot | North America | 3309C |
| III | ERR557223 | USA | Merlot | North America | 3309C |
| III | ERR557225 | USA | Merlot | North America | 3309C |
| III | ERR557226 | USA | Merlot | North America | 3309C |
| III | ERR557227 | USA | Merlot | North America | 3309C |
| III | ERR557228 | USA | Merlot | North America | 3309C |
| III | ERR557230 | USA | Merlot | North America | 3309C |
| III | ERR557232 | USA | Merlot | North America | 3309C |
| III | ERR557233 | USA | Merlot | North America | 3309C |
| III | ERR557236 | USA | Merlot | North America | 3309C |
| III | ERR557237 | USA | Merlot | North America | 3309C |
| III | ERR557238 | USA | Merlot | North America | 3309C |
| III | ERR557239 | USA | Merlot | North America | 3309C |
| III | ERR557240 | USA | Merlot | North America | 3309C |
| III | ERR557241 | USA | Merlot | North America | 3309C |
| III | ERR557242 | USA | Merlot | North America | 3309C |
| III | ERR557244 | USA | Merlot | North America | 3309C |
| III | ERR557248 | USA | Merlot | North America | 3309C |
| III | ERR557249 | USA | Merlot | North America | 3309C |

| | | | | | |
|---|---|---|---|---|---|
| III | ERR557250 | USA | Merlot | North America | 3309C |
| III | ERR557251 | USA | Merlot | North America | 3309C |
| III | ERR557252 | France | Merlot | Europe | 3309C |
| III | ERR557254 | USA | Merlot | North America | 3309C |
| III | ERR557259 | USA | Merlot | North America | 3309C |
| III | ERR557260 | USA | Merlot | North America | 3309C |
| III | ERR557266 | USA | Merlot | North America | 3309C |
| III | ERR557267 | USA | Merlot | North America | 3309C |
| III | ERR557268 | USA | Merlot | North America | 3309C |
| III | ERR557270 | USA | Merlot | North America | 3309C |
| III | ERR557272 | USA | Merlot | North America | 3309C |
| III | ERR557273 | USA | Merlot | North America | 3309C |
| III | ERR557275 | USA | Merlot | North America | 3309C |
| III | ERR557278 | USA | Merlot | North America | 3309C |
| III | ERR557279 | USA | Merlot | North America | 3309C |
| III | ERR557280 | USA | Merlot | North America | 3309C |
| III | ERR557281 | France | Merlot | Europe | 3309C |
| III | ERR557284 | USA | Merlot | North America | 3309C |
| III | ERR557285 | USA | Merlot | North America | 3309C |
| III | ERR557286 | USA | Merlot | North America | 3309C |
| III | ERR557287 | USA | Merlot | North America | 3309C |
| III | ERR557289 | USA | Merlot | North America | 3309C |
| III | ERR557291 | France | Merlot | Europe | 3309C |
| III | ERR557296 | USA | Merlot | North America | 3309C |
| III | ERR557297 | USA | Merlot | North America | 3309C |
| III | ERR557298 | USA | Merlot | North America | 3309C |
| III | ERR557300 | USA | Merlot | North America | 3309C |
| III | ERR557301 | USA | Merlot | North America | 3309C |
| III | ERR557302 | USA | Merlot | North America | 3309C |
| III | ERR557303 | USA | Merlot | North America | 3309C |
| III | ERR557304 | USA | Merlot | North America | 3309C |
| III | ERR557306 | USA | Merlot | North America | 3309C |
| III | ERR557309 | France | Merlot | Europe | 3309C |
| III | ERR557310 | USA | Merlot | North America | 3309C |
| III | ERR557311 | USA | Merlot | North America | 3309C |
| III | ERR557313 | France | Merlot | Europe | 3309C |
| III | ERR557314 | USA | Merlot | North America | 3309C |
| III | ERR557315 | USA | Merlot | North America | 3309C |
| III | ERR557316 | USA | Merlot | North America | 3309C |
| III | ERR557317 | USA | Merlot | North America | 3309C |
| III | ERR557319 | USA | Merlot | North America | 3309C |
| III | ERR557321 | USA | Merlot | North America | 3309C |
| III | ERR557324 | USA | Merlot | North America | 3309C |
| III | ERR557326 | USA | Merlot | North America | 3309C |

| | | | | | |
|---|---|---|---|---|---|
| III | ERR557328 | USA | Merlot | North America | 3309C |
| III | ERR557329 | USA | Merlot | North America | 3309C |
| III | ERR557330 | USA | Merlot | North America | 3309C |
| III | ERR557331 | USA | Merlot | North America | 3309C |
| III | ERR557333 | USA | Merlot | North America | 3309C |
| III | ERR557336 | USA | Merlot | North America | 3309C |
| III | ERR557337 | France | Merlot | Europe | 3309C |
| III | ERR557338 | USA | Merlot | North America | 3309C |
| III | ERR557339 | USA | Merlot | North America | 3309C |
| III | ERR557340 | USA | Merlot | North America | 3309C |
| III | ERR557343 | USA | Merlot | North America | 3309C |
| III | ERR557344 | USA | Merlot | North America | 3309C |
| III | ERR557345 | USA | Merlot | North America | 3309C |
| III | ERR557346 | USA | Merlot | North America | 3309C |
| III | ERR557348 | USA | Merlot | North America | 3309C |
| III | ERR557349 | USA | Merlot | North America | 3309C |
| III | ERR557350 | USA | Merlot | North America | 3309C |
| III | ERR557352 | USA | Merlot | North America | 3309C |
| III | ERR557353 | USA | Merlot | North America | 3309C |
| III | ERR557354 | USA | Merlot | North America | 3309C |
| III | ERR557355 | USA | Merlot | North America | 3309C |
| III | ERR557356 | USA | Merlot | North America | 3309C |
| III | ERR557359 | USA | Merlot | North America | 3309C |
| III | ERR557360 | USA | Merlot | North America | 3309C |
| III | ERR557365 | USA | Merlot | North America | 3309C |
| III | ERR557366 | USA | Merlot | North America | 3309C |
| III | ERR557367 | USA | Merlot | North America | 3309C |
| III | ERR557369 | USA | Merlot | North America | 3309C |
| III | ERR557370 | USA | Merlot | North America | 3309C |
| III | ERR557371 | USA | Merlot | North America | 3309C |
| III | ERR557372 | USA | Merlot | North America | 3309C |
| III | ERR557373 | USA | Merlot | North America | 3309C |
| III | ERR557376 | USA | Merlot | North America | 3309C |
| III | ERR557377 | USA | Merlot | North America | 3309C |
| III | ERR557379 | France | Merlot | Europe | 3309C |
| III | ERR557380 | USA | Merlot | North America | 3309C |
| III | ERR557381 | USA | Merlot | North America | 3309C |
| III | ERR557382 | USA | Merlot | North America | 3309C |
| III | ERR557386 | USA | Merlot | North America | 3309C |
| III | ERR557388 | USA | Merlot | North America | 3309C |
| III | ERR557390 | USA | Merlot | North America | 3309C |
| III | ERR557391 | USA | Merlot | North America | 3309C |
| III | ERR557392 | USA | Merlot | North America | 3309C |
| III | ERR557396 | USA | Merlot | North America | 3309C |

| | | | | | |
|---|---|---|---|---|---|
| III | ERR557398 | USA | Merlot | North America | 3309C |
| III | ERR557399 | USA | Merlot | North America | 3309C |
| III | ERR557400 | France | Merlot | Europe | 3309C |
| III | ERR557401 | France | Merlot | Europe | 3309C |
| III | ERR557402 | USA | Merlot | North America | 3309C |
| III | ERR557403 | USA | Merlot | North America | 3309C |
| III | ERR557405 | France | Merlot | Europe | 3309C |
| III | ERR557406 | USA | Merlot | North America | 3309C |
| III | ERR557407 | USA | Merlot | North America | 3309C |
| III | ERR557409 | USA | Merlot | North America | 3309C |
| III | ERR557411 | USA | Merlot | North America | 3309C |
| III | ERR557412 | USA | Merlot | North America | 3309C |
| III | ERR557413 | USA | Merlot | North America | 3309C |
| III | ERR557414 | USA | Merlot | North America | 3309C |
| III | ERR557416 | USA | Merlot | North America | 3309C |
| III | ERR557421 | USA | Merlot | North America | 3309C |
| III | ERR557424 | USA | Merlot | North America | 3309C |
| III | ERR557425 | USA | Merlot | North America | 3309C |
| III | ERR557426 | USA | Merlot | North America | 3309C |
| III | ERR557428 | USA | Merlot | North America | 3309C |
| III | ERR557429 | USA | Merlot | North America | 3309C |
| III | ERR557430 | USA | Merlot | North America | 3309C |
| III | ERR557431 | USA | Merlot | North America | 3309C |
| III | ERR557432 | USA | Merlot | North America | 3309C |
| III | ERR557434 | USA | Merlot | North America | 3309C |
| III | ERR557435 | USA | Merlot | North America | 3309C |
| III | ERR557436 | USA | Merlot | North America | 3309C |
| III | ERR557437 | USA | Merlot | North America | 3309C |
| III | ERR557439 | USA | Merlot | North America | 3309C |
| III | ERR557440 | USA | Merlot | North America | 3309C |
| III | ERR557442 | USA | Merlot | North America | 3309C |
| III | ERR557444 | France | Merlot | Europe | 3309C |
| III | ERR557445 | USA | Merlot | North America | 3309C |
| III | ERR557446 | USA | Merlot | North America | 3309C |
| III | ERR557447 | France | Merlot | Europe | 3309C |
| III | ERR557451 | USA | Merlot | North America | 3309C |
| III | ERR557455 | USA | Merlot | North America | 3309C |
| III | ERR557457 | USA | Merlot | North America | 3309C |
| III | ERR557461 | USA | Merlot | North America | 3309C |
| III | ERR557462 | USA | Merlot | North America | 3309C |
| III | ERR557463 | USA | Merlot | North America | 3309C |
| III | ERR557464 | USA | Merlot | North America | 3309C |
| III | ERR557465 | USA | Merlot | North America | 3309C |
| III | ERR557466 | USA | Merlot | North America | 3309C |

| | | | | | |
|---|---|---|---|---|---|
| III | ERR557467 | USA | Merlot | North America | 3309C |
| III | ERR557468 | USA | Merlot | North America | 3309C |
| III | ERR557469 | USA | Merlot | North America | 3309C |
| III | ERR557470 | USA | Merlot | North America | 3309C |
| III | ERR557472 | USA | Merlot | North America | 3309C |
| III | ERR557473 | USA | Merlot | North America | 3309C |
| III | ERR557474 | USA | Merlot | North America | 3309C |
| III | ERR557475 | USA | Merlot | North America | 3309C |
| III | ERR557476 | USA | Merlot | North America | 3309C |
| III | ERR557477 | USA | Merlot | North America | 3309C |
| III | ERR557479 | USA | Merlot | North America | 3309C |
| III | ERR557481 | USA | Merlot | North America | 3309C |
| III | ERR557482 | USA | Merlot | North America | 3309C |
| III | ERR557483 | USA | Merlot | North America | 3309C |
| III | ERR557485 | USA | Merlot | North America | 3309C |
| III | ERR557486 | USA | Merlot | North America | 3309C |
| III | ERR557487 | USA | Merlot | North America | 3309C |
| III | ERR557488 | USA | Merlot | North America | 3309C |
| III | ERR557489 | USA | Merlot | North America | 3309C |
| III | ERR557490 | USA | Merlot | North America | 3309C |
| III | ERR557493 | USA | Merlot | North America | 3309C |
| III | ERR557494 | France | Merlot | Europe | 3309C |
| III | ERR557495 | USA | Merlot | North America | 3309C |
| III | ERR557497 | USA | Merlot | North America | 3309C |
| III | ERR557499 | USA | Merlot | North America | 3309C |
| III | ERR557500 | USA | Merlot | North America | 3309C |
| III | ERR557502 | USA | Merlot | North America | 3309C |
| III | ERR557503 | USA | Merlot | North America | 3309C |
| III | ERR557507 | USA | Merlot | North America | 3309C |
| III | ERR557510 | USA | Merlot | North America | 3309C |
| III | ERR557511 | USA | Merlot | North America | 3309C |
| III | ERR557512 | USA | Merlot | North America | 3309C |
| III | ERR557514 | USA | Merlot | North America | 3309C |
| III | ERR557518 | USA | Merlot | North America | 3309C |
| III | ERR557519 | USA | Merlot | North America | 3309C |
| III | ERR557520 | USA | Merlot | North America | 3309C |
| III | ERR557521 | USA | Merlot | North America | 3309C |
| III | ERR557522 | USA | Merlot | North America | 3309C |
| III | ERR557523 | USA | Merlot | North America | 3309C |
| III | 1024.DPOO1.C | USA | Merlot | North America | 3309C |
| III | 1024.DPOO1.C | USA | Merlot | North America | 3309C |
| III | 1024.DPOO1.C | USA | Merlot | North America | 3309C |
| III | 1024.DPOO1.C | USA | Merlot | North America | 3309C |
| III | 1024.DPOO1.C | USA | Merlot | North America | 3309C |

| | | | | | |
|---|---|---|---|---|---|
| III | 1024.DPOO1.C | USA | Merlot | North America | 3309C |
| III | 1024.DPOO1.C | USA | Merlot | North America | 3309C |
| III | 1024.DPOO1.C | USA | Merlot | North America | 3309C |
| III | 1024.DPOO1.C | USA | Merlot | North America | 3309C |
| III | 1024.DPOO1.C | USA | Merlot | North America | 3309C |
| III | 1024.DPOO1.C | USA | Merlot | North America | 3309C |
| III | 1024.DPOO1.C | USA | Merlot | North America | 3309C |
| III | 1024.DPOO1.C | USA | Merlot | North America | 3309C |
| III | 1024.DPOO1.C | USA | Merlot | North America | 3309C |
| III | 1024.DPOO1.C | USA | Merlot | North America | 3309C |
| III | 1024.DPOO1.C | USA | Merlot | North America | 3309C |
| III | 1024.DPOO1.C | USA | Merlot | North America | 3309C |
| III | 1024.DPOO1.C | USA | Merlot | North America | 3309C |
| III | 1024.DPOO1.C | USA | Merlot | North America | 3309C |
| III | 1024.DPOO1.C | USA | Merlot | North America | 3309C |
| III | 1024.DPOO1.C | USA | Merlot | North America | 3309C |
| III | 1024.DPOO1.C | USA | Merlot | North America | 3309C |
| III | 1024.DPOO1.C | USA | Merlot | North America | 3309C |
| III | 1024.DPOO1.C | USA | Merlot | North America | 3309C |
| III | 1024.DPOO1.C | USA | Merlot | North America | 3309C |
| III | 1024.HE003.C1 | USA | Merlot | North America | 3309C |
| III | 1024.HE003.C1 | USA | Merlot | North America | 3309C |
| III | 1024.HE003.C1 | USA | Merlot | North America | 3309C |
| III | 1024.HE003.C1 | USA | Merlot | North America | 3309C |
| III | 1024.HE003.C1 | USA | Merlot | North America | 3309C |
| III | 1024.HE003.C1 | USA | Merlot | North America | 3309C |
| III | 1024.HE003.C1 | USA | Merlot | North America | 3309C |
| III | 1024.HE003.C1 | USA | Merlot | North America | 3309C |
| III | 1024.HE003.C1 | USA | Merlot | North America | 3309C |
| III | 1024.HE003.C1 | USA | Merlot | North America | 3309C |
| III | 1024.HE003.C1 | USA | Merlot | North America | 3309C |
| III | 1024.HE003.C1 | USA | Merlot | North America | 3309C |
| III | 1024.HE003.C1 | USA | Merlot | North America | 3309C |
| III | 1024.HE003.C1 | USA | Merlot | North America | 3309C |
| III | 1024.HE003.C1 | USA | Merlot | North America | 3309C |
| III | 1024.HE003.C1 | USA | Merlot | North America | 3309C |
| III | 1024.HE003.C1 | USA | Merlot | North America | 3309C |
| III | 1024.HE003.C1 | USA | Merlot | North America | 3309C |
| III | 1024.HE003.C1 | USA | Merlot | North America | 3309C |
| III | 1024.HE003.C1 | USA | Merlot | North America | 3309C |
| III | 1024.HE003.C1 | USA | Merlot | North America | 3309C |
| III | 1024.HE003.C1 | USA | Merlot | North America | 3309C |
| III | 1024.MU002.C | USA | Merlot | North America | 3309C |
| III | 1024.MU002.C | USA | Merlot | North America | 3309C |

| | | | | | |
|---|---|---|---|---|---|
| III | 1024.MU002.C | USA | Merlot | North America | 3309C |
| III | 1024.MU002.C | USA | Merlot | North America | 3309C |
| III | 1024.MU002.C | USA | Merlot | North America | 3309C |
| III | 1024.MU002.C | USA | Merlot | North America | 3309C |
| III | 1024.MU002.C | USA | Merlot | North America | 3309C |
| III | 1024.MU002.C | USA | Merlot | North America | 3309C |
| III | 1024.MU002.C | USA | Merlot | North America | 3309C |
| III | 1024.MU002.C | USA | Merlot | North America | 3309C |
| III | 1024.MU002.C | USA | Merlot | North America | 3309C |
| III | 1024.MU002.C | USA | Merlot | North America | 3309C |
| III | 1024.MU002.C | USA | Merlot | North America | 3309C |
| III | 1024.MU002.C | USA | Merlot | North America | 3309C |
| III | 1024.MU002.C | USA | Merlot | North America | 3309C |
| III | 1024.MU002.C | USA | Merlot | North America | 3309C |
| III | 1024.MU002.C | USA | Merlot | North America | 3309C |
| III | 1024.MU002.C | USA | Merlot | North America | 3309C |
| III | 1024.MU002.C | USA | Merlot | North America | 3309C |
| III | 1024.MU002.C | USA | Merlot | North America | 3309C |
| III | 1024.MU002.C | USA | Merlot | North America | 3309C |
| III | 1024.MU002.C | USA | Merlot | North America | 3309C |
| III | 1024.MU002.C | USA | Merlot | North America | 3309C |
| III | 1024.MU002.C | USA | Merlot | North America | 3309C |
| III | 1024.MU002.C | USA | Merlot | North America | 3309C |
| III | 1024.RU006.C1 | USA | Merlot | North America | 3309C |
| III | 1024.RU006.C1 | USA | Merlot | North America | 3309C |
| III | 1024.RU006.C1 | USA | Merlot | North America | 3309C |
| III | 1024.RU006.C1 | USA | Merlot | North America | 3309C |
| III | 1024.RU006.C1 | USA | Merlot | North America | 3309C |
| III | 1024.RU006.C1 | USA | Merlot | North America | 3309C |
| III | 1024.RU006.C1 | USA | Merlot | North America | 3309C |
| III | 1024.RU006.C1 | USA | Merlot | North America | 3309C |
| III | 1024.RU006.C1 | USA | Merlot | North America | 3309C |
| III | 1024.RU006.C1 | USA | Merlot | North America | 3309C |
| III | 1024.RU006.C1 | USA | Merlot | North America | 3309C |
| III | 1024.RU006.C1 | USA | Merlot | North America | 3309C |
| III | 1024.RU006.C1 | USA | Merlot | North America | 3309C |
| III | 1024.RU006.C1 | USA | Merlot | North America | 3309C |
| III | 1024.RU006.C1 | USA | Merlot | North America | 3309C |
| III | 1024.RU006.C1 | USA | Merlot | North America | 3309C |
| III | 1024.RU006.C1 | USA | Merlot | North America | 3309C |
| III | 1024.RU006.C1 | USA | Merlot | North America | 3309C |
| III | 1024.RU006.C1 | USA | Merlot | North America | 3309C |
| III | 1024.RU006.C1 | USA | Merlot | North America | 3309C |
| III | 1024.RU006.C1 | USA | Merlot | North America | 3309C |

| | | | | | |
|---|---|---|---|---|---|
| III | 1024.RU006.C1 | USA | Merlot | North America | 3309C |
| III | 1024.RU006.C1 | USA | Merlot | North America | 3309C |
| III | 1024.SH004.C1 | USA | Merlot | North America | 3309C |
| III | 1024.SH004.C1 | USA | Merlot | North America | 3309C |
| III | 1024.SH004.C1 | USA | Merlot | North America | 3309C |
| III | 1024.SH004.C1 | USA | Merlot | North America | 3309C |
| III | 1024.SH004.C1 | USA | Merlot | North America | 3309C |
| III | 1024.SH004.C1 | USA | Merlot | North America | 3309C |
| III | 1024.SH004.C1 | USA | Merlot | North America | 3309C |
| III | 1024.SH004.C1 | USA | Merlot | North America | 3309C |
| III | 1024.SH004.C1 | USA | Merlot | North America | 3309C |
| III | 1024.SH004.C1 | USA | Merlot | North America | 3309C |
| III | 1024.SH004.C1 | USA | Merlot | North America | 3309C |
| III | 1024.SH004.C1 | USA | Merlot | North America | 3309C |
| III | 1024.SH004.C1 | USA | Merlot | North America | 3309C |
| III | 1024.SH004.C1 | USA | Merlot | North America | 3309C |
| III | 1024.SH004.C1 | USA | Merlot | North America | 3309C |
| III | 1024.SH004.C1 | USA | Merlot | North America | 3309C |
| III | 1024.SH004.C1 | USA | Merlot | North America | 3309C |
| III | 1024.SH004.C1 | USA | Merlot | North America | 3309C |
| III | 1024.SH004.C1 | USA | Merlot | North America | 3309C |
| III | 1024.SH004.C1 | USA | Merlot | North America | 3309C |
| III | 1024.SH005.C3 | USA | Merlot | North America | 3309C |
| III | 1024.SH005.C3 | USA | Merlot | North America | 3309C |
| III | 1024.SH005.C3 | USA | Merlot | North America | 3309C |
| III | 1024.SH005.C3 | USA | Merlot | North America | 3309C |
| III | 1024.SH005.C3 | USA | Merlot | North America | 3309C |
| III | 1024.SH005.C3 | USA | Merlot | North America | 3309C |
| III | 1024.SH005.C3 | USA | Merlot | North America | 3309C |
| III | 1024.SH005.C3 | USA | Merlot | North America | 3309C |
| III | 1024.SH005.C3 | USA | Merlot | North America | 3309C |
| III | 1024.SH005.C3 | USA | Merlot | North America | 3309C |
| III | 1024.SH005.C3 | USA | Merlot | North America | 3309C |
| III | 1024.SH005.C3 | USA | Merlot | North America | 3309C |
| III | 1024.SH005.C3 | USA | Merlot | North America | 3309C |
| III | 1024.SH005.C3 | USA | Merlot | North America | 3309C |
| III | 1024.SH005.C3 | USA | Merlot | North America | 3309C |
| III | 1024.SH005.C3 | USA | Merlot | North America | 3309C |
| III | 1024.SH005.C3 | USA | Merlot | North America | 3309C |
| III | 1024.SH005.C3 | USA | Merlot | North America | 3309C |
| III | 1024.SH005.C3 | USA | Merlot | North America | 3309C |
| III | 1024.SH007.C6 | USA | Merlot | North America | 3309C |
| III | 1024.SH007.C6 | USA | Merlot | North America | 3309C |
| III | 1024.SH007.C6 | USA | Merlot | North America | 3309C |

| | | | | | |
|---|---|---|---|---|---|
| III | 1024.SH007.C6 | USA | Merlot | North America | 3309C |
| III | 1024.SH007.C6 | USA | Merlot | North America | 3309C |
| III | 1024.SH007.C6 | USA | Merlot | North America | 3309C |
| III | 1024.SH007.C6 | USA | Merlot | North America | 3309C |
| III | 1024.SH007.C6 | USA | Merlot | North America | 3309C |
| III | 1024.SH007.C6 | USA | Merlot | North America | 3309C |
| III | 1024.SH007.C6 | USA | Merlot | North America | 3309C |
| III | 1024.SH007.C6 | USA | Merlot | North America | 3309C |
| III | 1024.SH007.C6 | USA | Merlot | North America | 3309C |
| III | 1024.SH007.C6 | USA | Merlot | North America | 3309C |
| III | 1024.SH007.C6 | USA | Merlot | North America | 3309C |
| III | 1024.SH007.C6 | USA | Merlot | North America | 3309C |
| III | 1024.SH007.C6 | USA | Merlot | North America | 3309C |
| III | 1024.SH007.C6 | USA | Merlot | North America | 3309C |
| III | 1024.SH007.C6 | USA | Merlot | North America | 3309C |
| III | 1024.SH007.C6 | USA | Merlot | North America | 3309C |
| III | 1024.SH007.C6 | USA | Merlot | North America | 3309C |
| III | 1024.SH008.C6 | USA | Merlot | North America | 3309C |
| III | 1024.SH008.C6 | USA | Merlot | North America | 3309C |
| III | 1024.SH008.C6 | USA | Merlot | North America | 3309C |
| III | 1024.SH008.C6 | USA | Merlot | North America | 3309C |
| III | 1024.SH008.C6 | USA | Merlot | North America | 3309C |
| III | 1024.SH008.C6 | USA | Merlot | North America | 3309C |
| III | 1024.SH008.C6 | USA | Merlot | North America | 3309C |
| III | 1024.SH008.C6 | USA | Merlot | North America | 3309C |
| III | 1024.SH008.C6 | USA | Merlot | North America | 3309C |
| III | 1024.SH008.C6 | USA | Merlot | North America | 3309C |
| III | 1024.SH008.C6 | USA | Merlot | North America | 3309C |
| III | 1024.SH008.C6 | USA | Merlot | North America | 3309C |
| III | 1024.SH008.C6 | USA | Merlot | North America | 3309C |
| III | 1024.SH008.C6 | USA | Merlot | North America | 3309C |
| III | 1024.SH008.C6 | USA | Merlot | North America | 3309C |
| III | 1024.SH008.C6 | USA | Merlot | North America | 3309C |
| III | 1024.SH008.C6 | USA | Merlot | North America | 3309C |
| III | 1024.SH008.C6 | USA | Merlot | North America | 3309C |
| III | 1024.SH008.C6 | USA | Merlot | North America | 3309C |
| III | 1024.SH008.C6 | USA | Merlot | North America | 3309C |
| III | 1024.SH008.C6 | USA | Merlot | North America | 3309C |
| III | 1024.SH008.C6 | USA | Merlot | North America | 3309C |
| III | 1024.SH008.C6 | USA | Merlot | North America | 3309C |
| III | 1024.SH008.C6 | USA | Merlot | North America | 3309C |
| III | 1024.SH008.C6 | USA | Merlot | North America | 3309C |
| IV | SRR18047940 | Italy | Chardonnay | Europe | 420A |
| IV | SRR18047941 | Italy | Chardonnay | Europe | 420A |

| | | | | | |
|---|---|---|---|---|---|
| IV | SRR18047942 | Italy | Chardonnay | Europe | 420A |
| IV | SRR18047943 | Italy | Chardonnay | Europe | 420A |
| IV | SRR18047944 | Italy | Chardonnay | Europe | 420A |
| IV | SRR18047945 | Italy | Chardonnay | Europe | 420A |
| IV | SRR18047946 | Italy | Chardonnay | Europe | 420A |
| IV | SRR18047947 | Italy | Chardonnay | Europe | 420A |
| IV | SRR18047948 | Italy | Chardonnay | Europe | 420A |
| IV | SRR18047949 | Italy | Chardonnay | Europe | 420A |
| IV | SRR18047950 | Italy | Chardonnay | Europe | 420A |
| IV | SRR18047951 | Italy | Chardonnay | Europe | 420A |
| IV | SRR18047952 | Italy | Chardonnay | Europe | 420A |
| IV | SRR18047953 | Italy | Chardonnay | Europe | 420A |
| IV | SRR18047954 | Italy | Chardonnay | Europe | 420A |
| IV | SRR18047955 | Italy | Chardonnay | Europe | 420A |
| IV | SRR18047956 | Italy | Sangiovese | Europe | 110R |
| IV | SRR18047957 | Italy | Sangiovese | Europe | 110R |
| IV | SRR18047958 | Italy | Sangiovese | Europe | 110R |
| IV | SRR18047959 | Italy | Sangiovese | Europe | 110R |
| IV | SRR18047960 | Italy | Sangiovese | Europe | 110R |
| IV | SRR18047961 | Italy | Sangiovese | Europe | 110R |
| IV | SRR18047962 | Italy | Merlot | Europe | 420A |
| IV | SRR18047963 | Italy | Merlot | Europe | 420A |
| IV | SRR18047964 | Italy | Merlot | Europe | 420A |
| IV | SRR18047965 | Italy | Merlot | Europe | 420A |
| IV | SRR18047966 | Italy | Sangiovese | Europe | 420A |
| IV | SRR18047967 | Italy | Merlot | Europe | 420A |
| IV | SRR18047968 | Italy | Merlot | Europe | 420A |
| IV | SRR18047969 | Italy | Merlot | Europe | 420A |
| IV | SRR18047970 | Italy | Merlot | Europe | 420A |
| IV | SRR18047971 | Italy | Merlot | Europe | 420A |
| IV | SRR18047972 | Italy | Merlot | Europe | 420A |
| IV | SRR18047973 | Italy | Merlot | Europe | 420A |
| IV | SRR18047974 | Italy | Merlot | Europe | 420A |
| IV | SRR18047975 | Italy | Sangiovese | Europe | 420A |
| IV | SRR18047976 | Italy | Sangiovese | Europe | 420A |
| IV | SRR18047977 | Italy | Sangiovese | Europe | 420A |
| IV | SRR18047978 | Italy | Sangiovese | Europe | 420A |
| IV | SRR18047979 | Italy | Sangiovese | Europe | 420A |
| IV | SRR18047980 | Italy | Sangiovese | Europe | 420A |
| IV | SRR18047981 | Italy | Sangiovese | Europe | 420A |
| IV | SRR18047982 | Italy | Chardonnay | Europe | 3309C |
| IV | SRR18047983 | Italy | Chardonnay | Europe | 3309C |
| IV | SRR18047984 | Italy | Chardonnay | Europe | 3309C |
| IV | SRR18047985 | Italy | Chardonnay | Europe | 3309C |

| | | | | | |
|---|---|---|---|---|---|
| IV | SRR18047986 | Italy | Chardonnay | Europe | 3309C |
| IV | SRR18047987 | Italy | Chardonnay | Europe | 3309C |
| IV | SRR18047988 | Italy | Sangiovese | Europe | 420A |
| IV | SRR18047989 | Italy | Chardonnay | Europe | 110R |
| IV | SRR18047990 | Italy | Chardonnay | Europe | 110R |
| IV | SRR18047991 | Italy | Chardonnay | Europe | 110R |
| IV | SRR18047992 | Italy | Chardonnay | Europe | 110R |
| IV | SRR18047993 | Italy | Chardonnay | Europe | 110R |
| IV | SRR18047994 | Italy | Chardonnay | Europe | 110R |
| IV | SRR18047995 | Italy | Chardonnay | Europe | SO4 |
| IV | SRR18047996 | Italy | Chardonnay | Europe | SO4 |
| IV | SRR18047997 | Italy | Chardonnay | Europe | SO4 |
| IV | SRR18047998 | Italy | Chardonnay | Europe | SO4 |
| IV | SRR18047999 | Italy | Sangiovese | Europe | 420A |
| IV | SRR18048000 | Italy | Chardonnay | Europe | SO4 |
| IV | SRR18048001 | Italy | Chardonnay | Europe | SO4 |
| IV | SRR18048002 | Italy | Merlot | Europe | 420A |
| IV | SRR18048003 | Italy | Merlot | Europe | 420A |
| IV | SRR18048004 | Italy | Merlot | Europe | 420A |
| IV | SRR18048005 | Italy | Merlot | Europe | 420A |
| IV | SRR18048006 | Italy | Merlot | Europe | 420A |
| IV | SRR18048007 | Italy | Merlot | Europe | 420A |
| IV | SRR18048008 | Italy | Chardonnay | Europe | 420A |
| IV | SRR18048009 | Italy | Chardonnay | Europe | 420A |
| IV | SRR18048010 | Italy | Sangiovese | Europe | 420A |
| IV | SRR18048011 | Italy | Sangiovese | Europe | 420A |
| IV | SRR18048234 | Italy | Merlot | Europe | 420A |
| IV | SRR18048235 | Italy | Merlot | Europe | 420A |
| IV | SRR18048236 | Italy | Merlot | Europe | 420A |
| IV | SRR18048237 | Italy | Sangiovese | Europe | 420A |
| IV | SRR18048238 | Italy | Sangiovese | Europe | 420A |
| IV | SRR18048239 | Italy | Sangiovese | Europe | 420A |
| IV | SRR18048240 | Italy | Sangiovese | Europe | 420A |
| IV | SRR18048241 | Italy | Sangiovese | Europe | 420A |
| IV | SRR18048242 | Italy | Sangiovese | Europe | 420A |
| IV | SRR18048243 | Italy | Sangiovese | Europe | 420A |
| IV | SRR18048244 | Italy | Chardonnay | Europe | 3309C |
| IV | SRR18048245 | Italy | Chardonnay | Europe | 3309C |
| IV | SRR18048246 | Italy | Chardonnay | Europe | 3309C |
| IV | SRR18048247 | Italy | Chardonnay | Europe | 3309C |
| IV | SRR18048248 | Italy | Chardonnay | Europe | 3309C |
| IV | SRR18048249 | Italy | Chardonnay | Europe | 3309C |
| IV | SRR18048250 | Italy | Sangiovese | Europe | 420A |
| IV | SRR18048251 | Italy | Chardonnay | Europe | 110R |

| | | | | | |
|---|---|---|---|---|---|
| IV | SRR18048252 | Italy | Chardonnay | Europe | 110R |
| IV | SRR18048253 | Italy | Chardonnay | Europe | 110R |
| IV | SRR18048254 | Italy | Chardonnay | Europe | 110R |
| IV | SRR18048255 | Italy | Chardonnay | Europe | 110R |
| IV | SRR18048256 | Italy | Chardonnay | Europe | 110R |
| IV | SRR18048257 | Italy | Chardonnay | Europe | SO4 |
| IV | SRR18048258 | Italy | Chardonnay | Europe | SO4 |
| IV | SRR18048259 | Italy | Chardonnay | Europe | SO4 |
| IV | SRR18048260 | Italy | Chardonnay | Europe | SO4 |
| IV | SRR18048261 | Italy | Sangiovese | Europe | 420A |
| IV | SRR18048262 | Italy | Chardonnay | Europe | SO4 |
| IV | SRR18048263 | Italy | Chardonnay | Europe | SO4 |
| IV | SRR18048264 | Italy | Merlot | Europe | 420A |
| IV | SRR18048265 | Italy | Merlot | Europe | 420A |
| IV | SRR18048266 | Italy | Merlot | Europe | 420A |
| IV | SRR18048267 | Italy | Merlot | Europe | 420A |
| IV | SRR18048268 | Italy | Merlot | Europe | 420A |
| IV | SRR18048269 | Italy | Chardonnay | Europe | 420A |
| IV | SRR18048270 | Italy | Merlot | Europe | 420A |
| IV | SRR18048271 | Italy | Chardonnay | Europe | 420A |
| IV | SRR18048272 | Italy | Chardonnay | Europe | 420A |
| IV | SRR18048273 | Italy | Chardonnay | Europe | 420A |
| IV | SRR18048274 | Italy | Chardonnay | Europe | SO4 |
| IV | SRR18048275 | Italy | Sangiovese | Europe | 420A |
| IV | SRR18048276 | Italy | Chardonnay | Europe | 420A |
| IV | SRR18048277 | Italy | Chardonnay | Europe | 420A |
| IV | SRR18048278 | Italy | Chardonnay | Europe | 420A |
| IV | SRR18048279 | Italy | Chardonnay | Europe | 420A |
| IV | SRR18048280 | Italy | Chardonnay | Europe | 420A |
| IV | SRR18048281 | Italy | Chardonnay | Europe | 420A |
| IV | SRR18048282 | Italy | Chardonnay | Europe | 420A |
| IV | SRR18048283 | Italy | Sangiovese | Europe | 110R |
| IV | SRR18048284 | Italy | Sangiovese | Europe | 110R |
| IV | SRR18048285 | Italy | Sangiovese | Europe | 110R |
| IV | SRR18048286 | Italy | Sangiovese | Europe | 110R |
| IV | SRR18048287 | Italy | Sangiovese | Europe | 110R |
| IV | SRR18048288 | Italy | Sangiovese | Europe | 110R |
| IV | SRR18048289 | Italy | Merlot | Europe | 420A |
| IV | SRR18048290 | Italy | Merlot | Europe | 420A |
| IV | SRR18048291 | Italy | Merlot | Europe | 420A |
| IV | SRR18048292 | Italy | Merlot | Europe | 420A |
| IV | SRR18048293 | Italy | Sangiovese | Europe | 420A |
| IV | SRR18048294 | Italy | Merlot | Europe | 420A |
| IV | SRR18048295 | Italy | Merlot | Europe | 420A |

| | | | | | |
|---|---|---|---|---|---|
| IV | SRR18048296 | Italy | Merlot | Europe | 420A |
| IV | SRR18048297 | Italy | Merlot | Europe | 420A |
| IV | SRR18048298 | Italy | Merlot | Europe | 420A |
| IV | SRR18048299 | Italy | Chardonnay | Europe | 420A |
| IV | SRR18048300 | Italy | Chardonnay | Europe | 420A |
| IV | SRR18048301 | Italy | Sangiovese | Europe | 420A |
| IV | SRR18048302 | Italy | Sangiovese | Europe | 420A |

| Description |
| --- |
| bulk soil |
| bulk soil |
| bulk soil |
| bulk soil |
| bulk soil |
| bulk soil |
| bulk soil |
| bulk soil |
| bulk soil |
| bulk soil |
| bulk soil |
| bulk soil |
| bulk soil |
| bulk soil |
| bulk soil |
| bulk soil |
| bulk soil |
| bulk soil |
| bulk soil |
| bulk soil |
| bulk soil |
| bulk soil |
| bulk soil |
| bulk soil |
| bulk soil |
| bulk soil |
| bulk soil |
| bulk soil |
| bulk soil |
| bulk soil |
| bulk soil |
| bulk soil |
| bulk soil |
| bulk soil |
| bulk soil |
| bulk soil |
| bulk soil |
| bulk soil |
| bulk soil |
| bulk soil |
| bulk soil |
| bulk soil |

bulk soil
bulk soil
bulk soil
bulk soil
bulk soil
bulk soil
bulk soil
bulk soil
bulk soil
bulk soil
bulk soil
bulk soil
bulk soil
bulk soil
bulk soil
bulk soil
bulk soil
bulk soil
bulk soil
bulk soil
bulk soil
bulk soil
bulk soil
bulk soil
bulk soil
bulk soil
bulk soil
bulk soil
bulk soil
bulk soil
bulk soil
bulk soil
bulk soil
bulk soil
bulk soil
bulk soil
bulk soil
bulk soil
bulk soil
bulk soil
bulk soil
bulk soil
bulk soil
bulk soil

bulk soil
bulk soil
bulk soil
bulk soil
bulk soil
bulk soil
bulk soil
bulk soil
bulk soil
bulk soil
bulk soil
bulk soil
bulk soil
bulk soil
bulk soil
bulk soil
bulk soil
bulk soil
bulk soil
bulk soil
bulk soil
bulk soil
bulk soil
bulk soil
bulk soil
bulk soil
bulk soil
bulk soil
bulk soil
bulk soil
bulk soil
bulk soil
bulk soil
bulk soil
bulk soil
bulk soil
bulk soil
bulk soil
bulk soil
bulk soil
bulk soil
bulk soil
bulk soil
bulk soil

bulk soil
bulk soil
bulk soil
bulk soil
bulk soil
bulk soil
bulk soil
bulk soil
bulk soil
bulk soil
bulk soil
bulk soil
bulk soil
bulk soil
bulk soil
bulk soil
bulk soil
bulk soil
bulk soil
bulk soil
bulk soil
bulk soil
bulk soil
bulk soil
bulk soil
bulk soil
bulk soil
bulk soil
bulk soil
bulk soil
bulk soil
bulk soil
bulk soil
bulk soil
bulk soil
bulk soil
bulk soil
bulk soil
bulk soil
bulk soil
bulk soil
bulk soil
bulk soil
bulk soil

bulk soil
bulk soil
bulk soil
bulk soil
bulk soil
bulk soil
bulk soil
bulk soil
bulk soil
bulk soil
bulk soil
bulk soil
bulk soil
bulk soil
bulk soil
bulk soil
bulk soil
bulk soil
bulk soil
bulk soil
bulk soil
bulk soil
bulk soil
bulk soil
bulk soil
bulk soil
bulk soil
bulk soil
bulk soil
bulk soil
bulk soil
bulk soil
bulk soil
bulk soil
bulk soil
bulk soil
bulk soil
bulk soil
bulk soil
bulk soil
bulk soil
bulk soil
bulk soil
bulk soil

bulk soil
bulk soil
bulk soil
bulk soil
bulk soil
bulk soil
bulk soil
bulk soil
bulk soil
bulk soil
bulk soil
bulk soil
bulk soil
bulk soil
bulk soil
bulk soil
bulk soil
bulk soil
bulk soil
bulk soil
bulk soil
bulk soil
bulk soil
bulk soil
bulk soil
bulk soil
bulk soil
bulk soil
bulk soil
bulk soil
bulk soil
bulk soil
bulk soil
bulk soil
bulk soil
bulk soil
bulk soil
bulk soil
bulk soil
bulk soil
bulk soil
bulk soil
bulk soil
bulk soil

bulk soil
bulk soil
bulk soil
bulk soil
bulk soil
bulk soil
bulk soil
bulk soil
bulk soil
bulk soil
bulk soil
bulk soil
bulk soil
bulk soil
bulk soil
bulk soil
bulk soil
bulk soil
bulk soil
bulk soil
bulk soil
bulk soil
bulk soil
bulk soil
bulk soil
bulk soil
bulk soil
bulk soil
bulk soil
bulk soil
bulk soil
bulk soil
bulk soil
bulk soil
bulk soil
bulk soil
bulk soil
bulk soil
bulk soil
bulk soil
bulk soil
bulk soil
bulk soil
bulk soil

bulk soil
bulk soil
bulk soil
bulk soil
bulk soil
bulk soil
bulk soil
bulk soil
bulk soil
bulk soil
bulk soil
bulk soil
rizosphere
rizosphere
bulk soil
bulk soil
bulk soil
rizosphere
bulk soil
rizosphere
grapes
bulk soil
rizosphere
bulk soil
bulk soil
rizosphere
bulk soil
bulk soil
rizosphere
bulk soil
bulk soil
bulk soil
bulk soil
bulk soil
bulk soil
bulk soil
rizosphere
bulk soil
bulk soil
rizosphere
rizosphere
rizosphere
bulk soil
bulk soil

bulk soil
bulk soil
bulk soil
bulk soil
bulk soil
bulk soil
rizosphere
bulk soil
leaves
rizosphere
rizosphere
leaves
bulk soil
rizosphere
bulk soil
rizosphere
bulk soil
bulk soil
bulk soil
bulk soil
rizosphere
bulk soil
rizosphere
rizosphere
bulk soil
rizosphere
rizosphere
bulk soil
bulk soil
bulk soil
rizosphere
bulk soil
rizosphere
bulk soil
rizosphere
bulk soil
bulk soil
bulk soil
bulk soil
bulk soil
bulk soil
rizosphere
rizosphere
bulk soil

bulk soil
rizosphere
bulk soil
bulk soil
rizosphere
bulk soil
bulk soil
rizosphere
rizosphere
rizosphere
bulk soil
rizosphere
bulk soil
bulk soil
bulk soil
bulk soil
grapes
rizosphere
bulk soil
bulk soil
bulk soil
rizosphere
grapes
rizosphere
bulk soil
bulk soil
rizosphere
bulk soil
bulk soil
bulk soil
bulk soil
bulk soil
leaves
rizosphere
rizosphere
bulk soil
rizosphere
bulk soil
bulk soil
bulk soil
rizosphere
rizosphere
bulk soil
bulk soil

bulk soil
bulk soil
bulk soil
bulk soil
bulk soil
bulk soil
bulk soil
rizosphere
rizosphere
bulk soil
bulk soil
rizosphere
rizosphere
rizosphere
bulk soil
bulk soil
rizosphere
rizosphere
bulk soil
bulk soil
bulk soil
bulk soil
bulk soil
bulk soil
bulk soil
rizosphere
bulk soil
rizosphere
bulk soil
bulk soil
rizosphere
rizosphere
bulk soil
rizosphere
leaves
bulk soil
bulk soil
rizosphere
rizosphere
bulk soil
bulk soil
rizosphere
rizosphere
rizosphere

rizosphere
leaves
leaves
bulk soil
bulk soil
bulk soil
grapes
bulk soil
rizosphere
bulk soil
rizosphere
rizosphere
rizosphere
bulk soil
bulk soil
bulk soil
bulk soil
bulk soil
rizosphere
bulk soil
bulk soil
bulk soil
bulk soil
bulk soil
rizosphere
bulk soil
bulk soil
bulk soil
grapes
rizosphere
rizosphere
bulk soil
rizosphere
rizosphere
rizosphere
rizosphere
rizosphere
bulk soil
bulk soil
rizosphere
rizosphere
bulk soil
rizosphere
bulk soil

bulk soil
bulk soil
bulk soil
rizosphere
bulk soil
rizosphere
bulk soil
bulk soil
bulk soil
bulk soil
bulk soil
bulk soil
bulk soil
rizosphere
bulk soil
rizosphere
bulk soil
bulk soil
bulk soil
rizosphere
rizosphere
grapes
rizosphere
rizosphere
bulk soil
rizosphere
bulk soil
rizosphere
bulk soil
bulk soil
bulk soil
bulk soil
rizosphere
bulk soil
bulk soil
bulk soil
rizosphere
bulk soil
bulk soil
bulk soil
bulk soil
rhizosphere
rhizosphere
bulk soil

bulk soil
rhizosphere
bulk soil
rhizosphere
bulk soil
bulk soil
rhizosphere
bulk soil
rhizosphere
bulk soil
bulk soil
rhizosphere
bulk soil
rhizosphere
bulk soil
bulk soil
bulk soil
rhizosphere
bulk soil
rhizosphere
bulk soil
rhizosphere
bulk soil
rhizosphere
bulk soil
bulk soil
rhizosphere
bulk soil
rhizosphere
bulk soil
bulk soil
rhizosphere
bulk soil
rhizosphere
bulk soil
rhizosphere
bulk soil
rhizosphere
bulk soil
rhizosphere
bulk soil
rhizosphere
bulk soil
rhizosphere

rhizosphere
bulk soil
bulk soil
rhizosphere
bulk soil
rhizosphere
bulk soil
rhizosphere
bulk soil
rhizosphere
bulk soil
bulk soil
rhizosphere
bulk soil
rhizosphere
bulk soil
bulk soil
rhizosphere
bulk soil
rhizosphere
bulk soil
bulk soil
rhizosphere
bulk soil
rhizosphere
bulk soil
rhizosphere
bulk soil
bulk soil
rhizosphere
bulk soil
rhizosphere
bulk soil
rhizosphere
bulk soil
rhizosphere
bulk soil
bulk soil
rhizosphere
bulk soil
rhizosphere
bulk soil
bulk soil
bulk soil

rhizosphere
bulk soil
bulk soil
bulk soil
rhizosphere
rhizosphere
bulk soil
rhizosphere
bulk soil
rhizosphere
rhizosphere
bulk soil
rhizosphere
bulk soil
bulk soil
rhizosphere
bulk soil
rhizosphere
bulk soil
rhizosphere
bulk soil
rhizosphere
bulk soil
rhizosphere
bulk soil
rhizosphere
bulk soil
rhizosphere
bulk soil
rhizosphere
bulk soil
rhizosphere
rhizosphere
bulk soil
rhizosphere
bulk soil
bulk soil
bulk soil
rhizosphere
bulk soil
rhizosphere
bulk soil
rhizosphere
bulk soil

rhizosphere
bulk soil
rhizosphere
bulk soil
rhizosphere
rhizosphere
bulk soil
bulk soil
rhizosphere
bulk soil
rhizosphere
bulk soil
rhizosphere
bulk soil
rhizosphere
bulk soil
rhizosphere
bulk soil
rhizosphere
bulk soil
rhizosphere
bulk soil
bulk soil
rhizosphere
bulk soil
rhizosphere
bulk soil
bulk soil
rhizosphere
bulk soil
rhizosphere
bulk soil
bulk soil
rhizosphere
bulk soil
rhizosphere
bulk soil
bulk soil
rhizosphere
bulk soil
rhizosphere
bulk soil
bulk soil
bulk soil

bulk soil
bulk soil
bulk soil
bulk soil
bulk soil
bulk soil
bulk soil
bulk soil
bulk soil
bulk soil
bulk soil
bulk soil
bulk soil
bulk soil
bulk soil
bulk soil
bulk soil
bulk soil
bulk soil
bulk soil
bulk soil
bulk soil
bulk soil
bulk soil
bulk soil
bulk soil
bulk soil
bulk soil
bulk soil
bulk soil
bulk soil
bulk soil
bulk soil
bulk soil
bulk soil
bulk soil
bulk soil
bulk soil
bulk soil
bulk soil
bulk soil
bulk soil
bulk soil
bulk soil

bulk soil
bulk soil
bulk soil
bulk soil
bulk soil
bulk soil
bulk soil
bulk soil
bulk soil
bulk soil
bulk soil
bulk soil
bulk soil
bulk soil
bulk soil
bulk soil
bulk soil
bulk soil
bulk soil
bulk soil
bulk soil
bulk soil
bulk soil
bulk soil
bulk soil
bulk soil
rhizosphere
rhizosphere
rhizosphere
rhizosphere
rhizosphere
rhizosphere
rhizosphere
rhizosphere
rhizosphere
rhizosphere
rhizosphere
rhizosphere
rhizosphere
rhizosphere
rhizosphere
rhizosphere
rhizosphere
rhizosphere

rhizosphere
rhizosphere
rhizosphere
rhizosphere
rhizosphere
rhizosphere
rhizosphere
rhizosphere
rhizosphere
rhizosphere
rhizosphere
rhizosphere
rhizosphere
rhizosphere
rhizosphere
rhizosphere
rhizosphere
rhizosphere
rhizosphere
rhizosphere
rhizosphere
rhizosphere
rhizosphere
rhizosphere
rhizosphere
rhizosphere
rhizosphere
rhizosphere
rhizosphere
rhizosphere
rhizosphere
rhizosphere
rhizosphere
rhizosphere
rhizosphere
rhizosphere
rhizosphere
rhizosphere
rhizosphere
rhizosphere
rhizosphere
rhizosphere
rhizosphere
rhizosphere

rhizosphere
rhizosphere
rhizosphere
rhizosphere
rhizosphere
rhizosphere
rhizosphere
