## Supplementary Table 3 for "Grapevine Rootstock and Scion Genotypes’ Symbiosis with Soil Microbiome: A Machine Learning Revelation for Climate-Resilient Viticulture"

| Dataset_ID | PROJ_alias | Sample_ID | common_name | Organism | orig_sample_name |
|---|---|---|---|---|---|
| III | merlot_project | ERR557127 | soil metagenome | soil metagenome | SH008.C6.HA.2.735 |
| III | merlot_project | ERR557128 | soil metagenome | soil metagenome | HE003.C181.HA.4.769 |
| III | merlot_project | ERR557132 | soil metagenome | soil metagenome | SH005.C3.RH.1.655 |
| III | merlot_project | ERR557137 | soil metagenome | soil metagenome | GM.Grapes.181.R1 |
| III | merlot_project | ERR557142 | soil metagenome | soil metagenome | SH004.C1.RH.3.614 |
| III | merlot_project | ERR557146 | soil metagenome | soil metagenome | SH007.C6.RH.5.725 |
| III | merlot_project | ERR557147 | soil metagenome | soil metagenome | HE003.C181.HA.2.759 |
| III | merlot_project | ERR557151 | soil metagenome | soil metagenome | MU002.C3.HA.5.699 |
| III | merlot_project | ERR557152 | soil metagenome | soil metagenome | DPOO1.C1.HA.3.32 |
| III | merlot_project | ERR557156 | soil metagenome | soil metagenome | HE003.C181.HA.4.770 |
| III | merlot_project | ERR557159 | soil metagenome | soil metagenome | MU002.C3.HA.2.685 |
| III | merlot_project | ERR557160 | soil metagenome | soil metagenome | RU006.C181.RH.2.785 |
| III | merlot_project | ERR557161 | soil metagenome | soil metagenome | MU002.C3.HA.2.684 |
| III | merlot_project | ERR557163 | soil metagenome | soil metagenome | SH004.C1.RH.5.624 |
| III | merlot_project | ERR557170 | soil metagenome | soil metagenome | SH008.C6.HA.2.734 |
| III | merlot_project | ERR557181 | soil metagenome | soil metagenome | HE003.C181.HA.2.760 |
| III | merlot_project | ERR557182 | soil metagenome | soil metagenome | SH004.C1.RH.3.215.r1 |
| III | merlot_project | ERR557183 | soil metagenome | soil metagenome | DPOO1.C1.HA.2.634 |
| III | merlot_project | ERR557184 | soil metagenome | soil metagenome | SH008.C6.HA.4.745 |
| III | merlot_project | ERR557186 | soil metagenome | soil metagenome | DPOO1.C1.HA.3.640 |
| III | merlot_project | ERR557188 | soil metagenome | soil metagenome | DPOO1.C1.HA.5.649 |
| III | merlot_project | ERR557192 | soil metagenome | soil metagenome | SH005.C3.RH.5.675 |
| III | merlot_project | ERR557193 | soil metagenome | soil metagenome | GM.Leaves.181.R3 |
| III | merlot_project | ERR557196 | soil metagenome | soil metagenome | GM.Leaves.181.R4 |
| III | merlot_project | ERR557203 | soil metagenome | soil metagenome | HE003.C181.HA.5.774 |
| III | merlot_project | ERR557208 | soil metagenome | soil metagenome | HE003.C181.HA.3.765 |
| III | merlot_project | ERR557221 | soil metagenome | soil metagenome | MU002.C3.HA.4.695 |
| III | merlot_project | ERR557225 | soil metagenome | soil metagenome | HE003.C181.HA.1.754 |
| III | merlot_project | ERR557226 | soil metagenome | soil metagenome | SH008.C6.HA.3.740 |
| III | merlot_project | ERR557230 | soil metagenome | soil metagenome | HE003.C181.HA.3.764 |
| III | merlot_project | ERR557233 | soil metagenome | soil metagenome | DPOO1.C1.HA.5.650 |
| III | merlot_project | ERR557237 | soil metagenome | soil metagenome | RU006.C181.RH.1.779 |
| III | merlot_project | ERR557242 | soil metagenome | soil metagenome | DPOO1.C1.HA.1.629 |
| III | merlot_project | ERR557249 | soil metagenome | soil metagenome | MU002.C3.HA.4.295.r1 |
| III | merlot_project | ERR557250 | soil metagenome | soil metagenome | RU006.C181.RH.5.800 |
| III | merlot_project | ERR557260 | soil metagenome | soil metagenome | SH007.C6.RH.2.709 |
| III | merlot_project | ERR557278 | soil metagenome | soil metagenome | RU006.C181.RH.3.789 |
| III | merlot_project | ERR557281 | soil metagenome | soil metagenome | GM.Grapes.181.R4 |
| III | merlot_project | ERR557287 | soil metagenome | soil metagenome | SH004.C1.RH.2.609 |
| III | merlot_project | ERR557291 | soil metagenome | soil metagenome | GM.Grapes.181.R5 |
| III | merlot_project | ERR557298 | soil metagenome | soil metagenome | SH005.C3.RH.5.674 |
| III | merlot_project | ERR557301 | soil metagenome | soil metagenome | MU002.C3.HA.3.690 |
| III | merlot_project | ERR557302 | soil metagenome | soil metagenome | SH004.C1.RH.1.605 |
| III | merlot_project | ERR557303 | soil metagenome | soil metagenome | SH007.C6.RH.3.715 |
| III | merlot_project | ERR557304 | soil metagenome | soil metagenome | SH008.C6.HA.5.750 |
| III | merlot_project | ERR557306 | soil metagenome | soil metagenome | RU006.C181.RH.2.784 |
| III | merlot_project | ERR557309 | soil metagenome | soil metagenome | GM.Leaves.181.R5 |
| III | merlot_project | ERR557316 | soil metagenome | soil metagenome | MU002.C3.HA.4.694 |
| III | merlot_project | ERR557317 | soil metagenome | soil metagenome | SH005.C3.RH.4.270.r1 |

| | | | | | |
|---|---|---|---|---|---|
| III | merlot_project | ERR557324 | soil metagenome | soil metagenome | RU006.C181.RH.4.795 |
| III | merlot_project | ERR557328 | soil metagenome | soil metagenome | SH005.C3.RH.2.659 |
| III | merlot_project | ERR557329 | soil metagenome | soil metagenome | RU006.C181.RH.3.790 |
| III | merlot_project | ERR557331 | soil metagenome | soil metagenome | SH008.C6.HA.3.739 |
| III | merlot_project | ERR557336 | soil metagenome | soil metagenome | RU006.C181.RH.1.780 |
| III | merlot_project | ERR557343 | soil metagenome | soil metagenome | SH005.C3.RH.3.665 |
| III | merlot_project | ERR557349 | soil metagenome | soil metagenome | HE003.C181.HA.5.775 |
| III | merlot_project | ERR557354 | soil metagenome | soil metagenome | SH008.C6.HA.1.730 |
| III | merlot_project | ERR557355 | soil metagenome | soil metagenome | SH008.C6.HA.1.729 |
| III | merlot_project | ERR557371 | soil metagenome | soil metagenome | RU006.C181.RH.5.799 |
| III | merlot_project | ERR557376 | soil metagenome | soil metagenome | SH004.C1.RH.2.610 |
| III | merlot_project | ERR557379 | soil metagenome | soil metagenome | GM.Leaves.181.R1 |
| III | merlot_project | ERR557381 | soil metagenome | soil metagenome | SH005.C3.RH.1.654 |
| III | merlot_project | ERR557388 | soil metagenome | soil metagenome | SH007.C6.RH.4.720 |
| III | merlot_project | ERR557390 | soil metagenome | soil metagenome | SH008.C6.HA.5.749 |
| III | merlot_project | ERR557399 | soil metagenome | soil metagenome | SH008.C6.HA.1.SH008.leav |
| III | merlot_project | ERR557400 | soil metagenome | soil metagenome | GM.Leaves.181.R2 |
| III | merlot_project | ERR557405 | soil metagenome | soil metagenome | GM.Grapes.181.R2 |
| III | merlot_project | ERR557409 | soil metagenome | soil metagenome | MU002.C3.HA.1.679 |
| III | merlot_project | ERR557414 | soil metagenome | soil metagenome | SH004.C1.RH.4.619 |
| III | merlot_project | ERR557424 | soil metagenome | soil metagenome | SH008.C6.HA.4.744 |
| III | merlot_project | ERR557428 | soil metagenome | soil metagenome | SH004.C1.RH.4.620 |
| III | merlot_project | ERR557429 | soil metagenome | soil metagenome | SH007.C6.RH.5.724 |
| III | merlot_project | ERR557430 | soil metagenome | soil metagenome | HE003.C181.HA.1.755 |
| III | merlot_project | ERR557431 | soil metagenome | soil metagenome | DPOO1.C1.HA.3.639 |
| III | merlot_project | ERR557437 | soil metagenome | soil metagenome | RU006.C181.RH.4.794 |
| III | merlot_project | ERR557439 | soil metagenome | soil metagenome | SH008.C6.HA.1.SH008.gp |
| III | merlot_project | ERR557457 | soil metagenome | soil metagenome | SH007.C6.RH.3.714 |
| III | merlot_project | ERR557461 | soil metagenome | soil metagenome | SH004.C1.RH.1.604 |
| III | merlot_project | ERR557466 | soil metagenome | soil metagenome | MU002.C3.HA.5.700 |
| III | merlot_project | ERR557467 | soil metagenome | soil metagenome | MU002.C3.HA.3.689 |
| III | merlot_project | ERR557474 | soil metagenome | soil metagenome | SH005.C3.RH.3.664 |
| III | merlot_project | ERR557475 | soil metagenome | soil metagenome | SH004.C1.RH.3.615 |
| III | merlot_project | ERR557476 | soil metagenome | soil metagenome | DPOO1.C1.HA.2.635 |
| III | merlot_project | ERR557477 | soil metagenome | soil metagenome | DPOO1.C1.HA.1.630 |
| III | merlot_project | ERR557482 | soil metagenome | soil metagenome | SH004.C1.RH.2.8.r1 |
| III | merlot_project | ERR557485 | soil metagenome | soil metagenome | SH007.C6.RH.2.710 |
| III | merlot_project | ERR557487 | soil metagenome | soil metagenome | SH007.C6.RH.4.719 |
| III | merlot_project | ERR557489 | soil metagenome | soil metagenome | SH005.C3.RH.4.670 |
| III | merlot_project | ERR557494 | soil metagenome | soil metagenome | GM.Grapes.181.R3 |
| III | merlot_project | ERR557502 | soil metagenome | soil metagenome | MU002.C3.HA.4.76.r1 |
| III | merlot_project | ERR557507 | soil metagenome | soil metagenome | SH005.C3.RH.4.669 |
| III | merlot_project | ERR557511 | soil metagenome | soil metagenome | SH004.C1.RH.5.625 |
| III | merlot_project | ERR557512 | soil metagenome | soil metagenome | SH007.C6.RH.1.704 |
| III | merlot_project | ERR557518 | soil metagenome | soil metagenome | DPOO1.C1.HA.4.644 |
| III | merlot_project | ERR557522 | soil metagenome | soil metagenome | DPOO1.C1.HA.4.645 |
| III | merlot_project | ERR557523 | soil metagenome | soil metagenome | MU002.C3.HA.1.680 |

| Library.Name |
| --- |
| SH008.C6.HA.2.735.gp.9.12:374 |
| HE003.C181.HA.4.769.leav.9.12:110 |
| SH005.C3.RH.1.655.gp.9.12:272 |
| GM.181.R1.gp.10.12:49 |
| SH004.C1.RH.3.614.leav.9.12:244 |
| SH007.C6.RH.5.725.gp.9.12:352 |
| HE003.C181.HA.2.759.leav.9.12:89 |
| MU002.C3.HA.5.699.leav.9.12:167 |
| DPOO1.C1.HA.3.32.r1.leav.6.11:20 |
| HE003.C181.HA.4.770.gp.9.12:111 |
| MU002.C3.HA.2.685.gp.9.12:139 |
| RU006.C181.RH.2.785.gp.9.12:188 |
| MU002.C3.HA.2.684.leav.9.12:138 |
| SH004.C1.RH.5.624.leav.9.12:263 |
| SH008.C6.HA.2.734.leav.9.12:373 |
| HE003.C181.HA.2.760.gp.9.12:90 |
| SH004.C1.RH.3.215.r1.gp.9.11:237 |
| DPOO1.C1.HA.2.634.leav.9.12:16 |
| SH008.C6.HA.4.745.gp.9.12:392 |
| DPOO1.C1.HA.3.640.gp.9.12:29 |
| DPOO1.C1.HA.5.649.leav.9.12:46 |
| SH005.C3.RH.5.675.gp.9.12:307 |
| GM.181.R3.leav.10.12:60 |
| GM.181.R4.leav.10.12:65 |
| HE003.C181.HA.5.774.leav.9.12:120 |
| HE003.C181.HA.3.765.gp.9.12:101 |
| MU002.C3.HA.4.695.gp.9.12:157 |
| HE003.C181.HA.1.754.leav.9.12:80 |
| SH008.C6.HA.3.740.gp.9.12:383 |
| HE003.C181.HA.3.764.leav.9.12:100 |
| DPOO1.C1.HA.5.650.gp.9.12:47 |
| RU006.C181.RH.1.779.leav.9.12:177 |
| DPOO1.C1.HA.1.629.leav.9.12:8 |
| MU002.C3.HA.4.295.r1.gp.9.11:150 |
| RU006.C181.RH.5.800.gp.9.12:213 |
| SH007.C6.RH.2.709.leav.9.12:323 |
| RU006.C181.RH.3.789.leav.9.12:196 |
| GM.181.R4.gp.10.12:64 |
| SH004.C1.RH.2.609.leav.9.12:231 |
| GM.181.R5.gp.10.12:69 |
| SH005.C3.RH.5.674.leav.9.12:306 |
| MU002.C3.HA.3.690.gp.9.12:147 |
| SH004.C1.RH.1.605.gp.9.12:223 |
| SH007.C6.RH.3.715.gp.9.12:333 |
| SH008.C6.HA.5.750.gp.9.12:400 |
| RU006.C181.RH.2.784.leav.9.12:187 |
| GM.181.R5.leav.10.12:70 |
| MU002.C3.HA.4.694.leav.9.12:156 |
| SH005.C3.RH.4.270.r1.gp.9.11:289 |

RU006.C181.RH.4.795.gp.9.12:205
SH005.C3.RH.2.659.leav.9.12:279
RU006.C181.RH.3.790.gp.9.12:197
SH008.C6.HA.3.739.leav.9.12:382
RU006.C181.RH.1.780.gp.9.12:178
SH005.C3.RH.3.665.gp.9.12:287
HE003.C181.HA.5.775.gp.9.12:121
SH008.C6.HA.1.730.gp.9.12:361
SH008.C6.HA.1.729.leav.9.12:360
RU006.C181.RH.5.799.leav.9.12:212
SH004.C1.RH.2.610.gp.9.12:232
GM.181.R1.leav.10.12:50
SH005.C3.RH.1.654.leav.9.12:271
SH007.C6.RH.4.720.gp.9.12:342
SH008.C6.HA.5.749.leav.9.12:399
SH008.C6.HA.1.SH008.leav.leav.9.12:363
GM.181.R2.leav.10.12:55
GM.181.R2.gp.10.12:54
MU002.C3.HA.1.679.leav.9.12:129
SH004.C1.RH.4.619.leav.9.12:254
SH008.C6.HA.4.744.leav.9.12:391
SH004.C1.RH.4.620.gp.9.12:255
SH007.C6.RH.5.724.leav.9.12:351
HE003.C181.HA.1.755.gp.9.12:81
DPOO1.C1.HA.3.639.leav.9.12:28
RU006.C181.RH.4.794.leav.9.12:204
SH008.C6.HA.1.SH008.gp.gp.9.12:362
SH007.C6.RH.3.714.leav.9.12:332
SH004.C1.RH.1.604.leav.9.12:222
MU002.C3.HA.5.700.gp.9.12:168
MU002.C3.HA.3.689.leav.9.12:146
SH005.C3.RH.3.664.leav.9.12:286
SH004.C1.RH.3.615.gp.9.12:245
DPOO1.C1.HA.2.635.gp.9.12:17
DPOO1.C1.HA.1.630.gp.9.12:9
SH004.C1.RH.2.8.r1.leav.6.11:234
SH007.C6.RH.2.710.gp.9.12:324
SH007.C6.RH.4.719.leav.9.12:341
SH005.C3.RH.4.670.gp.9.12:297
GM.181.R3.gp.10.12:59
MU002.C3.HA.4.76.r1.leav.6.11:159
SH005.C3.RH.4.669.leav.9.12:296
SH004.C1.RH.5.625.gp.9.12:264
SH007.C6.RH.1.704.leav.9.12:315
DPOO1.C1.HA.4.644.leav.9.12:38
DPOO1.C1.HA.4.645.gp.9.12:39
MU002.C3.HA.1.680.gp.9.12:130
