## Supplementary Table 4 for "Grapevine Rootstock and Scion Genotypes’ Symbiosis with Soil Microbiome: A Machine Learning Revelation for Climate-Resilient Viticulture"

**Supplementary Table 4.** List of hyper-parameters used for the all ML algorithms, except neural networks, along with the respective set of values used for grid search and the values from the corresponding best-performing models. The cross validation (CV) values column depicts a list of the values used for the corresponding parameter. A list is depicted as a square bracket where elements in a list are comma separated.

| ML Method | parameters | CV values | Best Model Values | | |
|---|---|---|---|---|---|
| | | | Continent | Country | Cultivar |
| RF | n_estimators | [1000, 1500] | 1500 | 1000 | 1000 |
| | max_feature s | [10000, 35000, 45000, 60000], | 45000 | 45000 | 45000 |
| | class_weight | [balanced, balanced_subsample] | balanced subsample | balanced subsample | balanced subsample |
| | max_depth | [1000, 2000, 5000, 10000, 50000] | 1000 | 1000 | 1000 |
| ADA | n_estimators | [5000, 10000] | 10000 | 10000 | 10000 |
| | estimator | Decision Tree (max_depth = 1000, max_features = 45000, class_weight= 'balanced') | | | |
| GBM | n_estimators | [1000,1500] | 1000 | 1000 | 1000 |
| | max_feature s | [35000, 45000, 70000] | 35000 | 35000 | 35000 |
| | max_depth | [1000, 2000, 5000] | 1000 | 1000 | 1000 |
| GNB | priors | [None, 1/ ( no. of classes )] | 0.2 | 0.083 | 0.0526 |
| BNB | fit_prior | [false, true] | both | both | both |
| | binarize | None | | | |
| KNN | n_neighbors | [5, 10, 15, 20] | 10 | 5 | 5 |
| | leaf_size | [30, 100, 200, 1000, 10000, 30000] | 30 | 30 | 30 |
| SVML | class_weight | balanced | balanced | balanced | balanced |
| | gamma | [scale, auto] | scale | scale | scale |
| | C | [0.01, 0.1, 0.5, 1.0] | 0.01 | 0.01 | 0.01 |
| SVMR | class_weight | balanced | balanced | balanced | balanced |
| | gamma | [scale, auto] | scale | scale | scale |
| | C | [0.01, 0.1, 0.5, 1.0] | 1 | 1 | 1 |
