## Supplementary Table 5 for "Grapevine Rootstock and Scion Genotypes’ Symbiosis with Soil Microbiome: A Machine Learning Revelation for Climate-Resilient Viticulture"

**Supplementary Table 5. Neural Network hyper-parameters for Appr[illegible]**
approach A. These are the final hyperparameters used to obtain the b[illegible]

| Hyper-Parameter | Continent |
|---|---|
| | **Architectu[illegible]** |
| Hidden layers | 6 |
| Neurons per layer | 800 (all layers) |
| Activation function (hidden) | ReLu |
| Activation function (output) | softmax |
| Loss function | Sparse Categorical Cross Entropy |
| Optimizer | Adam |
| learning rate | 0.001 |
| Weight initializer (per layer) | HeNormal |
| Kernal regularizer | L2 (lr = 0.001) |
| | **Trainin[illegible]** |
| Epochs | 500 |
| Batch Size | 10 |
| Class weight | balanced |

**oach A.** List of hyper-parameters used to build and train the NN for
est model.

| | Country | Cultivar |
|---|---|---|
| **ıre Parameters** | | |
| | 10 | 10 |
| | L1,L4,L7: 1000 | L1,L4,L7: 1000 |
| | L2,L4,L9:1200 | L2,L4,L9:1200 |
| | L3,L6: 1500 | L3,L6: 1500 |
| | L8,L10: 800 | L8,L10: 800 |
| | ReLu | ReLu |
| | softmax | softmax |
| | Sparse Categorical Cross Entropy | Sparse Categorical Cross Entropy |
| | SGD | SGD |
| | 0.001 | 0.001 |
| | HeNormal | HeNormal |
| | L2 (lr = 0.001) | L2 (lr = 0.001) |
| **g Parameters** | | |
| | 1500 | 3000 |
| | 64 | 32 |
| | balanced | balanced |
