## Supplementary Table 6 for "Grapevine Rootstock and Scion Genotypes’ Symbiosis with Soil Microbiome: A Machine Learning Revelation for Climate-Resilient Viticulture"

**Supplementary Table 6. Neural Network hyper-parameters for Approach B.** List of hyper-parameters used for grid search and select the optimal NN architecture. A list is depicted as a square bracket where elements in a list are comma-separated.

| **Hyper-Parameter** | **Value** |
|---|---|
| **Architecture Parameters** | |
| Hidden layers | [ 1, 2, 3 ] |
| Neurons per layer | [8,16,32,64,128,256,512,1024] |
| Activation function (hidden) | ReLu |
| Activation function (output) | softmax |
| Loss function | Focal loss |
| Optimizer | Adam |
| learning rate | 0.001 |
| Kernal regularizer | L2 (lr = 0.001) |
| **Training Parameters** | |
| Epochs | Early Stopping callback |
| Batch Size | 8 |
