## Supplementary Table 7 for "Grapevine Rootstock and Scion Genotypes’ Symbiosis with Soil Microbiome: A Machine Learning Revelation for Climate-Resilient Viticulture"

**Supplementary Table 7. Performance scores for continent classification using approach A.** Prec
(macro and weighted) of the nine machine learning (ML) algorithms, Random Forests (RF), Adap
Support Vector Machines with linear (SVML) and radial (SVMR) kernels, Gaussian (GNB) and Berı
Neighbor (KNN), and Neural Networks (NN) on classification of 5 continents using soil microbiom
an excluded 25% out of the test data.

| | RF | ADA | GBM | GNB | BNB | KNN |
|---|---|---|---|---|---|---|
| **Accuracy** | 0.95 | 0.9 | 0.97 | 0.96 | 0.86 | 0.65 |
| **Macro Avg** | 0.58 | 0.66 | 0.59 | 0.58 | 0.5 | 0.26 |
| **Weighted Avg** | 0.95 | 0.9 | 0.96 | 0.96 | 0.86 | 0.58 |

diction accuracies and F1-averages
tive Boost (ADA), Gradient Boost (GBM),
noulli Naïve Bayes (BNB), k-Nearest
ne data. The performances obtained on

| SVML | SVMR | NN |
|---|---|---|
| 0.95 | 0.91 | 0.97 |
| 0.58 | 0.55 | 0.79 |
| 0.94 | 0.9 | 0.97 |
