## Supplementary Table 8 for "Grapevine Rootstock and Scion Genotypes’ Symbiosis with Soil Microbiome: A Machine Learning Revelation for Climate-Resilient Viticulture"

**Supplementary Table 8. Performance scores for country classification using approach A.** Predic
and weighted) of the nine machine learning (ML) algorithms, Random Forests (RF), Adaptive Boo
Support Vector Machines with linear (SVML) and radial (SVMR) kernels, Gaussian (GNB) and Berı
Neighbor (KNN), and Neural Networks (NN) on classification of 12 countries using soil microbiom
an excluded 25% of the test data

| | RF | ADA | GBM | GNB | BNB | KNN |
|---|---|---|---|---|---|---|
| **Accuracy** | 0.94 | 0.9 | 0.95 | 0.92 | 0.8 | 0.55 |
| **Macro. Avg.** | 0.53 | 0.69 | 0.6 | 0.47 | 0.33 | 0.15 |
| **Weighted Av** | 0.93 | 0.9 | 0.94 | 0.9 | 0.78 | 0.42 |

:tion accuracies and F1-averages (macro
ost (ADA), Gradient Boost (GBM),
noulli Naïve Bayes (BNB), k-Nearest
ne data. The performances obtained on

| SVML | SVMR | NN |
|---|---|---|
| 0.96 | 0.87 | 0.97 |
| 0.66 | 0.46 | 0.74 |
| 0.95 | 0.86 | 0.96 |
