## Supplementary Table 9 for "Grapevine Rootstock and Scion Genotypes’ Symbiosis with Soil Microbiome: A Machine Learning Revelation for Climate-Resilient Viticulture"

**Supplementary Table 9. Performance scores for cultivar classification using approach A.** Predictio and weighted) of the nine machine learning (ML) algorithms, Random Forests (RF), Adaptive Boost Vector Machines with linear (SVML) and radial (SVMR) kernels, Gaussian (GNB) and Bernoulli Naïve and Neural Networks (NN) on classification of 19 cultivars using soil microbiome data. The performa data.

| | RF | ADA | GBM | GNB | BNB | KNN |
|---|---|---|---|---|---|---|
| **Accuracy** | 0.82 | 0.72 | 0.83 | 0.77 | 0.7 | 0.49 |
| **Macro Avg** | 0.52 | 0.43 | 0.47 | 0.32 | 0.18 | 0.21 |
| **Weighted Avg** | 0.8 | 0.72 | 0.81 | 0.74 | 0.66 | 0.47 |

n accuracies and F1-averages (macro
(ADA), Gradient Boost (GBM), Support
Bayes (BNB), k-Nearest Neighbor (KNN),
ances obtained on 25% withheld test

| SVML | SVMR | NN |
|---|---|---|
| 0.83 | 0.65 | 0.85 |
| 0.61 | 0.32 | 0.72 |
| 0.82 | 0.66 | 0.85 |
