## Supplementary Table 10 for "Grapevine Rootstock and Scion Genotypes’ Symbiosis with Soil Microbiome: A Machine Learning Revelation for Climate-Resilient Viticulture"

**Supplementary Table 10.** Performance scores of NN model for scion/rootstock combination classification. Prediction accuracy, F1, Precision, and Recall averages for Neural Network (NN) model for classification of 8 scion/rootstock combinations. Support is the total number of test samples.

| | Precision | Recall | F1-Score | Support |
|---|---|---|---|---|
| **Accuracy** | - | - | 0.97 | 118 |
| **Macro average** | 0.97 | 0.9 | 0.91 | 118 |
| **Weighted average** | 0.98 | 0.97 | 0.97 | 118 |
