## Supplementary Table 11 for "Grapevine Rootstock and Scion Genotypes’ Symbiosis with Soil Microbiome: A Machine Learning Revelation for Climate-Resilient Viticulture"

**Supplementary Table 11. Most informative ASVs for continent classification using RF.** List o from Random Forest (RF) model. The taxonomy was then predicted with classifiers trained on Gre shows the predictions from both. The taxonomic level of the predictions are indicated with a letter are used for ‘kingdom’, ‘phylum’, ‘class’, ‘order’, ‘family’, ‘genus’, and ‘species’ respectively. Th the species level. The confidence scores from the classifier are depicted in the adjacent columns.

| ASV ID | Taxon GG | Confidence GG | Taxon SILVA |
|---|---|---|---|
| 136760 | d__Bacteria; p__Nitrospirota_A_437815; c__Nitrospiria; o__Nitrospirales; f__Nitrospiraceae; g__*Nitrospira_* C | 1.000 | d__Bacteria; p__Nitrospirota; c__Nitrospiria; o__Nitrospirales; f__Nitrospiraceae; g__*Nitrospira* ; s__*Nitrospira_japonica* |
| 39ad699282d51b96d4f507f7ff11d8c6 | d__Bacteria; p__Proteobacteria; c__Alphaproteobacteria; o__Rhizobiales_A_504705; f__Xanthobacteraceae_503485;g__VAZQ01; s__VAZQ01 sp005883115 | 0.918 | d__Bacteria; p__Proteobacteria; c__Alphaproteobacteria; o__Rhizobiales; f__Xanthobacteraceae; g__uncultured |
| 575740 | d__Bacteria; p__Acidobacteriota; c__Vicinamibacteria; o__Vicinamibacterales; f__UBA2999; g__WHSN01; s__ | 0.860 | d__Bacteria; p__Acidobacteriota; c__Vicinamibacteria; o__Vicinamibacterales; f__uncultured; g__uncultured |
| 0f8ac0d81dd6449a43b34fcb184efd8f | d__Archaea; p__Thermoproteota; c__Nitrososphaeria_A; o__Nitrososphaerales; f__Nitrososphaeraceae; g__*Nitrosocosmicus* ; s__*Nitrosocosmicus hydrocola* | 0.756 | d__Archaea; p__Crenarchaeota; c__Nitrososphaeria; o__Nitrososphaerales; f__Nitrososphaeraceae |

| | | | |
|---|---|---|---|
| 210475 | d__Bacteria; p__Proteobacteria; c__Gammaproteobacteria; o__Burkholderiales_597439; f__Usitatibacteraceae; g__; s__ | 0.910 | d__Bacteria; p__Proteobacteria; c__Gammaproteobacteria; o__Burkholderiales; f__Nitrosomonadaceae; g__IS-44 |
| 818123 | d__Bacteria; p__Acidobacteriota; c__Blastocatellia; o__UBA7656; f__UBA7656; g__QHVH01; s__QHVH01 sp003222245 | 0.982 | d__Bacteria; p__Acidobacteriota; c__Blastocatellia; o__11-24; f__11-24; g__11-24 |
| 107234 | d__Archaea; p__Thermoproteota; c__Nitrososphaeria_A; o__Nitrososphaerales; f__Nitrososphaeraceae; g__*Nitrosocosmicus*; s__*Nitrosocosmicus hydrocola* | 0.743 | d__Archaea; p__Crenarchaeota; c__Nitrososphaeria; o__Nitrososphaerales; f__Nitrososphaeraceae |
| 225825 | d__Bacteria; p__Actinobacteriota; c__Actinomycetia; o__Mycobacteriales; f__Geodermatophilaceae; g__*Geodermatophilus_A* | 0.762 | d__Bacteria; p__Actinobacteriota; c__Actinobacteria; o__Frankiales; f__Geodermatophilaceae |
| 4417137 | d__Bacteria; p__Gemmatimonadota; c__Gemmatimonadetes; o__Gemmatimonadales; f__Gemmatimonadaceae; g__AG11; s__ | 1.000 | d__Bacteria; p__Gemmatimonadota; c__Gemmatimonadetes; o__Gemmatimonadales; f__Gemmatimonadaceae; g__uncultured |

| | | | |
|---|---|---|---|
| 859313 | d__Bacteria; p__Actinobacteriota; c__Acidimicrobiia_401430; o__Acidimicrobiales; f__Ilumatobacteraceae; g__*Ilumatobacter_A*; s__*Ilumatobacter_A coccineus*_A_400748 | 0.989 | d__Bacteria; p__Actinobacteriota; c__Acidimicrobiia; o__Microtrichales; f__Ilumatobacteraceae; g__*Ilumatobacter* |

f top 10 important features that were obtained
enGenes2 and SILVA, respectively. The table
prefix where ‘d’, ‘p’, ‘c’, ‘o’, ‘f’,’g’, and ‘s’
e taxonomic classifications may not be up to

| Confidence SILVA | Importance |
|---|---|
| 0.944 | 0.119 |
| 0.781 | 0.097 |
| 0.997 | 0.094 |
| 1.000 | 0.092 |

| | |
|---|---|
| 0.997 | 0.086 |
| 0.998 | 0.046 |
| 1.000 | 0.045 |
| 0.984 | 0.043 |
| 0.997 | 0.026 |

| | |
|---|---|
| 0.752 | 0.014 |
