## Supplementary Table 12 for "Grapevine Rootstock and Scion Genotypes’ Symbiosis with Soil Microbiome: A Machine Learning Revelation for Climate-Resilient Viticulture"

**Supplementary Table 12. Most informative ASVs for continent classification using ADA.** List
were obtained from Adaptive Boost (ADA) model. The taxonomy was then predicted with classifie
respectively. The table shows the predictions from both. The taxonomic level of the predictions are
‘c’, ‘o’, ‘f’,’g’, and ‘s’ are used for ‘kingdom’, ‘phylum’, ‘class’, ‘order’, ‘family’, ‘genus’, and ‘sp
classifications may not be up to the species level. The confidence scores from the classifier are dep

| ASV ID | Taxon GG | Confidence GG | Taxon SILVA |
|---|---|---|---|
| 818123 | d__Bacteria; p__Acidobacteriota; c__Blastocatellia; o__UBA7656; f__UBA7656; g__QHVH01; s__QHVH01 sp003222245 | 0.982 | d__Bacteria; p__Acidobacteriota; c__Blastocatellia; o__11-24; f__11-24; g__11-24 |
| 107234 | d__Archaea; p__Thermoproteota; c__Nitrososphaeria_A; o__Nitrososphaerales; f__Nitrososphaeraceae; g__*Nitrosocosmicus*; s__*Nitrosocosmicus hydrocola* | 0.743 | d__Archaea; p__Crenarchaeota; c__Nitrososphaeria; o__Nitrososphaerales; f__Nitrososphaeraceae |
| 859313 | d__Bacteria; p__Actinobacteriota; c__Acidimicrobiia_401430; o__Acidimicrobiales; f__Ilumatobacteraceae; g__*Ilumatobacter*_A; s__*Ilumatobacter*_A *coccineus*_A_400748 | 0.989 | d__Bacteria; p__Actinobacteriota; c__Acidimicrobiia; o__Microtrichales; f__Ilumatobacteraceae; g__*Ilumatobacter* |

| | | | |
|---|---|---|---|
| 1062101 | d__Bacteria; p__Verrucomicrobiota; c__Verrucomicrobiae; o__Chthoniobacterales; f__UBA10450 | 1.000 | d__Bacteria; p__Verrucomicrobiota; c__Verrucomicrobiae; o__Chthoniobacterales; f__Chthoniobacteraceae; g__*Candidatus_Udaeobacter* |
| 5d8b6db4f71e15746b9fe58 | d__Bacteria; p__Planctomycetota; c__Planctomycetia; o__Pirellulales | 0.993 | d__Bacteria; p__Planctomycetota; c__Planctomycetes; o__Pirellulales; f__Pirellulaceae; g__*Pirellula* |
| 144242 | d__Bacteria; p__Proteobacteria; c__Gammaproteobacteria; o__Burkholderiales_597441; f__SG8-41; g__UBA5216; s__UBA5216 sp902825845 | 0.898 | d__Bacteria; p__Proteobacteria; c__Gammaproteobacteria; o__Burkholderiales; f__TRA3-20; g__TRA3-20 |
| 210475 | d__Bacteria; p__Proteobacteria; c__Gammaproteobacteria; o__Burkholderiales_597439; f__Usitatibacteraceae; g__; s__ | 0.910 | d__Bacteria; p__Proteobacteria; c__Gammaproteobacteria; o__Burkholderiales; f__Nitrosomonadaceae; g__IS-44 |
| 589261 | d__Bacteria; p__Acidobacteriota; c__Thermoanaerobaculia; o__UBA5066; f__Gp7-AA6; g__Gp7-AA6; s__Gp7-AA6 sp003222385 | 1.000 | d__Bacteria; p__Acidobacteriota; c__Holophagae; o__Subgroup_7; f__Subgroup_7; g__Subgroup_7 |

| | | | |
|---|---|---|---|
| f985f67cbd4663c3696418d | d__Bacteria; p__Actinobacteriota; c__Thermoleophilia; o__Gaiellales; f__Gaiellaceae | 0.952 | d__Bacteria; p__Actinobacteriota; c__Thermoleophilia; o__Gaiellales; f__uncultured; g__uncultured |
| 3bf6ac8e5acf2a2360fa5762 | d__Bacteria; p__Proteobacteria; c__Gammaproteobacteria; o__Xanthomonadales_613062; f__Rhodanobacteraceae_613062 | 1.000 | d__Bacteria; p__Proteobacteria; c__Gammaproteobacteria; o__Xanthomonadales; f__Rhodanobacteraceae |

t of top 10 important features (ASVs) that
ers trained on GreenGenes2 and SILVA,
indicated with a letter prefix where ‘d’, ‘p’,
ecies’ respectively. The taxonomic
icted in the adjacent columns.

| Confidence SILVA | Importance |
|---|---|
| 0.998 | 0.302 |
| 1.000 | 0.215 |
| 0.752 | 0.177 |

| | |
|---|---|
| 0.995 | 0.072 |
| 0.972 | 0.048 |
| 0.873 | 0.035 |
| 0.997 | 0.024 |
| 0.998 | 0.016 |

| | |
|---|---|
| 0.989 | 0.016 |
| 1.000 | 0.015 |
