## Supplementary Table 13 for "Grapevine Rootstock and Scion Genotypes’ Symbiosis with Soil Microbiome: A Machine Learning Revelation for Climate-Resilient Viticulture"

**Supplementary Table 13. Most informative ASVs for continent classification using GBM.** Lis were obtained from Gradient Boost (GBM) model. The taxonomy was then predicted with classifie respectively. The table shows the predictions from both. The taxonomic level of the predictions are 'c', 'o', 'f','g', and 's' are used for 'kingdom', 'phylum', 'class', 'order', 'family', 'genus', and 'sp classifications may not be up to the species level. The confidence scores from the classifier are dep

| ASV ID | Taxon GG | Confidence GG | Taxon SILVA |
|---|---|---|---|
| 210475 | d__Bacteria; p__Proteobacteria; c__Gammaproteobacteria; o__Burkholderiales_597439; f__Usitatibacteraceae; g__; s__ | 0.909931985 | d__Bacteria; p__Proteobacteria; c__Gammaproteobacteria; o__Burkholderiales; f__Nitrosomonadaceae; g__IS-44 |
| 4417137 | d__Bacteria; p__Gemmatimonadota; c__Gemmatimonadetes; o__Gemmatimonadales; f__Gemmatimonadaceae; g__AG11; s__ | 0.999998729 | d__Bacteria; p__Gemmatimonadota; c__Gemmatimonadetes; o__Gemmatimonadales; f__Gemmatimonadaceae; g__uncultured |
| 136760 | d__Bacteria; p__Nitrospirota_A_437815; c__Nitrospiria; o__Nitrospirales; f__Nitrospiraceae; g__Nitrospira_C | 0.999970949 | d__Bacteria; p__Nitrospirota; c__Nitrospiria; o__Nitrospirales; f__Nitrospiraceae; g__*Nitrospira* ; s__*Nitrospira_japonica* |
| 263435d5671de6e03efe970 | d__Bacteria; p__Actinobacteriota; c__Actinomycetia; o__Mycobacteriales; f__Geodermatophilaceae | 0.983344733 | d__Bacteria; p__Actinobacteriota; c__Actinobacteria; o__Frankiales; f__Geodermatophilaceae |

| | | | |
|---|---|---|---|
| c0d81dd6449a43b34fcb184 | d__Archaea; p__Thermoproteota; c__Nitrososphaeria_A; o__Nitrososphaerales; f__Nitrososphaeraceae; g__*Nitrosocosmicus*; s__*Nitrosocosmicus hydrocola* | 0.755605591 | d__Archaea; p__Crenarchaeota; c__Nitrososphaeria; o__Nitrososphaerales; f__Nitrososphaeraceae |
| 225825 | d__Bacteria; p__Actinobacteriota; c__Actinomycetia; o__Mycobacteriales; f__Geodermatophilaceae; g__*Geodermatophilus*_A | 0.761706826 | d__Bacteria; p__Actinobacteriota; c__Actinobacteria; o__Frankiales; f__Geodermatophilaceae |
| 1091605 | d__Bacteria; p__Actinobacteriota; c__Acidimicrobiia_401430; o__Acidimicrobiales; f__Ilumatobacteraceae | 0.998972564 | d__Bacteria; p__Actinobacteriota; c__Acidimicrobiia; o__Microtrichales; f__Ilumatobacteraceae; g__uncultured |
| 769222 | d__Bacteria; p__Bacteroidota; c__Bacteroidia; o__Bacteroidales; f__Marinilabiliaceae; g__JC017; s__JC017 sp004296775 | 0.999591552 | d__Bacteria; p__Cyanobacteria; c__Cyanobacteriia; o__Chloroplast; f__Chloroplast; g__Chloroplast |
| 107234 | d__Archaea; p__Thermoproteota; c__Nitrososphaeria_A; o__Nitrososphaerales; f__Nitrososphaeraceae; g__*Nitrosocosmicus*; s__*Nitrosocosmicus hydrocola* | 0.742836131 | d__Archaea; p__Crenarchaeota; c__Nitrososphaeria; o__Nitrososphaerales; f__Nitrososphaeraceae |

| | | | |
|---|---|---|---|
| b10914468e29b7337f5fa25 | d__Archaea; p__Thermoproteota; c__Nitrososphaeria_A; o__Nitrososphaerales; f__Nitrososphaeraceae; g__*Nitrosocosmicus* | 0.999887031 | d__Archaea; p__Crenarchaeota; c__Nitrososphaeria; o__Nitrososphaerales; f__Nitrososphaeraceae; g__*Nitrososphaeraceae* |

t of top 10 important features (ASVs) that
ers trained on GreenGenes2 and SILVA,
indicated with a letter prefix where ‘d’, ‘p’,
ecies’ respectively. The taxonomic
icted in the adjacent columns.

| Confidence SILVA | Importance |
|---|---|
| 0.99726338 | 0.24530146 |
| 0.99730609 | 0.13310921 |
| 0.94425584 | 0.06788043 |
| 0.95431377 | 0.06408626 |

| | |
|---|---|
| 0.99999868 | 0.04667511 |
| 0.98399053 | 0.04550395 |
| 0.81499884 | 0.02835947 |
| 0.99999925 | 0.0225382 |
| 0.99999947 | 0.01829339 |

| | |
|---|---|
| 0.80122517 | 0.0172962 |
