## Supplementary Table 14 for "Grapevine Rootstock and Scion Genotypes’ Symbiosis with Soil Microbiome: A Machine Learning Revelation for Climate-Resilient Viticulture"

**Supplementary Table 14. Most informative ASVs for country classification using RF.** List of obtained from Random Forests (RF) model. The taxonomy was then predicted with classifiers trair The table shows the predictions from both. The taxonomic level of the predictions are indicated wi and ‘s’ are used for ‘kingdom’, ‘phylum’, ‘class’, ‘order’, ‘family’, ‘genus’, and ‘species’ respectiv up to the species level. The confidence scores from the classifier are depicted in the adjacent colum

| ASV ID | Taxon GG | Confidence GG | Taxon SILVA |
|---|---|---|---|
| 6e093dc55c94afd3d6c3eb4 | d__Bacteria; p__Proteobacteria; c__Alphaproteobacteria; o__Rhizobiales_A_504705; f__Xanthobacteraceae_503485; g__VAZQ01; s__VAZQ01 sp005883115 | 0.961 | d__Bacteria; p__Proteobacteria; c__Alphaproteobacteria; o__Rhizobiales; f__Xanthobacteraceae; g__uncultured |
| 4362620 | d__Archaea; p__Thermoproteota; c__Nitrososphaeria_A; o__Nitrososphaerales; f__Nitrososphaeraceae; g__*Nitrosocosmicus*; s__*Nitrosocosmicus hydrocola* | 0.872 | d__Archaea; p__Crenarchaeota; c__Nitrososphaeria; o__Nitrososphaerales; f__Nitrososphaeraceae; g__*Candidatus_Nitrocosmicus* |
| 44ce6482692430da34be7e4 | d__Bacteria; p__Acidobacteriota; c__Vicinamibacteria; o__Vicinamibacterales | 0.986 | d__Bacteria; p__Acidobacteriota; c__Vicinamibacteria; o__Vicinamibacterales; f__Vicinamibacteraceae; g__*Vicinamibacteraceae* |
| 136760 | d__Bacteria; p__Nitrospirota_A_437815; c__Nitrospiria; o__Nitrospirales; f__Nitrospiraceae; g__*Nitrospira*_C | 1.000 | d__Bacteria; p__Nitrospirota; c__Nitrospiria; o__Nitrospirales; f__Nitrospiraceae; g__*Nitrospira*; s__*Nitrospira_japonica* |

| | | | |
|---|---|---|---|
| 575740 | d__Bacteria; p__Acidobacteriota; c__Vicinamibacteria; o__Vicinamibacterales; f__UBA2999; g__WHSN01; s__ | 0.860 | d__Bacteria; p__Acidobacteriota; c__Vicinamibacteria; o__Vicinamibacterales; f__uncultured; g__uncultured |
| 8efbdf43cd6d5e499ec3ceac | d__Bacteria; p__Proteobacteria; c__Alphaproteobacteria; o__Rhizobiales_A_504705; f__Xanthobacteraceae_503485; g__*Bradyrhizobium* | 0.845 | d__Bacteria; p__Proteobacteria; c__Alphaproteobacteria; o__Rhizobiales; f__Xanthobacteraceae; g__*Bradyrhizobium* |
| 1efc4ad517d4cd1fff23f0fd8 | d__Bacteria; p__Proteobacteria; c__Gammaproteobacteria; o__Burkholderiales_592522; f__Burkholderiaceae_A_592522 | 0.999 | d__Bacteria; p__Proteobacteria; c__Gammaproteobacteria; o__Burkholderiales; f__Comamonadaceae |
| 210475 | d__Bacteria; p__Proteobacteria; c__Gammaproteobacteria; o__Burkholderiales_597439; f__Usitatibacteraceae; g__; s__ | 0.910 | d__Bacteria; p__Proteobacteria; c__Gammaproteobacteria; o__Burkholderiales; f__Nitrosomonadaceae; g__IS-44 |
| ed3d7020f31a64a765244b2 | d__Archaea; p__Thermoproteota; c__Nitrososphaeria_A; o__Nitrososphaerales; f__Nitrososphaeraceae; g__*Nitrososphaera* | 0.899 | d__Archaea; p__Crenarchaeota; c__Nitrososphaeria; o__Nitrososphaerales; f__Nitrososphaeraceae; g__*Nitrososphaeraceae* |
| 77a56d88843bb75cbd7f75f | d__Bacteria; p__Actinobacteriota; c__Actinomycetia; o__Actinomycetales; f__Micrococcaceae | 1.000 | d__Bacteria; p__Actinobacteriota; c__Actinobacteria; o__Micrococcales; f__Micrococcaceae |

top 10 important features (ASVs) that were ned on GreenGenes2 and SILVA, respectively. th a letter prefix where ‘d’, ‘p’, ‘c’, ‘o’, ‘f’,’g’, ely. The taxonomic classifications may not be ns.

| Confidence SILVA | Importance |
|---|---|
| 0.712 | 0.144 |
| 0.703 | 0.113 |
| 0.960 | 0.091 |
| 0.944 | 0.058 |

| | |
|---|---|
| 0.997 | 0.054 |
| 0.712 | 0.037 |
| 1.000 | 0.029 |
| 0.997 | 0.026 |
| 0.921 | 0.025 |
| 1.000 | 0.022 |
