## Supplementary Table 15 for "Grapevine Rootstock and Scion Genotypes’ Symbiosis with Soil Microbiome: A Machine Learning Revelation for Climate-Resilient Viticulture"

**Supplementary Table 15. Most informative ASVs for country classification using ADA.** List [illegible]
obtained from Adaptive Boost (ADA) model. The taxonomy was then predicted with classifiers tra[illegible]
respectively. The table shows the predictions from both. The taxonomic level of the predictions are[illegible]
'c', 'o', 'f','g', and 's' are used for 'kingdom', 'phylum', 'class', 'order', 'family', 'genus', and 'sp[illegible]
classifications may not be up to the species level. The confidence scores from the classifier are dep[illegible]

| ASV ID | Taxon GG | Confidence GG | Taxon SILVA |
|---|---|---|---|
| 4ad517d4cd1fff23f0fd84c0[illegible] | d__Bacteria; p__Proteobacteria; c__Gammaproteobacteria; o__Burkholderiales_592522; f__Burkholderiaceae_A_592522 | 0.999 | d__Bacteria; p__Proteobacteria; c__Gammaproteobacteria; o__Burkholderiales; f__Comamonadaceae |
| 4362620 | d__Archaea; p__Thermoproteota; c__Nitrososphaeria_A; o__Nitrososphaerales; f__Nitrososphaeraceae; g__*Nitrosocosmicus*; s__Nitrosocosmicus hydrocola | 0.872 | d__Archaea; p__Crenarchaeota; c__Nitrososphaeria; o__Nitrososphaerales; f__Nitrososphaeraceae; g__*Candidatus_Nitrocosmicus* |
| [illegible]482692430da34be7e4359b[illegible] | d__Bacteria; p__Acidobacteriota; c__Vicinamibacteria; o__Vicinamibacterales | 0.986 | d__Bacteria; p__Acidobacteriota; c__Vicinamibacteria; o__Vicinamibacterales; f__Vicinamibacteraceae; g__*Vicinamibacteraceae* |
| 136760 | d__Bacteria; p__Nitrospirota_A_437815; c__Nitrospiria; o__Nitrospirales; f__Nitrospiraceae; g__*Nitrospira_C* | 1.000 | d__Bacteria; p__Nitrospirota; c__Nitrospiria; o__Nitrospirales; f__Nitrospiraceae; g__*Nitrospira*; s__*Nitrospira_japonica* |

| | | | |
|---|---|---|---|
| 3dc55c94afd3d6c3eb45929 | d__Bacteria; p__Proteobacteria; c__Alphaproteobacteria; o__Rhizobiales_A_504705; f__Xanthobacteraceae_503485; g__VAZQ01; s__VAZQ01 sp005883115 | 0.961 | d__Bacteria; p__Proteobacteria; c__Alphaproteobacteria; o__Rhizobiales; f__Xanthobacteraceae; g__uncultured |
| 5d8b6db4f71e15746b9fe58 | d__Bacteria; p__Planctomycetota; c__Planctomycetia; o__Pirellulales | 0.993 | d__Bacteria; p__Planctomycetota; c__Planctomycetes; o__Pirellulales; f__Pirellulaceae; g__*Pirellula* |
| 212493 | d__Bacteria; p__Acidobacteriota; c__Acidobacteriae; o__Acidobacteriales; f__SbA1 | 1.000 | d__Bacteria; p__Acidobacteriota; c__Acidobacteriae; o__Acidobacteriales; f__uncultured; g__uncultured |
| 773596 | d__Bacteria; p__Proteobacteria; c__Gammaproteobacteria; o__Steroidobacterales; f__Steroidobacteraceae; g__*Povalibacter; s* __ | 0.746 | d__Bacteria; p__Proteobacteria; c__Gammaproteobacteria; o__Steroidobacterales; f__Steroidobacteraceae; g__*Steroidobacter* |
| 1f3d5d896228e792535ae41 | d__Bacteria; p__Desulfobacterota_B; c__Binatia; o__UBA9968; f__UBA9968; g__; s__ | 0.722 | d__Bacteria |
| 697344 | d__Bacteria; p__Acidobacteriota; c__Acidobacteriae; o__Bryobacterales; f__Bryobacteraceae; g__KBS-96; s__KBS-96 sp000381625 | 1.000 | d__Bacteria; p__Acidobacteriota; c__Acidobacteriae; o__Bryobacterales; f__Bryobacteraceae; g__*Bryobacter* |

of top 10 important features (ASVs) that were
ained on GreenGenes2 and SILVA,
e indicated with a letter prefix where ‘d’, ‘p’,
ecies’ respectively. The taxonomic
icted in the adjacent columns.

| Confidence SILVA | Importance |
|---|---|
| 1.000 | 0.223 |
| 0.703 | 0.190 |
| 0.960 | 0.175 |
| 0.944 | 0.158 |

| | |
|---|---|
| 0.712 | 0.047 |
| 0.972 | 0.031 |
| 0.999 | 0.029 |
| 0.729 | 0.021 |
| 1.000 | 0.018 |
| 0.993 | 0.018 |
