## Supplementary Table 16 for "Grapevine Rootstock and Scion Genotypes’ Symbiosis with Soil Microbiome: A Machine Learning Revelation for Climate-Resilient Viticulture"

**Supplementary Table 16. Most informative ASVs for country classification using GBM.** List obtained from Gradient Boost (GBM) model. The taxonomy was then predicted with classifiers tra respectively. The table shows the predictions from both. The taxonomic level of the predictions are 'c', 'o', 'f','g', and 's' are used for 'kingdom', 'phylum', 'class', 'order', 'family', 'genus', and 'sp classifications may not be up to the species level. The confidence scores from the classifier are dep

| ASV ID | Taxon GG | Confidence GG | Taxon SILVA |
|---|---|---|---|
| 4417137 | d__Bacteria; p__Gemmatimonadota; c__Gemmatimonadetes; o__Gemmatimonadales; f__Gemmatimonadaceae; g__AG11; s__ | 1.000 | d__Bacteria; p__Gemmatimonadota; c__Gemmatimonadetes; o__Gemmatimonadales; f__Gemmatimonadaceae; g__uncultured |
| 5d88843bb75cbd7f75f1f130 | d__Bacteria; p__Actinobacteriota; c__Actinomycetia; o__Actinomycetales; f__Micrococcaceae | 1.000 | d__Bacteria; p__Actinobacteriota; c__Actinobacteria; o__Micrococcales; f__Micrococcaceae |
| 482692430da34be7e4359b | d__Bacteria; p__Acidobacteriota; c__Vicinamibacteria; o__Vicinamibacterales | 0.986 | d__Bacteria; p__Acidobacteriota; c__Vicinamibacteria; o__Vicinamibacterales; f__Vicinamibacteraceae; g__*Vicinamibacteraceae* |
| df43cd6d5e499ec3ceaccf75 | d__Bacteria; p__Proteobacteria; c__Alphaproteobacteria; o__Rhizobiales_A_504705; f__Xanthobacteraceae_503485; g__*Bradyrhizobium* | 0.845 | d__Bacteria; p__Proteobacteria; c__Alphaproteobacteria; o__Rhizobiales; f__Xanthobacteraceae; g__*Bradyrhizobium* |

| | | | |
|---|---|---|---|
| 225825 | d__Bacteria; p__Actinobacteriota; c__Actinomycetia; o__Mycobacteriales; f__Geodermatophilaceae; g__*Geodermatophilus*_A | 0.762 | d__Bacteria; p__Actinobacteriota; c__Actinobacteria; o__Frankiales; f__Geodermatophilaceae |
| 575740 | d__Bacteria; p__Acidobacteriota; c__Vicinamibacteria; o__Vicinamibacterales; f__UBA2999; g__WHSN01; s__ | 0.860 | d__Bacteria; p__Acidobacteriota; c__Vicinamibacteria; o__Vicinamibacterales; f__uncultured; g__uncultured |
| 1091605 | d__Bacteria; p__Actinobacteriota; c__Acidimicrobiia_401430; o__Acidimicrobiales; f__Ilumatobacteraceae | 0.999 | d__Bacteria; p__Actinobacteriota; c__Acidimicrobiia; o__Microtrichales; f__Ilumatobacteraceae; g__uncultured |
| 210475 | d__Bacteria; p__Proteobacteria; c__Gammaproteobacteria; o__Burkholderiales_597439; f__Usitatibacteraceae; g__; s__ | 0.910 | d__Bacteria; p__Proteobacteria; c__Gammaproteobacteria; o__Burkholderiales; f__Nitrosomonadaceae; g__IS-44 |
| [illegible]b10914468e29b7337f5fa25[illegible] | d__Archaea; p__Thermoproteota; c__Nitrososphaeria_A; o__Nitrososphaerales; f__Nitrososphaeraceae; g__*Nitrosocosmicus* | 1.000 | d__Archaea; p__Crenarchaeota; c__Nitrososphaeria; o__Nitrososphaerales; f__Nitrososphaeraceae; g__*Nitrososphaeraceae* |
| [illegible]263435d5671de6e03efe970[illegible] | d__Bacteria; p__Actinobacteriota; c__Actinomycetia; o__Mycobacteriales; f__Geodermatophilaceae | 0.983 | d__Bacteria; p__Actinobacteriota; c__Actinobacteria; o__Frankiales; f__Geodermatophilaceae |

of top 10 important features (ASVs) that were
ined on GreenGenes2 and SILVA,
indicated with a letter prefix where ‘d’, ‘p’,
ecies’ respectively. The taxonomic
icted in the adjacent columns.

| Confidence SILVA | Importance |
|---|---|
| 0.997 | 0.126 |
| 1.000 | 0.112 |
| 0.960 | 0.089 |
| 0.712 | 0.059 |

| | |
|---|---|
| 0.984 | 0.053 |
| 0.997 | 0.052 |
| 0.815 | 0.046 |
| 0.997 | 0.041 |
| 0.801 | 0.029 |
| 0.954 | 0.021 |
