## Supplementary Table 17 for "Grapevine Rootstock and Scion Genotypes’ Symbiosis with Soil Microbiome: A Machine Learning Revelation for Climate-Resilient Viticulture"

**Supplementary Table 17. Most informative ASVs for cultivar classification using RF.** List of obtained from Random Forest (RF) model. The taxonomy was then predicted with classifiers traine The table shows the predictions from both. The taxonomic level of the predictions are indicated wi and ‘s’ are used for ‘kingdom’, ‘phylum’, ‘class’, ‘order’, ‘family’, ‘genus’, and ‘species’ respectiv up to the species level. The confidence scores from the classifier are depicted in the adjacent colum

| ASV ID | Taxon GG | Confidence GG | Taxon SILVA |
|---|---|---|---|
| 136760 | d__Bacteria; p__Nitrospirota_A_437815; c__Nitrospiria; o__Nitrospirales; f__Nitrospiraceae; g__*Nitrospira* _C | 1.000 | d__Bacteria; p__Nitrospirota; c__Nitrospiria; o__Nitrospirales; f__Nitrospiraceae; g__*Nitrospira* ; s__*Nitrospira _japonica* |
| 4362620 | d__Archaea; p__Thermoproteota; c__Nitrososphaeria_A; o__Nitrososphaerales; f__Nitrososphaeraceae; g__*Nitrosocosmicus* ; s__*Nitrosocosmicus hydrocola* | 0.872 | d__Archaea; p__Crenarchaeota; c__Nitrososphaeria; o__Nitrososphaerales; f__Nitrososphaeraceae; g__*Candidatus _Nitrocosmicus* |
| ebb75a37c5b06ba598a461fb | d__Bacteria; p__Actinobacteriota; c__Rubrobacteria; o__Rubrobacterales; f__Rubrobacteraceae; g__; s__ | 0.940 | d__Bacteria; p__Actinobacteriota; c__Rubrobacteria; o__Rubrobacterales; f__Rubrobacteriaceae; g__*Rubrobacter* |
| 78821a8093d440270f0c873 | d__Bacteria; p__Bacteroidota; c__Bacteroidia; o__Cytophagales; f__Hymenobacteraceae; g__*Adhaeribacter* ; s__*Adhaeribacter rhizoryzae* | 0.713 | d__Bacteria; p__Bacteroidota; c__Bacteroidia; o__Cytophagales; f__Hymenobacteraceae; g__*Adhaeribacter* |

| | | | |
|---|---|---|---|
| 916442 | d__Bacteria; p__Actinobacteriota; c__Thermoleophilia; o__Solirubrobacterales; f__Parviterribacteraceae; g__*Parviterribacter* | 0.839 | d__Bacteria; p__Actinobacteriota; c__Thermoleophilia; o__Solirubrobacterales; f__Solirubrobacteraceae; g__Parviterribacter; s__uncultured_bacterium |
| 818123 | d__Bacteria; p__Acidobacteriota; c__Blastocatellia; o__UBA7656; f__UBA7656; g__QHVH01; s__QHVH01 sp003222245 | 0.982 | d__Bacteria; p__Acidobacteriota; c__Blastocatellia; o__11-24; f__11-24; g__11-24 |
| 1091605 | d__Bacteria; p__Actinobacteriota; c__Acidimicrobiia_401430; o__Acidimicrobiales; f__Ilumatobacteraceae | 0.999 | d__Bacteria; p__Actinobacteriota; c__Acidimicrobiia; o__Microtrichales; f__Ilumatobacteraceae; g__uncultured |
| c0d81dd6449a43b34fcb184[illegible] | d__Archaea; p__Thermoproteota; c__Nitrososphaeria_A; o__Nitrososphaerales; f__Nitrososphaeraceae; g__*Nitrosocosmicus*; s__*Nitrosocosmicus hydrocola* | 0.756 | d__Archaea; p__Crenarchaeota; c__Nitrososphaeria; o__Nitrososphaerales; f__Nitrososphaeraceae |
| 210475 | d__Bacteria; p__Proteobacteria; c__Gammaproteobacteria; o__Burkholderiales_597439; f__Usitatibacteraceae; g__; s__ | 0.910 | d__Bacteria; p__Proteobacteria; c__Gammaproteobacteria; o__Burkholderiales; f__Nitrosomonadaceae; g__IS-44 |

| | | | |
|---|---|---|---|
| 95d9acfe4501a83156f0b941 | d__Bacteria; p__Proteobacteria; c__Alphaproteobacteria; o__Geminicoccales; f__Geminicoccaceae; g__HRBIN40; s__HRBIN40 sp002898275 | 0.988 | d__Bacteria; p__Proteobacteria; c__Alphaproteobacteria; o__Tistrellales; f__Geminicoccaceae; g__*Candidatus _Alysiosphaera* |

top 10 important features (ASVs) that were
ed on GreenGenes2 and SILVA, respectively.
th a letter prefix where ‘d’, ‘p’, ‘c’, ‘o’, ‘f’,’g’,
ely. The taxonomic classifications may not be
ns.

| Confidence SILVA | Importance |
|---|---|
| 0.944 | 0.100 |
| 0.703 | 0.098 |
| 1.000 | 0.067 |
| 0.999 | 0.047 |

| | |
|---|---|
| 0.781 | 0.034 |
| 0.998 | 0.027 |
| 0.815 | 0.021 |
| 1.000 | 0.019 |
| 0.997 | 0.018 |

| | |
|---|---|
| 0.999 | 0.017 |
