## Supplementary Table 18 for "Grapevine Rootstock and Scion Genotypes’ Symbiosis with Soil Microbiome: A Machine Learning Revelation for Climate-Resilient Viticulture"

**Supplementary Table 18. Most informative ASVs for cultivar classification using ADA.** List c
obtained from Adaptive Boost (ADA) model. The taxonomy was then predicted with classifiers tra
respectively. The table shows the predictions from both. The taxonomic level of the predictions are
'c', 'o', 'f','g', and 's' are used for 'kingdom', 'phylum', 'class', 'order', 'family', 'genus', and 'sp
classifications may not be up to the species level. The confidence scores from the classifier are dep

| ASV ID | Taxon GG | Confidence GG | Taxon SILVA |
|---|---|---|---|
| f98debb75a37c5b06ba598a461fbd1bf | d__Bacteria; p__Actinobacteriota; c__Rubrobacteria; o__Rubrobacterales; f__Rubrobacteraceae; g__; s__ | 0.940 | d__Bacteria; p__Actinobacteriota; c__Rubrobacteria; o__Rubrobacterales; f__Rubrobacteriaceae; g__*Rubrobacter* |
| 4362620 | d__Archaea; p__Thermoproteota; c__Nitrososphaeria_A; o__Nitrososphaerales; f__Nitrososphaeraceae; g__*Nitrosocosmicus*; s__*Nitrosocosmicus hydrocola* | 0.872 | d__Archaea; p__Crenarchaeota; c__Nitrososphaeria; o__Nitrososphaerales; f__Nitrososphaeraceae; g__*Candidatus*_*Nitrocosmicus* |
| 136760 | d__Bacteria; p__Nitrospirota_A_437815; c__Nitrospiria; o__Nitrospirales; f__Nitrospiraceae; g__*Nitrospira*_C | 1.000 | d__Bacteria; p__Nitrospirota; c__Nitrospiria; o__Nitrospirales; f__Nitrospiraceae; g__*Nitrospira*; s__*Nitrospira*_*japonica* |
| 916442 | d__Bacteria; p__Actinobacteriota; c__Thermoleophilia; o__Solirubrobacterales; f__Parviterribacteraceae; g__*Parviterribacter* | 0.839 | d__Bacteria; p__Actinobacteriota; c__Thermoleophilia; o__Solirubrobacterales; f__Solirubrobacteraceae; g__*Parviterribacter*; s__uncultured_bacterium |

| | | | |
|---|---|---|---|
| 09a802948f82a388b8a9522a8887f3cd | d__Bacteria; p__Acidobacteriota; c__Blastocatellia; o__Pyrinomonadales; f__Pyrinomonadaceae_433871; g__PSRF01; s__ | 1.000 | d__Bacteria; p__Acidobacteriota; c__Blastocatellia; o__Pyrinomonadales; f__Pyrinomonadaceae; g__RB41 |
| 9aafc0120a37d269baf7c93cab16dfde | d__Bacteria; p__Verrucomicrobiota; c__Verrucomicrobiae; o__Pedosphaerales | 0.804 | d__Bacteria; p__Verrucomicrobiota; c__Verrucomicrobiae; o__Pedosphaerales; f__Pedosphaeraceae; g__*Pedosphaeraceae*; s__uncultured_bacterium |
| 2e83ec04e6a19cea2b2e59ca62676d53 | d__Bacteria; p__Desulfobacterota_B; c__Binatia; o__Bin18; f__Bin18; g__JABFSC01; s__ | 0.999 | d__Bacteria; p__Entotheonellaeota; c__Entotheonellia; o__Entotheonellales; f__Entotheonellaceae; g__*Entotheonellaceae* |
| 1073d92794bbabefa063efc7e17da64d | d__Bacteria; p__Acidobacteriota; c__Acidobacteriae; o__Acidobacteriales; f__Koribacteraceae; g__*Koribacter*; s__*Koribacter versatilis*_A | 0.994 | d__Bacteria; p__Acidobacteriota; c__Acidobacteriae; o__Acidobacteriales; f__Koribacteraceae; g__*Candidatus_Koribacter*; s__uncultured_bacterium |
| 1f01e277aeb09e117b913d10d8de6de5 | d__Bacteria; p__Proteobacteria; c__Gammaproteobacteria; o__Pseudomonadales_660879; f__Moraxellaceae; g__*Acinetobacter* | 1.000 | d__Bacteria; p__Proteobacteria; c__Gammaproteobacteria; o__Pseudomonadales; f__Moraxellaceae; g__*Acinetobacter* |

| | | | |
|---|---|---|---|
| 773596 | d__Bacteria; p__Proteobacteria; c__Gammaproteobacteria; o__Steroidobacterales; f__Steroidobacteraceae; g__*Povalibacter*; s__ | 0.746 | d__Bacteria; p__Proteobacteria; c__Gammaproteobacteria; o__Steroidobacterales; f__Steroidobacteraceae; g__*Steroidobacter* |

of top 10 important features (ASVs) that were
ined on GreenGenes2 and SILVA,
indicated with a letter prefix where ‘d’, ‘p’,
ecies’ respectively. The taxonomic
icted in the adjacent columns.

| Confidence SILVA | Importance |
|---|---|
| 1.000 | 0.156 |
| 0.703 | 0.155 |
| 0.944 | 0.151 |
| 0.781 | 0.118 |

| | |
|---|---|
| 1.000 | 0.036 |
| 0.868 | 0.031 |
| 1.000 | 0.030 |
| 0.837 | 0.025 |
| 1.000 | 0.024 |

| | |
|---|---|
| 0.729 | 0.022 |
