## Supplementary Table 19 for "Grapevine Rootstock and Scion Genotypes’ Symbiosis with Soil Microbiome: A Machine Learning Revelation for Climate-Resilient Viticulture"

**Supplementary Table 19. Most informative ASVs for cultivar classification using GBM.** List
obtained from Gradient Boost (GBM) model. The taxonomy was then predicted with classifiers tra
respectively. The table shows the predictions from both. The taxonomic level of the predictions are
'c', 'o', 'f','g', and 's' are used for 'kingdom', 'phylum', 'class', 'order', 'family', 'genus', and 'sp
classifications may not be up to the species level. The confidence scores from the classifier are dep

| ASV ID | Taxon GG | Confidence GG | Taxon SILVA |
|---|---|---|---|
| 818123 | d__Bacteria; p__Acidobacteriota; c__Blastocatellia; o__UBA7656; f__UBA7656; g__QHVH01; s__QHVH01 sp003222245 | 0.982 | d__Bacteria; p__Acidobacteriota; c__Blastocatellia; o__11-24; f__11-24; g__11-24 |
| 136760 | d__Bacteria; p__Nitrospirota_A_437815; c__Nitrospiria; o__Nitrospirales; f__Nitrospiraceae; g__*Nitrospira* _C | 1.000 | d__Bacteria; p__Nitrospirota; c__Nitrospiria; o__Nitrospirales; f__Nitrospiraceae; g__*Nitrospira* ; s__*Nitrospira _japonica* |
| f98debb75a37c5b06ba598a461fbd1bf | d__Bacteria; p__Actinobacteriota; c__Rubrobacteria; o__Rubrobacterales; f__Rubrobacteraceae; g__; s__ | 0.940 | d__Bacteria; p__Actinobacteriota; c__Rubrobacteria; o__Rubrobacterales; f__Rubrobacteriaceae; g__*Rubrobacter* |
| 1091605 | d__Bacteria; p__Actinobacteriota; c__Acidimicrobiia_401430; o__Acidimicrobiales; f__Ilumatobacteraceae | 0.999 | d__Bacteria; p__Actinobacteriota; c__Acidimicrobiia; o__Microtrichales; f__Ilumatobacteraceae; g__uncultured |
| 7406b09d2022974338c18e894f589425 | d__Bacteria; p__Proteobacteria; c__Alphaproteobacteria; o__Sphingomonadales; f__Sphingomonadaceae; g__*Sphingomicrobium* _483265 | 0.989 | d__Bacteria; p__Proteobacteria; c__Alphaproteobacteria; o__Sphingomonadales; f__Sphingomonadaceae; g__*Sphingomonas* |

| | | | |
|---|---|---|---|
| 217604 | d__Bacteria; p__Actinobacteriota; c__Actinomycetia; o__Mycobacteriales; f__Geodermatophilaceae; g__Blastococcus | 0.989 | d__Bacteria; p__Actinobacteriota; c__Actinobacteria; o__Frankiales; f__Geodermatophilaceae; g__*Blastococcus* |
| 96839 | d__Bacteria; p__Proteobacteria; c__Gammaproteobacteria; o__Xanthomonadales_616009; f__Xanthomonadaceae_616009; g__*Lysobacter* _A_615995 | 0.995 | d__Bacteria; p__Proteobacteria; c__Gammaproteobacteria; o__Xanthomonadales; f__Xanthomonadaceae; g__*Lysobacter* |
| 82fcfd09db3347c7600caba70d032e4a | d__Bacteria; p__Proteobacteria; c__Gammaproteobacteria; o__Burkholderiales_597433; f__SG8-39; g__SCGC-AG-212-J23; s__SCGC-AG-212-J23 sp005881595 | 0.972 | d__Bacteria; p__Proteobacteria; c__Gammaproteobacteria; o__Burkholderiales; f__Nitrosomonadaceae; g__MND1 |
| 96a1c701a49759c0d76762e249ac8bf2 | d__Bacteria; p__Proteobacteria; c__Alphaproteobacteria; o__Rhizobiales_A_504705; f__Xanthobacteraceae_503485; g__*Bradyrhizobium* | 0.782 | d__Bacteria; p__Proteobacteria; c__Alphaproteobacteria; o__Rhizobiales; f__Xanthobacteraceae |
| 7168e518f75c5361d801cbe0e2c37d6c | d__Archaea; p__Thermoproteota; c__Nitrososphaeria_A; o__Nitrososphaerales; f__Nitrososphaeraceae; g__UBA10452; s__UBA10452 sp003176995 | 0.779 | d__Archaea; p__Crenarchaeota; c__Nitrososphaeria; o__Nitrososphaerales; f__Nitrososphaeraceae; g__*Nitrososphaeraceae* |

of top 10 important features (ASVs) that were
ined on GreenGenes2 and SILVA,
indicated with a letter prefix where ‘d’, ‘p’,
ecies’ respectively. The taxonomic
icted in the adjacent columns.

| Confidence SILVA | Importance |
|---|---|
| 0.998 | 0.125 |
| 0.944 | 0.123 |
| 1.000 | 0.085 |
| 0.815 | 0.076 |
| 0.997 | 0.042 |

| | |
|---|---|
| 0.990 | 0.029 |
| 0.995 | 0.026 |
| 1.000 | 0.019 |
| 1.000 | 0.016 |
| 0.986 | 0.015 |
