## Supplementary Table 20 for "Grapevine Rootstock and Scion Genotypes’ Symbiosis with Soil Microbiome: A Machine Learning Revelation for Climate-Resilient Viticulture"

**Supplementary Table 20. Overlapping Features for Continent Classification. List of features (A**
The taxonomy of the features was predicted with classifiers trained on GreenGenes2 and SILVA. The
'o', 'f','g', and 's' are used for 'kingdom', 'phylum', 'class', 'order', 'family', 'genus', and 'species' r
confidence scores from the classifier are depicted in the adjacent columns.

| ID | Taxon GG | Confidence GG | Taxon SILVA |
|---|---|---|---|
| 07525d8b6db4f71e15746b9fe58f1c21 | d__Bacteria; p__Planctomycetota; c__Planctomycetia; o__Pirellulales | 0.993 | d__Bacteria; p__Planctomycetota; c__Planctomycetes; o__Pirellulales; f__Pirellulaceae; g__*Pirellula* |
| 136760 | d__Bacteria; p__Nitrospirota_A_437815; c__Nitrospiria; o__Nitrospirales; f__Nitrospiraceae; g__Nitrospira_C | 1.000 | d__Bacteria; p__Nitrospirota; c__Nitrospiria; o__Nitrospirales; f__Nitrospiraceae; g__Nitrospira; s__Nitrospira_japonica |
| 1efc4ad517d4cd1fff23f0fd84c0eaa6 | d__Bacteria; p__Proteobacteria; c__Gammaproteobacteria; o__Burkholderiales_592522; f__Burkholderiaceae_A_592522 | 0.999 | d__Bacteria; p__Proteobacteria; c__Gammaproteobacteria; o__Burkholderiales; f__Comamonadaceae |
| 212493 | d__Bacteria; p__Acidobacteriota; c__Acidobacteriae; o__Acidobacteriales; f__SbA1 | 1.000 | d__Bacteria; p__Acidobacteriota; c__Acidobacteriae; o__Acidobacteriales; f__uncultured; g__uncultured |
| 2de2695189645de769b2be0b536c2106 | d__Bacteria; p__Proteobacteria; c__Alphaproteobacteria; o__Sphingomonadales; f__Sphingomonadaceae; g__Novosphingobium_485351 | 0.969 | d__Bacteria; p__Proteobacteria; c__Alphaproteobacteria; o__Sphingomonadales; f__Sphingomonadaceae; g__Novosphingobium |

| | | | |
|---|---|---|---|
| 4362620 | d__Archaea; p__Thermoproteota; c__Nitrososphaeria_A; o__Nitrososphaerales; f__Nitrososphaeraceae; g__Nitrosocosmicus; s__Nitrosocosmicus hydrocola | 0.872 | d__Archaea; p__Crenarchaeota; c__Nitrososphaeria; o__Nitrososphaerales; f__Nitrososphaeraceae; g__Candidatus_Nitrocosmicus |
| 44ce6482692430da34be7e4359b43d08 | d__Bacteria; p__Acidobacteriota; c__Vicinamibacteria; o__Vicinamibacterales | 0.986 | d__Bacteria; p__Acidobacteriota; c__Vicinamibacteria; o__Vicinamibacterales; f__Vicinamibacteraceae; g__Vicinamibacteraceae |
| 4c687e9abe11199b993e2ab668384d5c | d__Bacteria; p__Nitrospirota_A_437815; c__Nitrospiria; o__Nitrospirales; f__Nitrospiraceae; g__Nitrospira_C | 1.000 | d__Bacteria; p__Nitrospirota; c__Nitrospiria; o__Nitrospirales; f__Nitrospiraceae; g__Nitrospira; s__Nitrospira_japonica |
| 575740 | d__Bacteria; p__Acidobacteriota; c__Vicinamibacteria; o__Vicinamibacterales; f__UBA2999; g__WHSN01; s__ | 0.860 | d__Bacteria; p__Acidobacteriota; c__Vicinamibacteria; o__Vicinamibacterales; f__uncultured; g__uncultured |
| 6e093dc55c94afd3d6c3eb459297f776 | d__Bacteria; p__Proteobacteria; c__Alphaproteobacteria; o__Rhizobiales_A_504705; f__Xanthobacteraceae_503485; g__VAZQ01; s__VAZQ01 sp005883115 | 0.961 | d__Bacteria; p__Proteobacteria; c__Alphaproteobacteria; o__Rhizobiales; f__Xanthobacteraceae; g__uncultured |

| | | | |
|---|---|---|---|
| 769222 | d__Bacteria; p__Bacteroidota; c__Bacteroidia; o__Bacteroidales; f__Marinilabiliaceae; g__JC017; s__JC017 sp004296775 | 1.000 | d__Bacteria; p__Cyanobacteria; c__Cyanobacteriia; o__Chloroplast; f__Chloroplast; g__Chloroplast |
| 888d0c20bfb737e06356e46b41352a40 | d__Bacteria; p__Proteobacteria; c__Alphaproteobacteria; o__Sphingomonadales; f__Sphingomonadaceae; g__Sphingomonas_L_486704 | 0.984 | d__Bacteria; p__Proteobacteria; c__Alphaproteobacteria; o__Sphingomonadales; f__Sphingomonadaceae; g__Sphingomonas |
| cabff1f3d5d896228e792535ae412543 | d__Bacteria; p__Desulfobacterota_B; c__Binatia; o__UBA9968; f__UBA9968; g__; s__ | 0.722 | d__Bacteria |

SVs) that are common among random forest, adaptive boost, and gradient boost.**
taxonomic level of the predictions are indicated with a letter prefix where ‘d’, ‘p’, ‘c’,
espectively. The taxonomic classifications may not be up to the species level. The

| Confidence SILVA | RF Importance | ADA Importance | GBM Importance |
|---|---|---|---|
| 0.972 | 0.003 | 0.031 | 0.001 |
| 0.944 | 0.058 | 0.158 | 0.007 |
| 1.000 | 0.029 | 0.223 | 0.000 |
| 0.999 | 0.003 | 0.029 | 0.019 |
| 0.938 | 0.000 | 0.000 | 0.013 |

| | | | |
|---|---|---|---|
| 0.703 | 0.113 | 0.190 | 0.000 |
| 0.960 | 0.091 | 0.175 | 0.089 |
| 0.850 | 0.000 | 0.000 | 0.001 |
| 0.997 | 0.054 | 0.002 | 0.052 |
| 0.712 | 0.144 | 0.047 | 0.006 |

| | | | |
|---|---|---|---|
| 1.000 | 0.020 | 0.018 | 0.011 |
| 0.962 | 0.000 | 0.000 | 0.002 |
| 1.000 | 0.006 | 0.018 | 0.010 |
