## Supplementary Table 21 for "Grapevine Rootstock and Scion Genotypes’ Symbiosis with Soil Microbiome: A Machine Learning Revelation for Climate-Resilient Viticulture"

**Supplementary Table 21. Overlapping Features for country Classification. List of features (AS[illegible]**
taxonomy of the features was predicted with classifiers trained on GreenGenes2 and SILVA. The tabl[illegible]
with a letter prefix where 'd', 'p', 'c', 'o', 'f','g', and 's' are used for 'kingdom', 'phylum', 'class', 'o[illegible]
not be up to the species level. The confidence scores from the classifier are depicted in the adjacent co[illegible]

| ID | Taxon GG | Confidence GG | Taxon SILVA |
|---|---|---|---|
| 1062101 | d__Bacteria; p__Verrucomicrobiota; c__Verrucomicrobiae; o__Chthoniobacterales; f__UBA10450 | 1.000 | d__Bacteria; p__Verrucomicrobiota; c__Verrucomicrobiae; o__Chthoniobacterales; f__Chthoniobacteraceae; g__Candidatus_Udaeobacter |
| 107234 | d__Archaea; p__Thermoproteota; c__Nitrososphaeria_A; o__Nitrososphaerales; f__Nitrososphaeraceae; g__Nitrosocosmicus; s__Nitrosocosmicus | 0.743 | d__Archaea; p__Crenarchaeota; c__Nitrososphaeria; o__Nitrososphaerales; f__Nitrososphaeraceae |
| 144242 | d__Bacteria; p__Proteobacteria; c__Gammaproteobacteria; o__Burkholderiales_597441; f__SG8-41; g__UBA5216; s__UBA5216 sp902825845 | 0.898 | d__Bacteria; p__Proteobacteria; c__Gammaproteobacteria; o__Burkholderiales; f__TRA3-20; g__TRA3-20 |
| 1efc4ad517d4cd1fff23f0fd84c0eaa6 | d__Bacteria; p__Proteobacteria; c__Gammaproteobacteria; o__Burkholderiales_592522; f__Burkholderiaceae_A_592522 | 0.999 | d__Bacteria; p__Proteobacteria; c__Gammaproteobacteria; o__Burkholderiales; f__Comamonadaceae |
| 210475 | d__Bacteria; p__Proteobacteria; c__Gammaproteobacteria; o__Burkholderiales_597439; f__Usitatibacteraceae; g__; s__ | 0.910 | d__Bacteria; p__Proteobacteria; c__Gammaproteobacteria; o__Burkholderiales; f__Nitrosomonadaceae; g__IS-44 |

| | | | |
|---|---|---|---|
| 4c687e9abe11199b993e2ab668384d5c | d__Bacteria; p__Nitrospirota_A_437815; c__Nitrospiria; o__Nitrospirales; f__Nitrospiraceae; g__Nitrospira_C | 1.000 | d__Bacteria; p__Nitrospirota; c__Nitrospiria; o__Nitrospirales; f__Nitrospiraceae; g__Nitrospira; s__Nitrospira_japonica |
| 818123 | d__Bacteria; p__Acidobacteriota; c__Blastocatellia; o__UBA7656; f__UBA7656; g__QHVH01; s__QHVH01 sp003222245 | 0.982 | d__Bacteria; p__Acidobacteriota; c__Blastocatellia; o__11-24; f__11-24; g__11-24 |
| 859313 | d__Bacteria; p__Actinobacteriota; c__Acidimicrobiia_401430; o__Acidimicrobiales; f__Ilumatobacteraceae; g__Ilumatobacter_A; s__Ilumatobacter_A coccineus_A_400748 | 0.989 | d__Bacteria; p__Actinobacteriota; c__Acidimicrobiia; o__Microtrichales; f__Ilumatobacteraceae; g__Ilumatobacter |
| 9fa6663bd5e645f13343a7556ffa6dc9 | d__Bacteria; p__Planctomycetota; c__Planctomycetia; o__Gemmatales; f__Gemmataceae; g__; s__ | 0.891 | d__Bacteria; p__Planctomycetota; c__Planctomycetes; o__Gemmatales; f__Gemmataceae; g__uncultured |
| f02acd0bd6fb7a88ba739b06acba9e00 | d__Bacteria; p__Eisenbacteria; c__RBG-16-71-46; o__SZUA-252; f__SZUA-252 | 0.932 | d__Bacteria |

**/s) that are common among random forest, adaptive boost, and gradient boost.** The
e shows the predictions from both. The taxonomic level of the predictions are indicated
rder', 'family', 'genus', and 'species' respectively. The taxonomic classifications may
olumns.

| Confidence SILVA | RF Importance | ADA Importance | GBM Importance |
|---|---|---|---|
| 0.995 | 0.003 | 0.072 | 0.000 |
| 1.000 | 0.045 | 0.215 | 0.018 |
| 0.873 | 0.003 | 0.035 | 0.010 |
| 1.000 | 0.000 | 0.000 | 0.000 |
| 0.997 | 0.086 | 0.024 | 0.245 |

| | | | |
|---|---|---|---|
| 0.850 | 0.000 | 0.000 | 0.001 |
| 0.998 | 0.046 | 0.302 | 0.012 |
| 0.752 | 0.014 | 0.177 | 0.011 |
| 0.993 | 0.000 | 0.003 | 0.001 |
| 0.997 | 0.000 | 0.006 | 0.000 |
