## Supplementary Table 22 for "Grapevine Rootstock and Scion Genotypes’ Symbiosis with Soil Microbiome: A Machine Learning Revelation for Climate-Resilient Viticulture"

**Supplementary Table 22. Overlapping Features for Cultivar Classification. List of features (AS**
The taxonomy of the features was predicted with classifiers trained on GreenGenes2 and SILVA, resp
predictions are indicated with a letter prefix where ‘d’, ‘p’, ‘c’, ‘o’, ‘f’,’g’, and ‘s’ are used for ‘kingdo
taxonomic classifications may not be up to the species level. The confidence scores from the classifie

| ID | Taxon GG | Confidence GG | Taxon SILVA |
|---|---|---|---|
| 09a802948f82a388b8a9522a8887f3cd | d__Bacteria; p__Acidobacteriota; c__Blastocatellia; o__Pyrinomonadales; f__Pyrinomonadaceae_433871; g__PSRF01; s__ | 1.000 | d__Bacteria; p__Acidobacteriota; c__Blastocatellia; o__Pyrinomonadales; f__Pyrinomonadaceae; g__RB41 |
| 115ba67dce9f8b06ed78ddfe4af2926e | d__Bacteria; p__Gemmatimonadota; c__Gemmatimonadetes; o__Gemmatimonadales; f__GWC2-71-9 | 1.000 | d__Bacteria; p__Gemmatimonadota; c__Gemmatimonadetes; o__Gemmatimonadales; f__Gemmatimonadaceae; g__uncultured |
| 136760 | d__Bacteria; p__Nitrospirota_A_437815; c__Nitrospiria; o__Nitrospirales; f__Nitrospiraceae; g__Nitrospira_C | 1.000 | d__Bacteria; p__Nitrospirota; c__Nitrospiria; o__Nitrospirales; f__Nitrospiraceae; g__Nitrospira; s__Nitrospira_japonica |
| 1f01e277aeb09e117b913d10d8de6de5 | d__Bacteria; p__Proteobacteria; c__Gammaproteobacteria; o__Pseudomonadales_660879; f__Moraxellaceae; g__Acinetobacter | 1.000 | d__Bacteria; p__Proteobacteria; c__Gammaproteobacteria; o__Pseudomonadales; f__Moraxellaceae; g__Acinetobacter |
| 2e83ec04e6a19cea2b2e59ca62676d53 | d__Bacteria; p__Desulfobacterota_B; c__Binatia; o__Bin18; f__Bin18; g__JABFSC01; s__ | 0.999 | d__Bacteria; p__Entotheonellaeota; c__Entotheonellia; o__Entotheonellales; f__Entotheonellaceae; g__Entotheonellaceae |

| | | | |
|---|---|---|---|
| 36d2b09d4c04a114dfc2ab35017b0286 | d__Bacteria; p__Proteobacteria; c__Alphaproteobacteria; o__Sphingomonadales; f__Sphingomonadaceae; g__Novosphingobium_485351 | 0.969 | d__Bacteria; p__Proteobacteria; c__Alphaproteobacteria; o__Sphingomonadales; f__Sphingomonadaceae; g__Novosphingobium |
| 39b87a8034d3a49fbb5bbe7dab3a213b | d__Bacteria; p__Verrucomicrobiota; c__Verrucomicrobiae; o__Pedosphaerales; f__AV2; g__AV2; s__AV2 sp003218935 | 0.978 | d__Bacteria; p__Verrucomicrobiota; c__Verrucomicrobiae; o__Pedosphaerales; f__Pedosphaeraceae |
| 3a17438a7e43a08f8213f2f99e20cb5a | d__Bacteria; p__Acidobacteriota; c__Vicinamibacteria; o__Vicinamibacterales | 0.999 | d__Bacteria; p__Acidobacteriota; c__Vicinamibacteria; o__Vicinamibacterales; f__Vicinamibacteraceae |
| 4362620 | d__Archaea; p__Thermoproteota; c__Nitrososphaeria_A; o__Nitrososphaerales; f__Nitrososphaeraceae; g__Nitrosocosmicus; s__Nitrosocosmicus hydrocola | 0.872 | d__Archaea; p__Crenarchaeota; c__Nitrososphaeria; o__Nitrososphaerales; f__Nitrososphaeraceae; g__Candidatus_Nitrocosmicus |
| 4c687e9abe11199b993e2ab668384d5c | d__Bacteria; p__Nitrospirota_A_437815; c__Nitrospiria; o__Nitrospirales; f__Nitrospiraceae; g__Nitrospira_C | 1.000 | d__Bacteria; p__Nitrospirota; c__Nitrospiria; o__Nitrospirales; f__Nitrospiraceae; g__Nitrospira; s__Nitrospira_japonica |

| | | | |
|---|---|---|---|
| 6f65cbfe4878c5e3675028b742767b33 | d__Bacteria | 0.910 | d__Eukaryota |
| 7894541d84b621a7ffab4312fe3458ad | d__Bacteria; p__Acidobacteriota; c__Blastocatellia; o__Pyrinomonadales; f__Pyrinomonadaceae_433871; g__PSRF01; s__PSRF01 sp002427845 | 1.000 | d__Bacteria; p__Acidobacteriota; c__Blastocatellia; o__Pyrinomonadales; f__Pyrinomonadaceae; g__RB41 |
| 9eb3f6b5cb485d4d9fdb587b062818bb | d__Bacteria; p__Chloroflexota; c__Anaerolineae; o__Anaerolineales; f__EnvOPS12; g__UBA12294; s__UBA12294 sp002050275 | 0.999 | d__Bacteria; p__Chloroflexi; c__Anaerolineae; o__Anaerolineales; f__Anaerolineaceae; g__UTCFX1 |
| a4cccde07215f83a7eaf210b716bd92d | d__Bacteria; p__Proteobacteria; c__Alphaproteobacteria; o__Sphingomonadales; f__Sphingomonadaceae; g__Sphingomonas_L_486704 | 0.981 | d__Bacteria; p__Proteobacteria; c__Alphaproteobacteria; o__Sphingomonadales; f__Sphingomonadaceae; g__Sphingomonas |
| f98debb75a37c5b06ba598a461fbd1bf | d__Bacteria; p__Actinobacteriota; c__Rubrobacteria; o__Rubrobacterales; f__Rubrobacteraceae; g__; s__ | 0.940 | d__Bacteria; p__Actinobacteriota; c__Rubrobacteria; o__Rubrobacterales; f__Rubrobacteriaceae; g__Rubrobacter |

**Vs) that are common among random forest, adaptive boost, and gradient boost.**
ectively. The table shows the predictions from both. The taxonomic level of the
om', 'phylum', 'class', 'order', 'family', 'genus', and 'species' respectively. The
r are depicted in the adjacent columns.

| Confidence SILVA | RF Importance | ADA Importance | GBM Importance |
|---|---|---|---|
| 1.000 | 0.016 | 0.036 | 0.015 |
| 0.997 | 0.000 | 0.004 | 0.000 |
| 0.944 | 0.100 | 0.151 | 0.123 |
| 1.000 | 0.001 | 0.024 | 0.001 |
| 1.000 | 0.006 | 0.030 | 0.001 |

| | | | |
|---|---|---|---|
| 0.946 | 0.000 | 0.000 | 0.002 |
| 1.000 | 0.001 | 0.010 | 0.013 |
| 0.980 | 0.005 | 0.020 | 0.001 |
| 0.703 | 0.098 | 0.155 | 0.000 |
| 0.850 | 0.003 | 0.011 | 0.001 |

| | | | |
|---|---|---|---|
| 0.865 | 0.000 | 0.001 | 0.002 |
| 1.000 | 0.001 | 0.012 | 0.004 |
| 0.995 | 0.003 | 0.014 | 0.009 |
| 0.955 | 0.001 | 0.001 | 0.002 |
| 1.000 | 0.067 | 0.156 | 0.085 |
